# Genomic alterations enable *BRCA1* methylation loss and promoter bypass to drive resistance in high-grade serous ovarian cancer

**DOI:** 10.64898/2026.07.28.740856

**Authors:** Lijun Xu, Ksenija Nesic, Sally Beard, Jacinta Simmons, Xue Lu, Cassandra J. Vandenberg, Sharon M. Hoyte, Binny Jaradi, Ratana Lim, Franziska Geissler, Stacey L. Edwards, Joep Vissers, Anthony T. Papenfuss, Sean Grimmond, John V. Pearson, Clare L. Scott, Matthew J. Wakefield, Nicola Waddell, Olga Kondrashova

## Abstract

*BRCA1* promoter methylation predicts sensitivity to PARP inhibitors in high-grade serous ovarian cancer, yet therapeutic resistance is common and mechanistically unresolved. Using long-read direct DNA sequencing of patient-derived xenografts and cell lines, we resolved *BRCA1* methylation at single-molecule resolution with structural and transcriptomic analyses. We revealed two convergent PARP inhibitor and platinum resistance mechanisms, validated in patient tumors. First, focal, allele-specific loss of *BRCA1* methylation arose through local *cis*-acting genomic alterations, instead of global epigenetic reprogramming. Engineered *in cis* sequence alterations near the methylated *BRCA1* promoter were sufficient to induce methylation loss, restore homologous recombination, and confer resistance. Similar associations were observed across the genome, suggesting this mechanism extends beyond *BRCA1*. Second, *BRCA1* expression was restored despite intact promoter methylation via structural variant-mediated promoter bypass or alternative transcription initiation. Together, these findings redefine *BRCA1* methylation loss as a locus-restricted process and reveal multiple routes by which tumors escape PARP inhibitor therapy.

**Statement of Significance:** We show that high-grade serous ovarian cancers can restore *BRCA1* expression after therapy through multiple genomic mechanisms, including local methylation loss and promoter bypass, thereby re-establishing homologous recombination and driving PARP inhibitor resistance. These findings challenge reliance on *BRCA1* methylation alone as a predictive biomarker and support rational combination therapies for more durable responses.

## Introduction

High-grade serous ovarian cancer (HGSOC) is characterized by frequent defects in homologous recombination repair (HRR), a vulnerability that underlies sensitivity to platinum-based chemotherapy and PARP inhibitors (PARPi). Around 30% of HRR-deficient (HRD) tumors lack pathogenic variants in *BRCA1*, *BRCA2* or other core HRR genes, and instead harbor *BRCA1* promoter methylation (*meBRCA1*), resulting in transcriptional silencing (1). Tumors with homozygous, also described as complete, *meBRCA1* molecularly resemble *BRCA1*-mutant cancers and are typically sensitive to platinum and PARP inhibition (2–4). In contrast, tumors with heterozygous or partial *meBRCA1* retain residual *BRCA1* expression, restore HRR capacity, and exhibit therapeutic resistance (2). Despite their phenotypic similarities, *meBRCA1* tumors have a poorer prognosis compared to *BRCA1*-mutant cases (5,6). Importantly, most prior studies have not accounted for *meBRCA1* zygosity. Emerging clinical and genomic studies show that heterozygous *meBRCA1* arises from methylation loss under therapeutic pressure, representing an epigenetic mechanism of acquired resistance (7). Notably, *BRCA1* restoration confers resistance beyond PARPi to platinum chemotherapy (8), while BRCA1-proficient and -deficient states also differ in tumor metabolism (9) and immune detection (10–12), providing multiple avenues for the selection of resistant clones.

Despite its clear clinical relevance, the molecular basis of *meBRCA1* instability remains incompletely understood. Progress has been limited by the scarcity of representative pre-clinical models and by technical constraints of conventional methylation assays, which quantify aggregate CpG methylation, but do not resolve allelic configuration or adjacent sequence context. Targeted bisulfite sequencing has demonstrated epiallelic homogeneity within a small region of the *BRCA1* promoter (2,13), but the full extent of methylation loss across the locus, and its relationship to local sequence architecture and structural alterations, remains unknown. Moreover, bisulfite- and enzymatic-based whole-methylome approaches do not preserve native DNA sequence, limiting direct integration of methylation profiles with cis-acting genomic variants.

Here, we apply long-read direct DNA sequencing to resolve *meBRCA1* at a single-molecule and allelic resolution across a spectrum of pre-clinical HGSOC models, including patient-derived xenografts (PDX), patient-derived and engineered isogenic cell line sets. By integrating native DNA methylation profiling with transcriptomic, chromatin, and structural variant analyses, we delineate the epigenetic profile across the extended *BRCA1* region, identify cases of transcriptional restoration despite intact promoter methylation, and uncover local cis-acting genomic alterations associated with methylation loss. Our findings reveal that *meBRCA1* instability is highly locus-restricted, and *BRCA1* restoration is frequently driven by local genomic events rather than global epigenetic reprogramming. These insights, refine the mechanistic understanding of HRD reversibility and improve interpretation of methylation-based biomarkers in the context of therapeutic resistance.

## Results

### HRD scarring detected across samples with BRCA1 promoter methylation

To define the functional impact of *BRCA1* promoter methylation zygosity, we analyzed ten PDX (four homozygous *meBRCA1* and six heterozygous *meBRCA1*) and three cell lines (one homozygous, one heterozygous, and one *BRCA1*–mutant control) derived from chemo-naïve or platinum/PARPi-exposed patient tumors. Two cell lines also included isogenic engineered counterparts: UWB1 and *BRCA1*-complemented UWB1 (UWB1+BRCA1), and OVCAR8 with either complete *BRCA1* knockout or a single methylated allele (14)(Fig. 1A-B, Supplementary Table 1).

**Figure 1.**
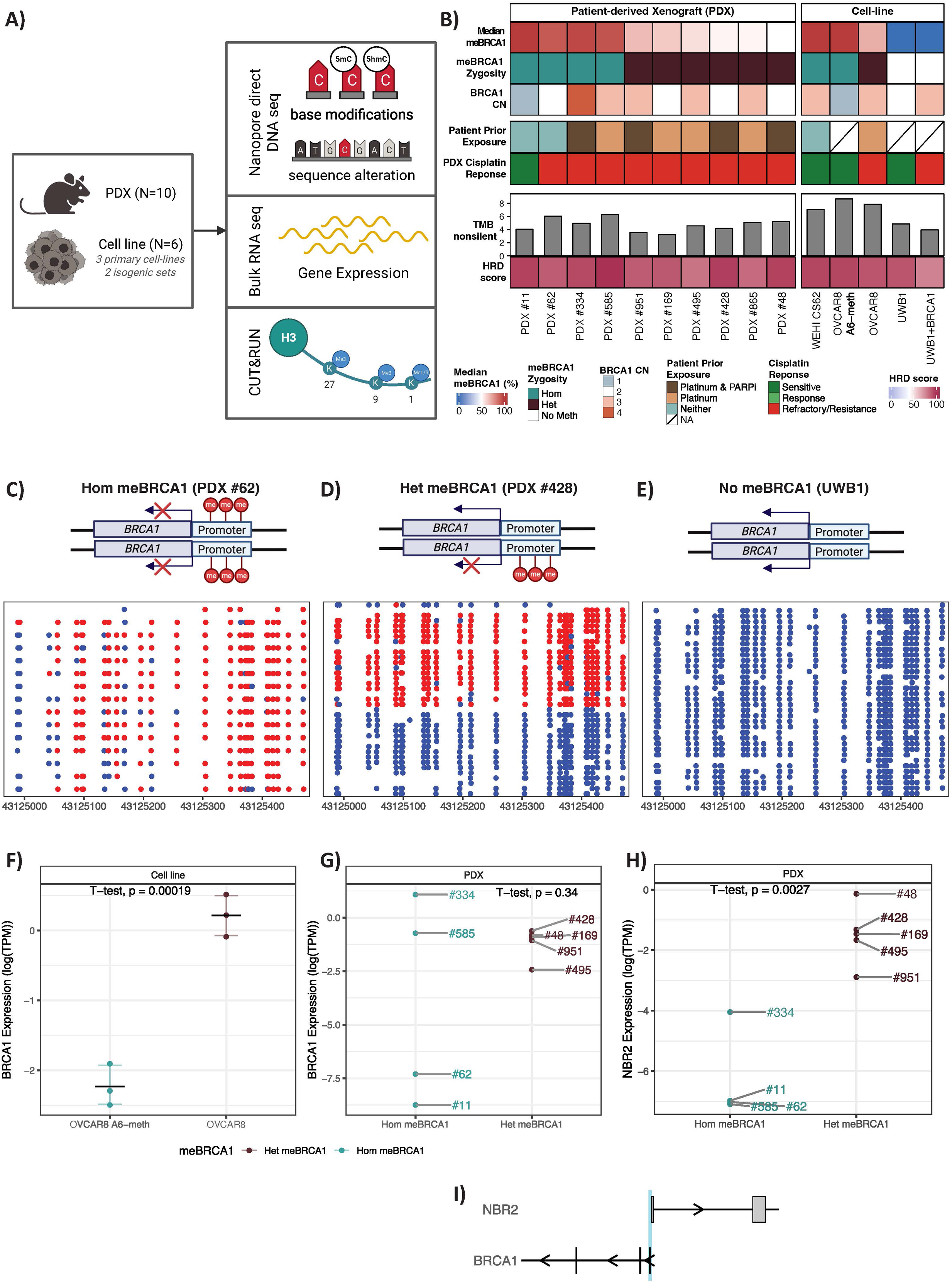
***BRCA1* promoter methylation zygosity concordance with treatment history and therapeutic response but does not uniformly dictate *BRCA1* expression.** (**A**) Overview of the experimental models and multi-omic profiling strategy. Patient-derived xenograft (PDX) models and cell-lines were analysed using long-read Nanopore sequencing, bulk RNA sequencing and CUT&RUN profiling. **(B)** Integrated molecular and phenotypic characterization of PDX and cell-line models. Heatmap summarizes *BRCA1* promoter methylation status, promoter methylation zygosity, copy number status, patient treatment exposure prior to model generation, PDX cisplatin response, tumour mutational burden (TMB, mutation per Mb), and homologous recombination deficiency (HRD) score. Models are grouped by *BRCA1* promoter methylation zygosity. **(C–E)** Examples of allele-resolved CpG methylation profiles across the BRCA1 promoter determined by Nanopore direct DNA sequencing for homozygous *meBRCA1* PDX #62 **(C)**, heterozygous *meBRCA1* PDX #428 **(D)** and BRCA1-unmethylated cell line UWB1 **(E)**. Each line represents an individual sequenced DNA molecule, while each dot represents a CpG site, coloured by methylation state (methylated, red; unmethylated, blue). Schematic illustrations above each panel depict the corresponding epiallelic configuration. (F) *BRCA1* mRNA expression levels in OVCAR8 isogenic cell line pair stratified by *BRCA1* promoter methylation zygosity. Expression per cell-line was profiled in triplicate and represented using log2-transformed transcripts per million (TPM), with horizontal line denoting median expression and error bars marking standard deviation. (G) *BRCA1* expression across PDX models. Despite complete promoter methylation, PDX #585 and PDX #334 retain *BRCA1* expression at levels comparable to heterozygous *meBRCA1* models. **(H)** Expression of *NBR2* gene, sharing the bidirectional *BRCA1* promoter, across PDX models. *NBR2* expression remains reduced in homozygous *meBRCA1* models, confirming intact promoter-level repression despite retained *BRCA1* expression in selected cases. **(I)** Schematic representing shared bidirectional promoter, marked in blue, between *BRCA1* and *NBR2* genes.

All models exhibited high HRD scores consistent with genomic scarring (Fig. 1B; Supplementary Fig. 1). The only exception was model #585, which had a high HRD score but a low HRDetect probability score, due to lack of detected SBS3 mutational and RS3 structural variant signatures (Supplementary Fig. 1). No pathogenic alterations or promoter methylation events were detected in other core HRR genes across *meBRCA1* models (Supplementary Fig. 2), indicating that *BRCA1* promoter methylation was the primary driver of HRD.

Notably, the patient-derived homozygous *meBRCA1* models originated from two treatment–naïve primary tumors and two pre-treated tumors, whereas all heterozygous models were derived from treatment-exposed tumors (Fig. 1B), supporting methylation loss under therapeutic pressure. Among treatment-naïve pre-clinical models, one was sensitive to cisplatin (#11), while the other was resistant (#62). For the pre-treated tumors, none were cisplatin-sensitive, two were resistant (#48, #585), while six were refractory (#334, #495, #951, #169, #428, #865) (Supplementary Table 2, Supplementary Fig. 3). For cell lines, the homozygous *meBRCA1* models (OVCAR8 A6–meth and WEHI CS62) were cisplatin sensitive, while the heterozygous OVCAR8 was resistant (Supplementary Fig. 3).

Using long–read Nanopore direct DNA sequencing, we resolved the allelic configuration of the *BRCA1* promoter across models. Since the human component in PDX material is made up of pure tumor cells, it enabled resolution of methylation and genomic alterations without contaminating non-tumor human cells. All *meBRCA1* models showed loss of heterozygosity (LOH) across chr17 regardless of methylation zygosity (Supplementary Fig. 4), consistent with heterozygous *BRCA1* methylation being followed by LOH as an early driver event (aligning with Knudson’s ‘2-hit’ hypothesis), with subsequent methylation loss driving treatment resistance. Across all models, methylation displayed a homogeneous epiallelic pattern, where individual alleles were either fully methylated or fully unmethylated across the CpG island (Fig. 1C-E; Supplementary Fig. 5). Homozygous models showed complete methylation (Fig. 1C), heterozygous models exhibited at least one unmethylated allele (Fig. 1D), and controls were unmethylated (Fig. 1E). This pattern enabled clear classification of methylation zygosity.

As expected, all heterozygous models expressed *BRCA1* mRNA (Fig. 1F–G), reflecting transcription from the unmethylated allele(s). Notably, two homozygous PDX models established from treatment-exposed patient tumors (#334: post-platinum and PARPi; #585: post-platinum), also exhibited *BRCA1* expression at levels similar to heterozygous models, despite complete promoter methylation (Fig. 1G). In contrast, *NBR2*, a gene sharing the bidirectional *BRCA1* promoter, remained suppressed in all homozygous *meBRCA1* models (Fig. 1H-I), confirming intact promoter-level repression. These findings indicate that *BRCA1* can escape canonical promoter silencing through gene–specific mechanisms.

### Restored BRCA1 expression and PARPi resistance despite intact homozygous promoter methylation

To investigate the mechanism of escape from promoter–mediated silencing in the two homozygous *meBRCA1* PDX models with restored *BRCA1* expression (#334 and #585), we performed transcript–level analyses. PDX #334 expressed a full–length *BRCA1* transcript originating from the canonical promoter (Fig. 2A). In contrast, in PDX #585, we identified a novel isoform of *BRCA1* transcript initiating at exon 9 rather than the canonical exon 1 start site (Fig. 2B). This model was also distinct in exhibiting a low HRDetect probability score described in the previous section, unlike other *meBRCA1* models (Supplementary Fig. 1). PDX #334 showed BRCA1 protein expression at levels consistent with the wild-type *BRCA1* model, whereas #585 showed negligible staining using an antibody against a C-terminal epitope retained in the predicted isoform, possibly reflecting inefficient translation initiation or instability of the shortened protein (Fig. 2C). Despite this, both PDX #334 and #585 exhibited *in vivo* resistance to both generations of PARP inhibitors (rucaparib – first generation, or saruparib – next generation), consistent with restored HR activity (Fig. 2D–E).

**Figure 2.**
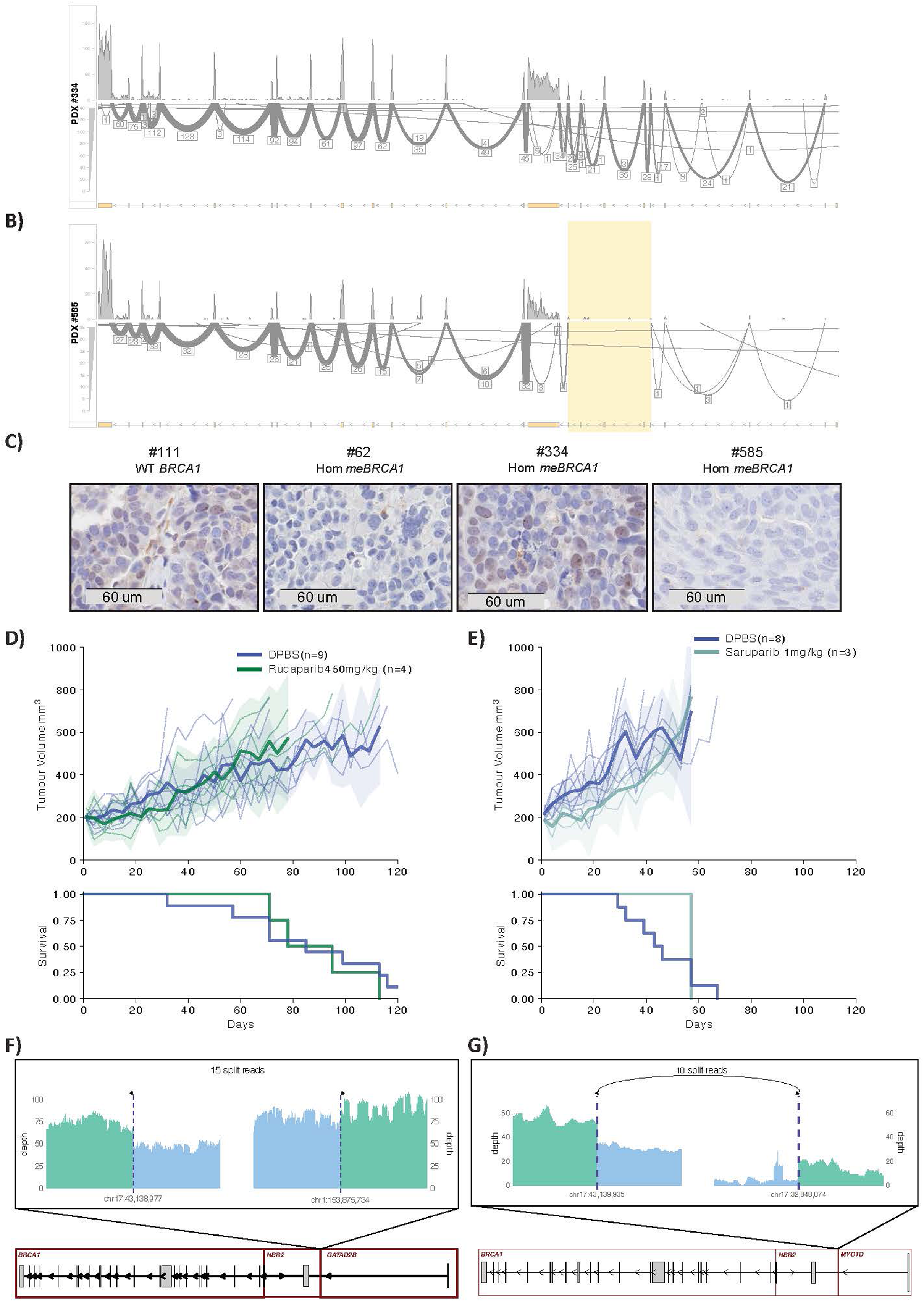
Restored *BRCA1* expression and PARPi resistance despite intact homozygous promoter methylation. (**A–B**) Sashimi plots showing transcript-level *BRCA1* expression in the two homozygous *meBRCA1* PDX models with restored *BRCA1* expression: PDX #334 showing expression of a full-length *BRCA1* transcript initiating from the canonical exon 1 promoter **(A)**; PDX #585 showing a non-canonical *BRCA1* transcript initiating downstream of the canonical promoter, with transcription beginning at exon 9 **(B)**. Region corresponding to alternative transcription initiation in PDX #585 is highlighted. **(C)** BRCA1 protein expression detected by immunohistochemistry. A representative image is shown for 4 tested independent PDX mouse samples. **(D–E)** Tumour growth curves and corresponding Kaplan-Meier survival plots showing *in vivo* PARP inhibitor responses for PDX #334 **(D)** and PDX #585 **(E)**. **(F–G)** Schematic representation of structural rearrangements at the *BRCA1* locus associated with promoter-independent *BRCA1* expression. Read-depth coverage (green) and the reciprocal breakpoint locus (blue) are shown over a 5 kb window; dashed lines indicate breakpoint positions and bracket labels denote supporting split-read counts. **(F)** In PDX #334, rearrangements reposition the *BRCA1* coding region under the influence of the *GATAD2B* promoter. **(G)** In PDX #585, rearrangements reposition the *BRCA1* coding region under the influence of the *MYO1D* promoter. Gene models for *BRCA1* (ENST00000357654.9), *NBR2* (ENST00000657841.1), *GATAD2B* (ENST00000368655.5) and *MYO1D* (ENST00000318217.10) are shown with rearranged segments highlighted.

Structural variant (SV) analysis revealed the likely mechanisms enabling this promoter-independent *BRCA1* expression. In #334, we identified rearrangements hijacking the *GATAD2B* promoter to drive ectopic *BRCA1* transcription (Fig. 2F). This rearrangement was also detected in the patient tumor sample that was used to generate this PDX model. In #585, we detected a more complex cluster of SVs upstream of *BRCA1* not resolvable with whole-genome sequencing (Supplementary Fig. 6). Using adaptive sequencing to enrich for coverage depth around *BRCA1*, we identified a candidate translocation between *NBR2* and *MYO1D* that is likely linked to restored expression (Fig. 2G). These findings implicate genomic reconfiguration as a shared mechanism enabling alternative transcription initiation. Importantly, these events occurred without loss of *BRCA1* promoter methylation, indicating that *BRCA1* reactivation arose through sequence alterations that bypass the silenced promoter.

### BNC1 methylation loss co-occurred with meBRCA1 loss

Reversion of *meBRCA1* silencing may also occur due to global changes in epigenetic silencing and methylation maintenance throughout the genome. To test this, we assessed genome–wide DNA methylation across models. We did not see consistent global methylation distribution correlated with *meBRCA1* zygosity (Supplementary Fig. 7A). Instead, methylation patterns were highly heterogeneous, with the greatest variability arising within partially methylated domains (PMDs) (median methylation: 31%–92%; Supplementary Fig. 7B). Additionally, differential methylation analysis revealed no significant methylation differences between homozygous and heterozygous *meBRCA1* models, including at the *BRCA1* locus, indicating that global epigenomic remodeling was not associated with *BRCA1* methylation loss in our models.

Given the absence of global methylation changes, and the fact that differential analysis did not detect homozygous-to-heterozygous changes at the *BRCA1* locus, we explored whether methylation loss was driven by locus-specific rather than genome-wide processes and refined our strategy for a more focused screen of promoter CpG islands that may co–evolve with *BRCA1* methylation loss. Screening promoter CpG islands for coordinated methylation and transcriptional changes identified five genes, *BNC1*, *HS3ST2*, *KATNAL2*, *MROH6*, and *SYCP2*, with consistent promoter methylation and transcriptional changes across PDX models (Supplementary Fig. 8). In each case, homozygous *meBRCA1* HRD PDX models showed higher promoter methylation and lower gene expression than the heterozygous *meBRCA1* models.

To validate these findings, we analyzed an independent ICGC dataset of 75 primary and 20 matched relapsed HGSOC tumors with WGS and 450K methylation data. Three primary– relapse pairs showed reduced (AOCS-091, AOCS-094) or unchanged (AOCS-093) *BRCA1* promoter methylation with high *BRCA1* expression at relapse (Supplementary Fig. 9A-B). Among candidate loci, *BNC1* was the only gene showing consistent promoter methylation loss between primary and relapse, consistent with *BRCA1* methylation loss during tumor evolution under therapeutic pressure (Supplementary Fig. 9C).

Across the ICGC HGSOC cohort, *BNC1* promoter methylation was higher in *meBRCA1* tumors; however, it was also present in many *BRCA1* unmethylated, chemo-naïve cases (Supplementary Fig. 9D). These findings suggest that while *BNC1* promoter methylation loss frequently co–occurs with *BRCA1* methylation loss, the initial events causing *BNC1* promoter methylation are more likely independent events that may respond to shared selective forces, while loss could be associated with treatment pressures or disease progression.

### BRCA1 methylation instability is locus-restricted

Since genome–wide methylation patterns did not explain the transition from homozygous to heterozygous *BRCA1* promoter methylation, we hypothesized that specific local events at the DNA or epigenetic level were responsible for methylation loss. To investigate the role of the local DNA changes, we next examined the *BRCA1* locus at high resolution, extending beyond the ∼250 bp region covered by methylation arrays (15).

Across all heterozygous *meBRCA1* models, methylation loss was highly focal, confined to the region between the *BRCA1* promoter and extending towards but not into the adjacent CpG island within the *NBR2* intron (Fig. 3A; Supplementary Fig. 10). The consistent boundary of demethylation supports a model in which *BRCA1* methylation loss arises from localized, rather than global, epigenetic processes. Additionally, given that LOH at the *BRCA1* locus is an early event (detected in all *meBRCA1* models), we hypothesized that focal promoter methylation loss in heterozygous models is influenced by somatic, allele-specific changes acquired after *BRCA1* LOH.

**Figure 3.**
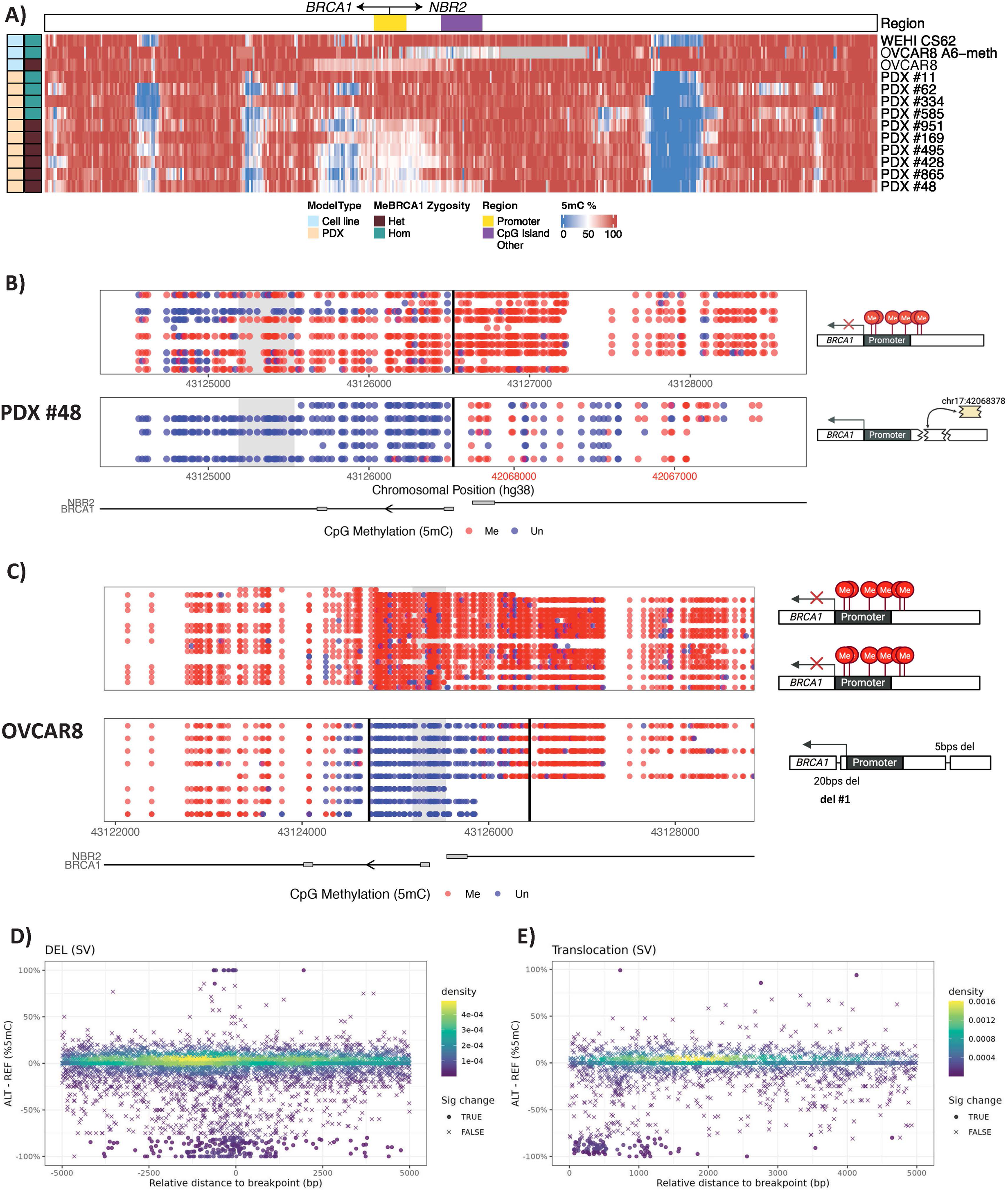
*BRCA1* promoter methylation instability is locus-restricted and associated with in cis genomic alterations. **(A)** High-resolution CpG methylation profiles across the extended *BRCA1* locus in patient-derived xenograft (PDX) and cell-line models. Methylation loss in heterozygous *meBRCA1* models is focal and confined to the region spanning the *BRCA1* promoter, but not the adjacent CpG island within the *NBR2* intron. Heatmap colours indicate percentage CpG methylation (unmethylated blue, methylated red). The genomic location is relative as heatmap shows all CpGs without adjustment for genomic distance. OVCAR8 A6-meth has a grey area because of the genetically engineered deletions proximal to *BRCA1*. **(B)** Allele-resolved CpG methylation profiles of PDX #48 showing a ∼10 kb translocation proximal to the *BRCA1* promoter that occurs *in cis* with the unmethylated allele. CpG methylation states are shown for reads aligned to each allele, with the translocation breakpoint indicated. **(C)** Allele-resolved CpG methylation profiles of OVCAR8 (*me*BRCA1/*me*BRCA1/wt) identifying two heterozygous deletions, one within the *BRCA1* promoter (del #1) and one within the *NBR2* intron. Both deletions are *in cis* to the unmethylated allele and were detected in the heterozygous *meBRCA1* parental OVCAR8 line, but absent in the homozygous *meBRCA1* OVCAR8 A6-meth derivative. Schematic diagrams indicate deletion positions. **(D-E)** Density plots showing CpG methylation changes relative to the nearest breakpoint for **(D)** large scale deletions (DEL SV ≥50 bp) and **(E)** translocations (BND SV). Each point represents an individual CpG site. The x-axis indicates the distance from the closest breakpoint (negative values, upstream; positive values, downstream), and the y-axis shows the methylation difference on the altered allele relative to the reference allele. CpGs exhibiting significant methylation change (Fisher’s exact test, *P* ≤ 0.05 and absolute methylation difference ≥80%) are marked with crosses, while all other CpGs are shown as circles. Color reflects CpG density, ranging from low (yellow) to high (navy).

We assessed single nucleotide variants (SNVs), indels, and SVs across the region. Seventeen recurrent homozygous SNVs, corresponding to known germline haplotypes, were identified in four of seven heterozygous models (Supplementary Fig. 11A). Their high population frequencies (∼0.3–0.5), homozygosity, and presence in samples lacking *BRCA1* methylation do not support a role in *meBRCA1* loss. Recurrent insertions co-segregated with these SNVs (Supplementary Fig. 11B), and their homozygous state suggests that these events predate LOH at *BRCA1*.

In contrast, two likely somatic variants exhibited clear spatial and allelic concordance with the unmethylated allele. PDX #48 carried a 10 kb translocation positioned proximal to the *BRCA1* promoter, occurring *in cis* with the unmethylated allele (Fig. 3B). This translocation was also confirmed in the patient tumor sample from which the PDX was derived (Supplementary Fig 12). In the OVCAR8 (*BRCA1*^me/me/wt^) cell line, two heterozygous deletions in the *BRCA1* promoter and *NBR2* intron (with 5’ deletion referred to as del #1, and 3’ as del #2) were also *in cis* with the unmethylated allele (Fig. 3C; Supplementary Fig. 11C). Supporting the unmethylated allele-specific data, both deletions were detected exclusively in the heterozygous *meBRCA1* parental OVCAR8 line and were absent from the fully methylated, *BRCA1*–silenced A6–meth derivative (*BRCA1*^me/-/-^).

Due to the locus-restricted pattern of *BRCA1* promoter methylation loss, we next considered whether histone modifications might contribute to its regulation. However, apart from a weak H3K9me3 signal that was below the peak calling threshold and is typically associated with DNA methylation maintenance (16), we observed no enrichment of the canonical repressive histone mark H3K27me3 at the promoter (Supplementary Fig. 13). Instead, prominent H3K4me3 peaks, indicative of active transcription, were detected in OVCAR8 and UWB1 cells, both of which expressed *BRCA1* (Supplementary Fig. 14). These results suggest that common repressive histone modifications, H3K9me3 and H3K27me3, do not play a significant role in maintaining *BRCA1* promoter methylation in HGSOC.

### Genome-wide association between somatic sequence alterations and in cis methylation patterns

Our locus–level analyses at *BRCA1* showed that promoter methylation loss was confined to a small region and linked to *in cis* sequence alterations. This raised the question of whether such sequence-methylation association is unique to *BRCA1*, or whether it reflects a broader genomic phenomenon. We therefore proceeded to investigate whether somatic sequence alterations across the genome similarly associate with focal methylation perturbations, whether this extends beyond deletions and translocations to other variant classes, and whether specific genomic features influence methylation stability following sequence disruption.

We examined two engineered cell line sets (OVCAR8 parental vs A6-meth and *BRCA1* KO; and UWB1±*BRCA1*) and leveraged heterozygous ‘somatic’ variants that arose in ‘daughter’ lines, but were absent from their ‘parental’ counterparts, enabling assessment of methylation changes linked to naturally occurring sequence disruptions (Supplementary Fig. 15). Since the base–calling error rate of long-read sequencing limits reliable SNV detection, we restricted our analysis to insertions, deletions and SVs, with multi–base evidence that support robust allele–specific methylation profiling.

Across assessed cell line sets, somatic alterations frequently coincided with focal methylation changes at nearby CpGs, rather than stochastic variation (Fig. 3D-E; Supplementary Fig. 16-18). Deletions showed the strongest and most frequent association. Both small indels (<50bp, INDEL) and larger structural deletions (≥50bp, SV) were significantly enriched for methylation loss at CpGs located near breakpoints (permutation test, 100,000 resamples, p ≤ 1 × 10⁻⁵; Fig. 3E and Supplementary Fig. 16A). In OVCAR8 derivatives, 15.6% (77/495) of small deletions and 56.9% (37/65) of large deletions were associated with methylation loss within 5 kb of the breakpoint. Methylation gains were less common, only 6.6% (21/314) of small deletions and 2.4% (1/41) of large deletions had co-occurring methylation gain (Supplementary Fig. 16B). Notably, within smaller deletions the likelihood of methylation changes increased with deletion length (median length for methylation changed is 7bp, methylation unchanged is 1bp; Supplementary Fig. 16C).

Insertions and duplications were also associated with local methylation perturbations (Fig. 3D and Supplementary Fig. 17-18), although at lower frequency (Supplementary Fig. 17C, 18B) and with no size association compared with deletions (Supplementary Fig. 17D-E, 18C). Translocations were less frequent overall but were commonly associated with proximal methylation change – 33% (7/21) of events induced methylation loss and 10% (2/21) induced methylation gain (Fig. 3E and Supplementary Fig. 18D).

Genomic feature analysis showed that repetitive elements, particularly LINE and DNA repeats, were significantly enriched at breakpoints associated with methylation alteration, whereas CpG island architecture, gene context, and long noncoding RNA (lncRNA) overlap did not show consistent associations (Supplementary Table 3).

We also noted sample–dependent variability, where OVCAR8 derivatives (HRD) exhibited more methylation changes than UWB1+*BRCA1* (HR-proficient) (Supplementary Fig. 16B, 17C-D and 18B,D). To test whether HR function contributed, we compared microhomology patterns between alterations associated with methylation change versus those without. We found no differences in microhomology usage (Supplementary Fig. 19A–B) and no enrichment of methylation–altering events among deletions repaired via non–microhomology mechanisms or theta-mediated end-joining, defined as deletions > 4 bp with ≥ 2 bp of microhomology (17) (Supplementary Fig. 19C).

### Cell-line validation of sequence disruption on meBRCA1 loss

To directly test whether local sequence disruption can drive *BRCA1* promoter methylation loss, we introduced CRISPR–guided deletions around del #1 into the single, fully methylated *BRCA1* allele of OVCAR8 A6–meth cells (*BRCA1*^me/-/-^) (Fig. 4A). Targeted bisulfite sequencing demonstrated reproducible reductions in *BRCA1* promoter methylation with two independent sgRNAs (parental: 95%; guide #1: 76%; guide #2: 84%; Fig. 4B-C).

**Figure 4.**
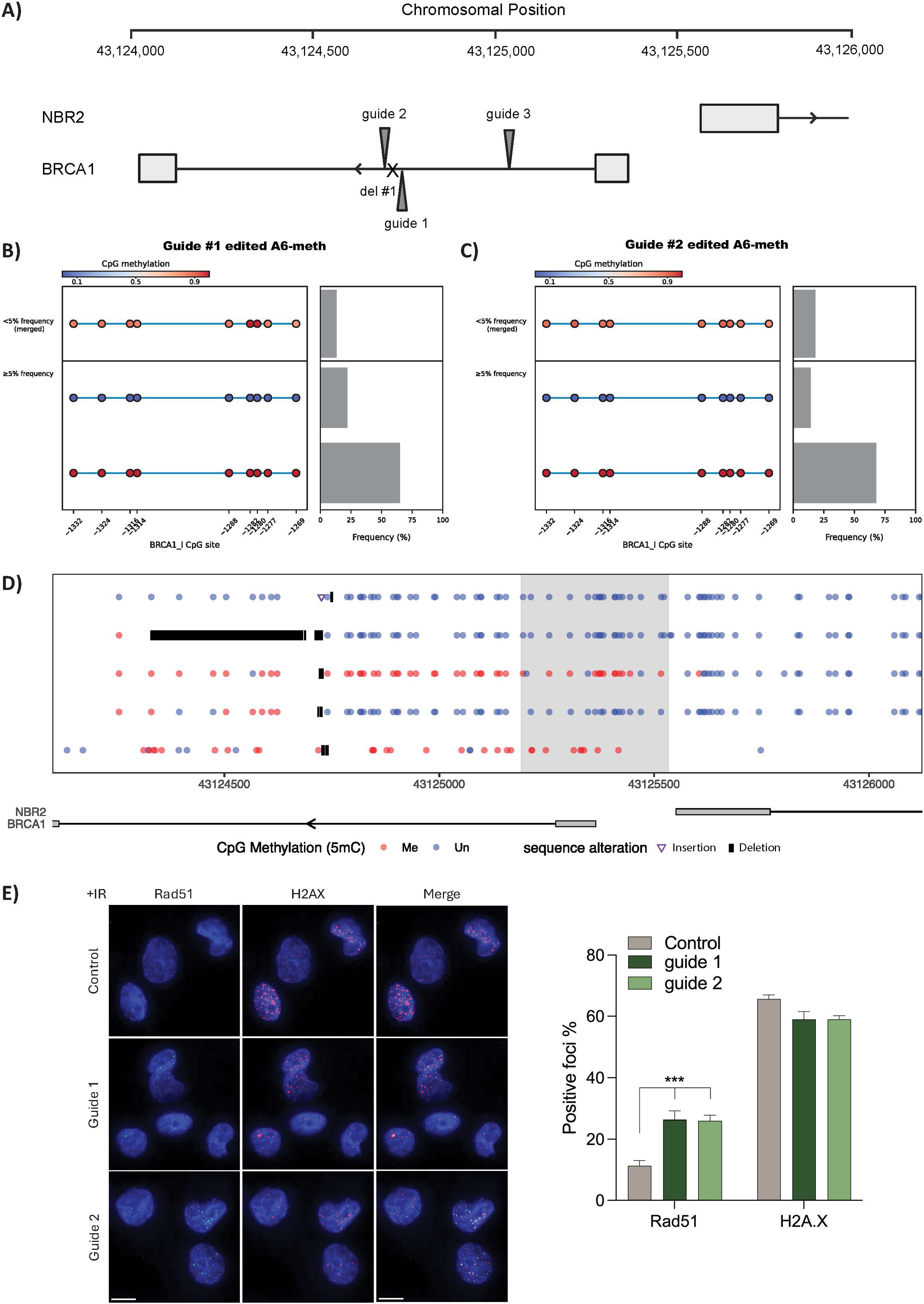
CRISPR-mediated deletion perturbs *BRCA1* methylation and restores homologous recombination. **(A)** Schematic of the *BRCA1/NBR2* locus showing positions of three guide RNAs targeting intron 1. Guides 1 and 2 flank the intended deletion (del #1); guide 3 targets a downstream site. **(B–C)** Single-molecule *BRCA1* methylation profiles in OVCAR8 A6-meth cells edited with **(B)** guide 1 or **(C)** guide 2. Each row represents an epiallele, with CpG methylation indicated from low (blue) to high (red) relative to the *BRCA1* transcription start site. **(D)** CpG methylation following CRISPR-induced deletion (del #1) in A6-meth cells. Black bars indicate deletion boundaries; edited clones show disrupted promoter methylation. **(E)** RAD51 immunofluorescence in edited cells. Representative images show RAD51 foci (magenta) and nuclei (DAPI, blue), with quantification of RAD51-positive cells at right, indicating restored homologous recombination.

Using long–read direct DNA sequencing with adaptive sampling to enrich for *BRCA1* promoter reads, we found that editing at chr17:43124711 (guide #1) produced heterogeneous sequence alterations, including both deletions and insertions. Despite this variability, four out of five edited clones exhibited perturbation of promoter methylation (Fig. 4D), while the remaining one had methylation changes downstream of the promoter. A similar reduction in methylation was observed using an independent sgRNA (guide #3; 322 bp upstream of the del #1; methylation of *BRCA1* promoter: 92%; Supplementary Fig. 20A-B). No promoter demethylation was detected in reads without edits or with deletions smaller than 5 bp (Supplementary Fig. 20C).

Following 10-days of selection with the PARP inhibitor, olaparib, guide #1- and guide #2-edited clones, with parental A6–meth control cells, showed selective enrichment of the unmethylated allele (parental: 5% to average 8%; guide #1: 24% to average 100%; guide #2: 16% to average 97%, Supplementary Fig. 21), indicating that subclonal *BRCA1* methylation loss is enriched under treatment pressure. Consistent with this, pre-selected edited clones exhibited increased resistance to both olaparib and cisplatin compared to unedited controls (Supplementary Fig. 22), and showed enhanced RAD51 foci formation, confirming restoration of HRR through *BRCA1* re-expression (Fig. 4E).

## Discussion

*BRCA1* promoter methylation is a prominent and well-established driver of HRD and therapeutic sensitivity in HGSOC (2,18). Here, we investigated the genomic and epigenomic determinants of *BRCA1* methylation loss to understand whether resistance arises through global epigenetic remodeling or locus-specific mechanisms. Using long-read direct DNA sequencing together with transcriptomic profiling, we demonstrate that *BRCA1* restoration and resistance can emerge through two distinct but convergent mechanisms – focal methylation loss and promoter bypass.

Across all models, *BRCA1* promoter methylation was largely homogeneous, and LOH was detected regardless of methylation zygosity, reinforcing the model that *BRCA1* promoter methylation is acquired early and followed by LOH during tumor establishment, leading to HRD. All models displayed HRD-associated genomic scars, consistent with *BRCA1* mutant tumors. However, homozygous promoter methylation alone did not universally dictate *BRCA1* transcriptional silencing. Instead, we demonstrated promoter bypass as a resistance mechanism in pre-treated HGSOC, where structural variants hijacked neighboring active regulatory elements to restore *BRCA1* expression, including one case expressing a truncated *BRCA1* isoform. Similar structural-variant-driven promoter hijacking has been described in treatment-resistant *BRCA1*-methylated breast cancer (3,19), supporting this as a convergent route to resistance across tumor types. These findings also have important clinical implications, indicating that *BRCA1* promoter methylation alone is insufficient as a biomarker for HRD and PARPi sensitivity in treatment-exposed tumors. Functional measures of HRD, such as RAD51 foci formation, together with emerging immune and circulating biomarkers (12) are likely to provide more accurate assessment of therapeutic response.

Focal promoter methylation loss represents a second, mechanistically distinct route to resistance, in some cases associated with *in cis* genomic alterations acquired after LOH, likely under treatment pressure. Unlike promoter bypass, methylation loss was localized to the *BRCA1* promoter (∼1 kb) without evidence of global epigenetic remodeling, arguing against genome-wide loss of methylation maintenance. Isogenic analyses further showed this genomic alteration-methylation association extends beyond *BRCA1*, though not uniformly across all genomic alterations. Beyond a modest enrichment at repetitive elements, we were unable to identify genomic features that consistently predicted methylation loss.

Mechanistically, our findings support a model in which DNA repair processes drive stochastic methylation loss, potentially through incomplete methylation maintenance by DNMT1 during DNA repair (20). HRD cells rely largely on error-prone non-homologous and theta-mediated end-joining (TMEJ) DNA repair. Although we found no enrichment of microhomology to suggest a specific role for TMEJ, it remains unclear whether different DNA repair pathways vary in their ability to recruit DNMT1 and therefore differ in their propensity to lose methylation. Additionally, methylation loss may in part be a consequence of the break-and-repair process itself rather than repair errors, as several heterozygous *meBRCA1* cases lacked detectable local genomic alterations.

Given that platinum and PARP inhibitors themselves induce DNA breaks and could promote methylation loss as an unintended consequence, the relative contribution of different repair pathways and therapies to this phenomenon warrants further investigation.

Together, our findings define a dual model of *BRCA1* restoration and resistance in *BRCA1*-methylated HGSOC, comprising focal promoter methylation loss linked to DNA repair and promoter bypass driven by structural rearrangements. Notably, *BRCA1* restoration was detected in every pre-treated model in our cohort, marking it as a recurrent route to acquired resistance. Compared with *BRCA1/2* reversions, which require specific coding sequence changes, promoter demethylation can arise from a broad range of local genomic events that occur at high frequencies in HRD cells. This study highlights the limitations of platinum chemotherapy and single-agent PARP inhibitor strategies in this population and underscores the urgent need for combination approaches guided by functional measures of HRR or alternative treatment biomarkers rather than static methylation markers, with longitudinal monitoring to capture restoration as it emerges under treatment.

## Methods

### Study Cohort and Ethics

Tumor samples were collected from patients enrolled in the WEHI-Stafford Fox Rare Cancer Program under a protocol approved by the Melbourne Health Human Research Ethics Committee (HREC/15/MH/396; Royal Melbourne Hospital Project 2015.300), with governance review by the WEHI HREC (Project number G16/02) and QIMR Berghofer HREC (P3456, P2095, P3871). All patients provided written informed consent to participate in the study. The study was conducted in compliance with the NHMRC National Statement on Ethical Conduct in Human Research 2007 (updated 2018). Animal experiments were conducted following the Australian Code for the Care and Use of Animals for Scientific Purposes 8th Edition, 2013 (updated 2021) and with approval by the WEHI animal ethics committee (Projects 2019.024, 2022.030 and 2022.038).

### Generation of PDX models and in vivo treatments

Fresh tumor samples were collected from surgery, biopsy, ascites or pleural tap from patients enrolled in the WEHI-Stafford Fox Rare Cancer Program. To generate patient-derived xenografts (PDX) from surgical or biopsy samples, tumor fragments 10 to 20 mm^3^ were immediately implanted subcutaneously into the flank of NOD.Cg-*Prkdc*^scid^ *Il2rg*^tm1Wjl^/SzJ (NSG) mice. For ascites or pleural fluid samples, tumor cell enrichment (immune cell depletion) was performed using the Tumor Cell Isolation Kit, human (Miltenyi Biotech, 130-108-339). The isolated tumor cells in DPBS were mixed 50:50 with Matrigel and 100 µL injected subcutaneously into the flank of NSG mice. Mice were monitored weekly for tumor growth and harvested when they approached the humane endpoint of 1000 mm^3^. Successfully established PDX models were perpetuated by serial transplantation. PDX models were verified histologically with pathologist review and DNA sequencing comparison with the patient tumor. PDX models #11, #48, #62 and #169 have been published previously (21).

Animals were housed in the WEHI Bioservices facility in Tecniplast IVC caging with food and water ad libitum; maintained at approximately 21°C (+/-3°C) and humidity 40-70%. The facility has a light dark cycle of 12h/12h, with red colored lighting for use during the dark cycle if required.

For treatment cohorts, tumors were measured twice weekly with digital calipers and recorded using StudyLog Desktop software (StudyLog Systems Inc, CA, USA). Mice were randomized to treatment at tumor volume 180-300 mm^3^: cisplatin (Pfizer) 4 mg/kg was administered by intraperitoneal injection (20 µL/g) on days 1, 8 and 18; AZD5305 (saruparib) 1 mg/kg in water/HCl pH 3.5–4 and rucaparib 450 mg/kg in 0.5% methylcellulose were administered orally (10 µL/g) 5 days on/ 2 days off for 4 weeks. Experimental endpoints were tumor volume 700-1000 mm^3^ or 120 days following randomization. SurvivalVolume was used to graph tumor volume and Kaplan-Meier curves and for statistical analysis (22).

### Cell culture

WEHI-CS62, OVCAR8, OVCAR8 A6-meth and OVCAR8 H4-KO, were cultured in DMEM/F-12 GlutaMAX media (Thermo Fisher, 10565018) supplemented with 10% (v/v) foetal bovine serum (FBS) (Thermo Fisher, 10099141), 5 μg/ml insulin (Sigma-Aldrich, 11376497001), 50 ng/ml EGF (Sigma-Aldrich, E9644-.2MG), 1 μg/ml hydrocortisone (Sigma-Aldrich, H0396-100MG) and 100 U/ml penicillin and streptomycin (Thermo Fisher, 15140122). UWB1.289, which harbors pathogenic *BRCA1* variant, and its *BRCA1*-restored counterpart, and UWB1.289 with *BRCA1* construct (referred to as UWB1 and UWB1+BRCA1, respectively), were cultured in 50% RPMI 1640 and 50% MEM basal media and SingleQuot additives, 200 μg/ml G-418 and 3% FBS.

### Bisulfite conversion and *meBRCA1* targeted sequencing

Genomic DNA extracted from cell lines underwent bisulfite conversion using the EZ DNA Methylation-Lightning Kit (Zymo Research, Cat# D5030), according to the manufacturer’s instructions. Targeted bisulfite sequencing of the *BRCA1* promoter was performed using the two-step PCR protocol described by Nesic et al. (23). In the first PCR, the *BRCA1* promoter region (GRCh38; chr17:43,125,457–43,125,336) was amplified using primer sequences originally designed by Wong et al. (13) with additional Illumina sequencing adapters. The second PCR step added indexes for multiplexing (Illumina DNA/RNA UD indexes Set C; Ref 20091648 or custom adapters). Both PCR steps used HotStarTaq DNA polymerase (QIAGEN; Cat #203203). Sequencing was performed with NextSeq 2000 or MiSeq using 2x150 bp reads to a minimum target depth of 8,000. The sequencing data was analyzed using MethAmplicons2, with a minimum read frequency threshold of 0.01.

### RNA sequencing

RNA was extracted from frozen PDX tissue (#11, #48, #62, #169, #334, #428, #495, #585, and #951), single replicate per model. Samples were sequenced in two batches (Supplementary Material 1). For cell lines, technical replicates (n = 3) of OVCAR8, and OVCAR8 A6-meth were independently cultured. RNA extraction was performed with Quick-RNA Miniprep Kit (ZYMO RESEARCH; Cat #R1054). The integrity of the extracted RNA was assessed using the Agilent TapeStation, with all samples achieving RNA integrity number scores above 8.5. mRNA library preparation was performed using the Illumina TruSeq Stranded mRNA Library Prep Kit, followed by sequencing on the NextSeq 2000 platform as paired-end 100bp reads (PDX: 42.8– 68.9 million read pairs per sample, OVCAR8: 68.4 – 74.8 million read pairs per sample).

Briefly, bulk RNA sequencing data were converted from BCL to FASTQ using bcl2fastq (Illumina). Cutadapt (v1.9) (24) was used to trim adapter sequences. Read groups from cell-lines were aligned to the GRCh38 reference (Ensembl v110), while read groups from PDX were aligned to a hybrid reference where GRCh38 was concatenated with GRCm38 (Ensembl v70). The alignment was carried out using STAR (v2.7.10a) in paired-end (PE) mode with quantMode implemented to output alignments in transcript coordinates. Per sample aligned read groups were merged using samtools (v1.9) (25), and then gene-level expected counts were estimated using RSEM (v1.2.30) (26).

Transcripts per million (TPM) values were computed from gene-level counts in edgeR (v3.40.2)(27) using a library-size–normalized approach incorporating a prior count to stabilize low-expression estimates. Specifically, counts were scaled by effective library sizes (adjusted by normalization factors), a prior count of 2 was added proportionally to library size, and expression values were converted to FPKM-like units prior to normalization by gene length to obtain TPM. Log-transformed expression values were calculated as log2(TPM).

### Whole genome short read sequencing and analysis of patient samples

The patient tumor used to generate PDX #48, along with matched blood sample, underwent whole-genome sequencing (WGS). Genomic DNA (2 μg) was used for library preparation with the TruSeq Nano kit (Illumina). Libraries were sequenced using 150 bp paired-end reads on Illumina HiSeq X Ten or NovaSeq platform, targeting a mean read depth of 30-40× for primary tumor samples and 30× for matched PDX and normal samples.

Sequencing reads were processed using Cutadapt (v1.18) (24) to trim low-quality bases from the 3′ end (-q 20) and remove adapter sequences. Cleaned reads were aligned to a combined human/mouse reference genome (GRCh38/GRCm38, NodShiLtJ background) using BWA-MEM (v0.7.15) (28). Resulting alignments were sorted and indexed with SAMtools (v1.9) (25), and duplicate reads were marked using Picard MarkDuplicates (v1.97) (29). For human mutation analysis, only read pairs mapped to the human reference with a mapping quality score of 60 were retained. Quality control and coverage assessment were performed using in-house tools, qProfiler and qCoverage, as previously described (30).

### BRCA1 Immunohistochemistry

BRCA1 was detected using a mouse anti-BRCA1 C-terminus primary antibody (Santa Cruz Biotechnology, sc-6954). Briefly, 4µm sections of FFPE PDX blocks were deparaffinised using an autostainer (LEICA Biosystems, VIC, Australia). Antigen retrieval was performed at 100°C for 20 minutes using a decloaking chamber (Biocare Medical, CA, USA), followed by a peroxidase block in 3% H2O2 for 10 minutes. The sections were washed in dH2O for 10 minutes and then blocked in blocking buffer (1X TBST + 1% BSA) for 10 minutes. The remaining protocol was carried out as described in Cruz et al.(31) Biocare Medical’s Da Vinci Green (PD900M) was used to dilute the primary antibody and MACH 2 Mouse HRP-Polymer (MHRP520) was used as the secondary antibody. Visualisation was performed using ImmPACT® DAB Substrate, Peroxidase (HRP) (Vector Laboratories, SK-4105). Haematoxylin staining was performed using an autostainer and sections were imaged using Aperio AT turbo (Leica Biosystems) and annotated using ImageJ.

### CUT&RUN assay

CUTANA CUT&RUN (EpiCypher) libraries were prepared following the manufacturer’s instructions. Briefly, cell lines (WEHI-CS62, OVCAR8, OVCAR8 A6-meth, and UWB1) were first cultured as n = 3 independent replicates. Cell permeabilization conditions were determined, with 0.01% digitonin selected as the optimal concentration for each line. Following the manufacturer’s instructions, 0.5 million cells were used with spike-in K-MetStat Panel and E. coli DNA in each condition, with the rest of the protocol unmodified. EpiCypher antibodies certified for CUT&RUN were used for H3K4me3 (positive control - 13-0041k), H3K4me1 (13–0057), H3K27me3 (13–0055), and IgG (negative control - 13-0042k) at 0.5 μg (1 μL) per reaction. For the H3K9me3 antibody (ActiveMotif, 39062), 1 μg (1 μL) was used per reaction. Sequencing libraries were quantified using the Qubit HS dsDNA assay, and library quality was assessed using an Agilent TapeStation with HS D1000 tape. Libraries were sequenced on the NextSeq 2000 platform as paired-end 50 or 150 bp reads to a minimum depth of 8.5 million read pairs per library.

After basecalling, reads were aligned to the human GRCh38 reference using bowtie2 (v 2.2.9)(32) using end-to-end, very sensitive, paired-end mode. Aligned reads were filtered with samtools (v1.20)(25) to retain paired reads with good mapping quality (MAPQ ≥30). PICARD (v2.27.4) (29) was used to mark duplicated reads to be removed during peak calling by MACS3 (v3.0.3)(33). Broad peaks were called for repressive marks H3K27me3 and H3K9me3, while narrow peaks were called for active marks H3K4me1 and H3K4me3. Read coverage was visualized at *BRCA1* region. deepTools (v3.5.6) was used for reads per genomic content (RPGC) normalization and visualization at peak regions.

### Nanopore long-read sequencing

Genomic DNA was extracted from cell pellets or frozen tissue using the QIAamp DNA Mini Kit (QIAGEN, Cat# 51304) according to the manufacturer’s instructions. DNA concentration was measured using the Qubit dsDNA High Sensitivity assay on a Qubit 3.0 fluorometer (Life Technologies). DNA purity was assessed using an Implen NanoPhotometer® NP80, and DNA size distribution was evaluated using the genomic DNA quality assay on an Agilent TapeStation.

High-molecular-weight DNA was adjusted to 40 ng/µL (50 µL total) and fragmented using a Covaris g-TUBE (5,500 rpm for 60 s). Libraries were prepared using the Oxford Nanopore Technologies (ONT) Ligation Sequencing Kit (SQK-LSK114). Each library was sequenced on a PromethION device using a single R10.4.1 flow cell per library.

Nanopore reads in POD5 format were basecalled using Dorado (v0.7.1) with the SUP model (dna_r10.4.1_e8.2_400bps_sup@v5.0.0). Base modifications (5mC and 5hmC) were detected using the integrated Remora framework with the corresponding pretrained model (dna_r10.4.1_e8.2_400bps_sup@v5.0.0_5mCG_5hmCG@v1).

Read trimming parameters were selected based on NanoQC reports (34), 40 nucleotides were removed from the 5′ end and 20 nucleotides from the 3′ end of each read using Chopper (v0.7.0). Reads were aligned using minimap2 (v2.27) (35). For cell line samples, the reference genome was human GRCh38 (GCA_000001405.15). For PDX samples, alignment was performed against a combined human–mouse reference (GRCh38 + GRCm39) and reads aligning preferentially to the mouse reference were removed from downstream analyses. Alignments were sorted and indexed using samtools (v1.20) (25).

Per-read and pileup summaries of 5mC and 5hmC modification calls at CpG sites were generated using Modkit (v0.2.4). Only CpG calls with base modification probabilities >70% were retained. Downstream methylation analyses were restricted to CpG sites with coverage of ≥5 reads and ≤100 reads, and base calls with a minimum Phred quality score ≥20. Data quality was evaluated using read depth, read length distributions, and base quality distributions (Supplementary Material 2).

Small variants (germline and somatic SNVs/indels) were called using Clair3 (v1.0.4) (36). The identified variants were annotated using gnomAD (v3.0) (37) by VEP (v102) (38). The mapped reads were haplo-tagged by whatshap (v2.2) (39) based on phased variants identified by SNV and indels called from Clair3. Somatic only SNVs and indels were predicted with ClairS-TO (v.0.4.1) (40). Throughout variant analyses, only bases with a Phred quality score of ≥20 were considered. Only variants present in ≥10% of sequencing reads and passing Clair3 quality filters are included. For variants within the *BRCA1* promoter region, the presence of large-scale loss of heterozygosity prompted additional curation. Heterozygous variants detected by Clair3 in more than half of the samples were manually reviewed for potential miscalls and excluded where necessary.

Somatic structural variants (SVs) were identified using Severus (v1.5) (41) and NanomonSV (v0.7.1) (42) were used. Following recommendations by Liu et al. (43), Severus was run in tumor-only mode using default settings, whereas NanomonSV was run with a pre-built control panel (44). Events detected by both callers, predicted to be somatic, and not overlapping variable number tandem repeats were considered high-confidence and used for analyses linking methylation changes to sequence alterations. Breakpoints were visualized using IGV (v2.16.0) (44). Somatic copy number alterations were estimated using Savana (v1.3.1) (45).

### Adaptive and native barcoding of Nanopore long-read sequencing

For PDX #585 and guide-edited A6–meth samples, a targeted multiplexing and enrichment strategy was used to increase sequencing coverage across the *BRCA1* locus. Genomic DNA was adjusted to 56–66 ng/µL (65 µL total input) and fragmented using a Megaruptor 3 (Diagenode) at a shear speed of 55 to achieve 10 kb fragment sizes that is optimal for adaptive sequencing.

Library preparation was performed using the Oxford Nanopore Native Barcoding Kit (lot: NBD1424.60.0003–1076), following the manufacturer’s instructions. A single-line target BED file (chr17:33044295–53125364) was provided to enable adaptive sequencing for on-device enrichment of the *BRCA1* region. The pooled library was loaded onto the flow cell three times, with fresh library added at a concentration of 31 ng/µL for each loading. Raw POD5 files were basecalled and demultiplexed using Dorado (v0.7.1), as described above.

### HRD scarring assessment

HRDetect was run following the HRDetect pipeline scripts (https://github.com/eyzhao/hrdetect-pipeline) (46). Somatic indel, SNVs, CNV and SV for this pipeline were generated as described above. The contribution of each sample’s SNVs (derived from ClairS-TO (v.0.4.1)) to the 30 known COSMIC v2 SBS signatures was estimated using deconstructSigs (v1.8.0) (47). We used the COSMIC SBS v2 reference instead of the more recent v3, since the HRDetect framework was originally developed and validated using v2, and based on findings from study by Ding et al. (48), where some samples previously classified with SBS Signature 3 (v2) were reassigned to SBS Signature 39 under v3.

### Global differential methylation analysis

Global differential methylation analysis (DMA) was performed for autosomal CpG sites using methylKit (v1.28.0) across the following isogenic pairs: OVCAR8 vs OVCAR8 A6-meth, OVCAR8 vs OVCAR8 H4-KO, and UWB1+BRCA1 vs UWB1. A total of 26,197,298 autosomal CpG loci were included in the analysis. Differential methylation was assessed at single-CpG resolution using methylKit’s logistic regression framework and multiple-testing correction was applied using FDR with a threshold of 10% for methylation difference and 0.05 for FDR.

### Copy number adjusted methylation analysis at promoters and CpG islands

CpGs loci were filtered to retain those that overlapped with promoters and/or CpG islands. Promoter coordinates were obtained from the Eukaryotic Promoter Database (EPDnew) (human version 006) (49). CpG island information was obtained from the UCSC genome browser (50,51). Analyses were restricted to autosomal CpG loci within promoters and/or CpG islands ±4 kb.To account for copy-number alterations, promoter/CpG island methylation was assessed incorporating copy-number. Copy-number–adjusted methylation was calculated as: adjusted methylation = ⌈fraction methylation × copy number⌉/copy number.

Given the limited sample size, formal statistical testing of partial methylation shifts from fully methylated loci was not performed. Instead, loci were categorized by consistency across groups. Loci with full methylation (adjusted methylation = 1) in homozygous *meBRCA1* PDX models (#11 and #62) and partial or absent methylation (0 ≤ adjusted methylation < 1)in at least one het *meBRCA1* PDX were retained. Conversely, loci with no methylation (adjusted methylation = 0) in both homozygous *meBRCA1* PDX models and partial or full methylation in at least one het *meBRCA1* PDX were also retained. Only genes containing ≥3 consecutive promoter CpG loci with concordant directional change were included. The functional impact of methylation changes was estimated and visualized by mRNA expression, only genes with concordant direction of mRNA expression differences were included in the final list.

### Global analysis for sequence alterations and methylation

Somatic clonal sequence alteration events, including indels, large deletions, large insertions, duplications, and translocations, were identified as events present exclusively in engineered lines (OVCAR8 H4-KO, OVCAR8 A6-meth, and UWB1+BRCA1) and absent in their corresponding parental lines (OVCAR8 and UWB1). We firstly examined the methylation stability using heterozygous variants (i.e., present on a subset of alleles) and confirmed methylation stability of reference allele between engineered and parental lines (Supplementary Material 3). High-confidence heterozygous events were defined as those with ≥4 reads supporting the alternative allele and ≥4 reads supporting the reference allele.

CpG loci within ±5,000 bp of each sequence alteration breakpoint were interrogated for allele-specific methylation changes. Analyses were restricted to CpGs that were highly methylated (≥80%) or lowly methylated (≤20%) in the parental line, to focus on loci with clearly defined baseline methylation states. To exclude CpGs prone to methylation drift unrelated to the alteration, CpGs were removed if methylation levels in REF reads from engineered lines differed from parental methylation levels by an absolute difference of ≥20%.

For each retained CpG, association between sequence alteration status and methylation state was assessed using Fisher’s exact test, comparing methylation calls between ALT and REF allele-supporting reads. Multiple-testing correction was applied per event using Benjamini– Hochberg. Events were considered to show methylation alteration if they contained ≥1 CpG with adjusted p-value ≤0.05 and an absolute methylation difference ≥80% between ALT and REF alleles.

Permutation testing was performed to evaluate whether methylation-changed CpGs were non-randomly distributed relative to breakpoints. Using the R package coin (v1.4-3), CpGs within ±5 kb of each breakpoint were randomly relabeled (“methylation changed” vs “not changed”) 100,000 times while keeping genomic positions fixed. For each iteration, the difference in mean breakpoint-to-CpG distance between the two groups was recomputed to generate a null distribution.

Each side of a breakpoint from a sequence alteration event was annotated for the presence of overlap with the following genomic features: CpG islands (CGIs), shores, shelves, and open sea were annotated using the UCSC Genome Browser CGI tracks (50). Shores were defined as 2 kb regions flanking CGIs, and shelves were defined as 2 kb regions flanking shores. For breakpoints overlapping multiple CGI features, a prioritization hierarchy was applied: island>shore>shelf; breakpoints without annotation were assigned open sea.

Gene context was annotated using the EPDnew database (GRCh38 alignment) for promoters (51), Ensembl v112 for gene bodies, and all other regions were classified as intergenic. Repetitive elements were annotated using UCSC Genome Browser tracks (52). For breakpoints overlapping multiple repeat types, a prioritization hierarchy was applied: LINE>SINE>LTR>DNA; breakpoints without annotated repeats were assigned NA. lncRNA presence was annotated using lncRNAKB (53). Breakpoints were classified using a binary indicator for the presence or absence of overlap with lncRNA transcripts.

Statistical significance was assessed using multinomial logistic regression implemented in the R package VGAM (v1.1-12).

### CRISPR guided deletion

CRISPR RNAs (crRNAs) were designed using the Integrated DNA Technologies (IDT) “Design custom gRNA” tool to target the first intron of *BRCA1*. Guide #1 (CTGGCACCTCTTCTTCCACA) targeted chr17:43124694-43124713 (+), guide #2 (CACCGGCTGGTATGTATGAG) targeted chr17:43124726-43124745 (-), guide #3 targeted (CTATCACGAGGATTCCCCCA) chr17:43125036-43125055 (+). Each crRNA was combined with Alt-R CRISPR-Cas9 tracrRNA, ATTO 550 (IDT, Cat #1075927) at equimolar amounts and incubated at 95°C for 5 min. RNP complex was formed by combining 1.2 µL of crRNA:tracrRNA duplex with 1.7 µL Alt-R S.p. HiFi Cas9 Nuclease V3 (IDT, Cat #1081061) and 2.1 µL of DPBS (Gibco, Cat #14190-144) and incubating at room temperature for 20 min. 1 × 10^5^ OVCAR8 A6-meth cells were resuspended in 20 µL of nucleofector solution combined with supplement at 4.5:1 from the Amaxa SE Cell Line 4D-Nucleofector X Kit S (Lonza, Cat #V4XC-1032), before addition of 5 µL RNP complex and 1 µl of Electroporation enhancer. 25 µL of this mix was transferred to a nucleocuvette strip and run on program CA137 in a 4D Nucleofector (Amaxa) before transfer to culture medium for expansion.

### Olaparib selection of CRISPR edited OVCAR8 A6-meth clones

CRISPR-edited OVCAR8 A6-meth cells generated using guide #1 or guide #2 were subjected to olaparib selection to enrich for subpopulations consistent with BRCA1 promoter methylation loss. The parental OVCAR8 A6-meth line was used as control in parallel. All experiments were performed in independent triplicate for olaparib selection.

Cells were seeded at approximately 20% confluency and allowed to attach for 24 hours. Cells were treated with 0.15 μM olaparib (IC80 of parental OVCAR8 A6-meth) for ten days, with twice weekly passaging. Following olaparib treatment, surviving cells were allowed to recover for 24 hours before further analysis.

RAD51 foci was performed as previously described (54) on cells grown to 80% confluency on coverslips.

For dose–response assays, cells were seeded in white-walled 96-well plates (Corning, NY) at densities of 500 cells per well for OVCAR8 and UWB1.289 cell lines, and 1,500 cells per well for WEHI-CS62 cells. Drugs were added 24 hours after seeding, and plates were incubated for an additional 7 days. Drug concentrations ranged from 0.34 nM to 5 μM for both olaparib and cisplatin. Vehicle controls consisted of DMSO at the corresponding highest concentration.

Cell viability was assessed using the CellTiter-Glo® 2.0 assay (Promega, Madison, WI) at a reagent-to-medium ratio of 1:3. Luminescence was measured using a BioTek Synergy 2 plate reader.

## Acknowledgements

We would like to acknowledge all members of the Molecular Oncology, Medical Genomics, Genome Informatics laboratories and Scientific Services teams at QIMR Berghofer for their technical support. We would specifically like to thank Brett Liddell for updating MethAmplicons code, Nadine Fitzpatrick and Jenny Quiatchon for setting up and optimising long-read sequencing, Emma Todman for conducting cell line treatment experiments. We would like to thank Oxford Nanopore Technologies for providing trial reagents and technical advice on multiplexing and adaptive sampling. We would like to thank Clovis Oncology for provision of drug rucaparib for in vivo treatments. We gratefully acknowledge Silvia Stoev, Chloe Neagle and Stephanie Bound for technical assistance with the animal studies, and the Walter and Eliza Hall Institute of Medical Research (WEHI) Bioservices and Advanced Histotechnology Facility for their support and assistance in this work. All authors, the WEHI Stafford Fox Rare Cancer Program and the AOCS would like to thank all of the patients who participated in these research programs. The AOCS would also like to acknowledge the contribution of the study nurses, research assistants, and all clinical and scientific collaborators to the study. The complete AOCS Study Group can be found at www.aocstudy.org. The Australian Ovarian Cancer Study gratefully acknowledges additional support from Ovarian Cancer Australia and the Peter MacCallum Foundation. The Australian Ovarian Cancer Study Group was supported by the U.S. Army Medical Research and Materiel Command under DAMD17-01-1-0729, The Cancer Council Victoria, Queensland Cancer Fund, The Cancer Council New South Wales, The Cancer Council South Australia, The Cancer Council Tasmania and The Cancer Foundation of Western Australia (Multi-State Applications 191, 211 and 182) and the National Health and Medical Research Council of Australia (NHMRC; ID199600; ID400413 and ID400281).

## Availability of data and materials

All sequencing data will be available from European Genome-phenome Archive (EGA) repository at the time of publication.

## Code availability

The code used to perform all genomic and epigenomic analyses and generate figures is available here: https://github.com/molonc-lab/MeBRCA1_instability.

## References

1. Patch AM, Christie EL, Etemadmoghadam D, Garsed DW, George J, Fereday S, et al. Whole-genome characterization of chemoresistant ovarian cancer. Nature 2015;521(7553):489–94.

2. Kondrashova O, Topp M, Nesic K, Lieschke E, Ho GY, Harrell MI, et al. Methylation of all BRCA1 copies predicts response to the PARP inhibitor rucaparib in ovarian carcinoma. Nat Commun 2018;9(1):3970 doi 10.1038/s41467-018-05564-z.

3. Menghi F, Banda K, Kumar P, Straub R, Dobrolecki L, Rodriguez IV, et al. Genomic and epigenomic BRCA alterations predict adaptive resistance and response to platinum-based therapy in patients with triple-negative breast and ovarian carcinomas. Sci Transl Med 2022;14(652):eabn1926 doi 10.1126/scitranslmed.abn1926.

4. Glodzik D, Bosch A, Hartman J, Aine M, Vallon-Christersson J, Reuterswärd C, et al. Comprehensive molecular comparison of BRCA1 hypermethylated and BRCA1 mutated triple negative breast cancers. Nature Communications 2020;11(1):3747 doi 10.1038/s41467-020-17537-2.

5. Kalachand RD, Stordal B, Madden S, Chandler B, Cunningham J, Goode EL, et al. BRCA1 Promoter Methylation and Clinical Outcomes in Ovarian Cancer: An Individual Patient Data Meta-Analysis. J Natl Cancer Inst 2020;112(12):1190–203 doi 10.1093/jnci/djaa070.

6. Chiang JW, Karlan BY, Cass L, Baldwin RL. BRCA1 promoter methylation predicts adverse ovarian cancer prognosis. Gynecol Oncol 2006;101(3):403–10 doi 10.1016/j.ygyno.2005.10.034.

7. Swisher EM, Kwan TT, Oza AM, Tinker AV, Ray-Coquard I, Oaknin A, et al. Molecular and clinical determinants of response and resistance to rucaparib for recurrent ovarian cancer treatment in ARIEL2 (Parts 1 and 2). Nat Commun 2021;12(1):2487 doi 10.1038/s41467-021-22582-6.

8. Swisher EM, Sakai W, Karlan BY, Wurz K, Urban N, Taniguchi T. Secondary BRCA1 mutations in BRCA1-mutated ovarian carcinomas with platinum resistance. Cancer Res 2008;68(8):2581–6 doi 10.1158/0008-5472.CAN-08-0088.

9. Privat M, Radosevic-Robin N, Aubel C, Cayre A, Penault-Llorca F, Marceau G, et al. BRCA1 induces major energetic metabolism reprogramming in breast cancer cells. PLoS One 2014;9(7):e102438 doi 10.1371/journal.pone.0102438.

10. Bruand M, Barras D, Mina M, Ghisoni E, Morotti M, Lanitis E, et al. Cell-autonomous inflammation of BRCA1-deficient ovarian cancers drives both tumor-intrinsic immunoreactivity and immune resistance via STING. Cell Rep 2021;36(3):109412 doi 10.1016/j.celrep.2021.109412.

11. Chen T, Yu T, Zhuang S, Geng Y, Xue J, Wang J, et al. Upregulation of CXCL1 and LY9 contributes to BRCAness in ovarian cancer and mediates response to PARPi and immune checkpoint blockade. British Journal of Cancer 2022;127(5):916–26 doi 10.1038/s41416-022-01836-0.

12. Lee CK, Kartikasari AER, Bound NT, Francis KE, Shield-Artin K, Bedo J, et al. Olaparib, durvalumab, and cyclophosphamide, and a prognostic blood signature in platinum-sensitive ovarian cancer: the randomized phase 2 SOLACE2 trial. Nat Commun 2025;16(1):9756 doi 10.1038/s41467-025-64130-6.

13. Wong EM, Southey MC, Fox SB, Brown MA, Dowty JG, Jenkins MA, et al. Constitutional methylation of the BRCA1 promoter is specifically associated with BRCA1 mutation-associated pathology in early-onset breast cancer. Cancer Prev Res (Phila) 2011;4(1):23–33 doi 10.1158/1940-6207.CAPR-10-0212.

14. Nesic K, Beard S, Xu L, Kondrashova O, Vandenberg CJ, Rubin AF, et al. Generation of a PARPi-sensitive homozygous BRCA1-methylated OVCAR8 cell line using targeted CRISPR gene editing. bioRxiv 2024:2024.09. 23.614644.

15. Xu L, Liddell B, Nesic K, Geissler F, Ashwood LM, Wakefield MJ, et al. High-level tumour methylation of BRCA1 and RAD51C is required for homologous recombination deficiency in solid cancers. NAR Cancer 2024;6(3):zcae033 doi 10.1093/narcan/zcae033.

16. Rothbart SB, Krajewski K, Nady N, Tempel W, Xue S, Badeaux AI, et al. Association of UHRF1 with methylated H3K9 directs the maintenance of DNA methylation. Nat Struct Mol Biol 2012;19(11):1155–60 doi 10.1038/nsmb.2391.

17. van Schendel R, van Heteren J, Welten R, Tijsterman M. Genomic Scars Generated by Polymerase Theta Reveal the Versatile Mechanism of Alternative End-Joining. PLOS Genetics 2016;12(10):e1006368 doi 10.1371/journal.pgen.1006368.

18. Swisher EM, Lin KK, Oza AM, Scott CL, Giordano H, Sun J, et al. Rucaparib in relapsed, platinum-sensitive high-grade ovarian carcinoma (ARIEL2 Part 1): an international, multicentre, open-label, phase 2 trial. The Lancet Oncology 2017;18(1):75–87.

19. Ter Brugge P, Kristel P, van der Burg E, Boon U, de Maaker M, Lips E, et al. Mechanisms of Therapy Resistance in Patient-Derived Xenograft Models of BRCA1-Deficient Breast Cancer. J Natl Cancer Inst 2016;108(11) doi 10.1093/jnci/djw148.

20. Nesic K, Geissler F, Xu L, Kyran E, Beard S, Vandenberg CJ, et al. Demethylating agents drive PARP inhibitor resistance in ovarian carcinomas with BRCA1 gene silencing. bioRxiv 2025:2025.04. 07.647418.

21. Topp MD, Hartley L, Cook M, Heong V, Boehm E, McShane L, et al. Molecular correlates of platinum response in human high-grade serous ovarian cancer patient-derived xenografts. Mol Oncol 2014;8(3):656–68.

22. Wakefield MJ. Xenomapper: Mapping reads in a mixed species context. Journal of Open Source Software 2016;1(1):18 doi 10.21105/joss.00018.

23. Nesic K, Kondrashova O, Hurley RM, McGehee CD, Vandenberg CJ, Ho GY, et al. Acquired RAD51C Promoter Methylation Loss Causes PARP Inhibitor Resistance in High-Grade Serous Ovarian Carcinoma. Cancer Res 2021;81(18):4709–22 doi 10.1158/0008-5472.CAN-21-0774.

24. Martin M. Cutadapt removes adapter sequences from high-throughput sequencing reads. 2011 2011;17(1):3 doi 10.14806/ej.17.1.200.

25. Danecek P, Bonfield JK, Liddle J, Marshall J, Ohan V, Pollard MO, et al. Twelve years of SAMtools and BCFtools. GigaScience 2021;10(2):giab008 doi 10.1093/gigascience/giab008.

26. Li B, Dewey CN. RSEM: accurate transcript quantification from RNA-Seq data with or without a reference genome. BMC Bioinformatics 2011;12(1):323 doi 10.1186/1471-2105-12-323.

27. Robinson MD, McCarthy DJ, Smyth GK. edgeR: a Bioconductor package for differential expression analysis of digital gene expression data. Bioinformatics 2010;26(1):139–40 doi 10.1093/bioinformatics/btp616.

28. Li H. Aligning sequence reads, clone sequences and assembly contigs with BWA-MEM. ArXiv 2013;1303 doi 10.48550/arXiv.1303.3997.

29. Picard toolkit. Broad Institute, GitHub repository https://broadinstitute.github.io/picard/:BroadInstitute; 2019.

30. Bonazzi VF, Kondrashova O, Smith D, Nones K, Sengal AT, Ju R, et al. Patient-derived xenograft models capture genomic heterogeneity in endometrial cancer. Genome Medicine 2022;14(1):3 doi 10.1186/s13073-021-00990-z.

31. Cruz C, Castroviejo-Bermejo M, Gutierrez-Enriquez S, Llop-Guevara A, Ibrahim YH, Gris-Oliver A, et al. RAD51 foci as a functional biomarker of homologous recombination repair and PARP inhibitor resistance in germline BRCA-mutated breast cancer. Ann Oncol 2018;29(5):1203–10 doi 10.1093/annonc/mdy099.

32. Langmead B, Salzberg SL. Fast gapped-read alignment with Bowtie 2. Nature Methods 2012;9(4):357–9 doi 10.1038/nmeth.1923.

33. Zhang Y, Liu T, Meyer CA, Eeckhoute J, Johnson DS, Bernstein BE, et al. Model-based Analysis of ChIP-Seq (MACS). Genome Biology 2008;9(9):R137 doi 10.1186/gb-2008-9-9-r137.

34. De Coster W, D’Hert S, Schultz DT, Cruts M, Van Broeckhoven C. NanoPack: visualizing and processing long-read sequencing data. Bioinformatics 2018;34(15):2666–9 doi 10.1093/bioinformatics/bty149.

35. Li H. Minimap2: pairwise alignment for nucleotide sequences. Bioinformatics 2018;34(18):3094–100 doi 10.1093/bioinformatics/bty191.

36. Zheng Z, Li S, Su J, Leung AW-S, Lam T-W, Luo R. Symphonizing pileup and full-alignment for deep learning-based long-read variant calling. bioRxiv 2022:2021.12.29.474431 doi 10.1101/2021.12.29.474431.

37. Ellrott K, Bailey MH, Saksena G, Covington KR, Kandoth C, Stewart C, et al. Scalable Open Science Approach for Mutation Calling of Tumor Exomes Using Multiple Genomic Pipelines. Cell Syst 2018;6(3):271–81.e7 doi 10.1016/j.cels.2018.03.002.

38. McLaren W, Gil L, Hunt SE, Riat HS, Ritchie GR, Thormann A, et al. The Ensembl Variant Effect Predictor. Genome Biol 2016;17(1):122 doi 10.1186/s13059-016-0974-4.

39. Patterson M, Marschall T, Pisanti N, van Iersel L, Stougie L, Klau GW, et al. WhatsHap: Weighted Haplotype Assembly for Future-Generation Sequencing Reads. Journal of Computational Biology 2015;22(6):498–509 doi 10.1089/cmb.2014.0157.

40. Chen L, Zheng Z, Su J, Yu X, Wong AOK, Zhang J, et al. ClairS-TO: A deep-learning method for long-read tumor-only somatic small variant calling. bioRxiv 2025:2025.03.10.642523 doi 10.1101/2025.03.10.642523.

41. Keskus A, Bryant A, Ahmad T, Yoo B, Aganezov S, Goretsky A, et al. Severus: accurate detection and characterization of somatic structural variation in tumor genomes using long reads. medRxiv 2024:2024.03.22.24304756 doi 10.1101/2024.03.22.24304756.

42. Shiraishi Y, Koya J, Chiba K, Okada A, Arai Y, Saito Y, et al. Precise characterization of somatic complex structural variations from tumor/control paired long-read sequencing data with nanomonsv. Nucleic Acids Research 2023;51(14):e74-e doi 10.1093/nar/gkad526.

43. Liu L, Zhang J, Wood S, Newell F, Leonard C, Koufariotis LT, et al. Performance of somatic structural variant calling in lung cancer using Oxford Nanopore sequencing technology. BMC Genomics 2024;25(1):898 doi 10.1186/s12864-024-10792-3.

44. Shafin K, Pesout T, Lorig-Roach R, Haukness M, Olsen HE, Bosworth C, et al. Nanopore sequencing and the Shasta toolkit enable efficient de novo assembly of eleven human genomes. Nature Biotechnology 2020;38(9):1044–53 doi 10.1038/s41587-020-0503-6.

45. Elrick H, Sauer CM, Espejo Valle-Inclan J, Trevers K, Tanguy M, Zumalave S, et al. SAVANA: reliable analysis of somatic structural variants and copy number aberrations using long-read sequencing. Nature Methods 2025;22(7):1436–46 doi 10.1038/s41592-025-02708-0.

46. Zhao EY, Shen Y, Pleasance E, Kasaian K, Leelakumari S, Jones M, et al. Homologous Recombination Deficiency and Platinum-Based Therapy Outcomes in Advanced Breast Cancer. Clin Cancer Res 2017;23(24):7521–30 doi 10.1158/1078-0432.Ccr-17-1941.

47. Rosenthal R, McGranahan N, Herrero J, Taylor BS, Swanton C. DeconstructSigs: delineating mutational processes in single tumors distinguishes DNA repair deficiencies and patterns of carcinoma evolution. Genome Biol 2016;17:31 doi 10.1186/s13059-016-0893-4.

48. Ding YC, Tao S, Mao A, Ziv E, Neuhausen SL. Cosmic signature SBS39 is associated with homologous recombination deficiency. medRxiv 2024:2024.10.23.24316019 doi 10.1101/2024.10.23.24316019.

49. Dreos R, Ambrosini G, Cavin Périer R, Bucher P. EPD and EPDnew, high-quality promoter resources in the next-generation sequencing era. Nucleic Acids Research 2013;41(D1):D157–D64 doi 10.1093/nar/gks1233.

50. Gardiner-Garden M, Frommer M. CpG Islands in vertebrate genomes. Journal of Molecular Biology 1987;196(2):261–82 doi 10.1016/0022-2836(87)90689-9.

51. Perez G, Barber GP, Benet-Pages A, Casper J, Clawson H, Diekhans M, et al. The UCSC Genome Browser database: 2025 update. Nucleic Acids Res 2025;53(D1):D1243–d9 doi 10.1093/nar/gkae974.

52. Jurka J. Repbase update: a database and an electronic journal of repetitive elements. Trends Genet 2000;16(9):418–20 doi 10.1016/s0168-9525(00)02093-x.

53. Seifuddin F, Singh K, Suresh A, Judy JT, Chen Y-C, Chaitankar V, et al. lncRNAKB, a knowledgebase of tissue-specific functional annotation and trait association of long noncoding RNA. Scientific Data 2020;7(1):326 doi 10.1038/s41597-020-00659-z.

54. Wang L, Bitar M, Lu X, Jacquelin S, Nair S, Sivakumaran H, et al. CRISPR-Cas13d screens identify KILR, a breast cancer risk-associated lncRNA that regulates DNA replication and repair. Mol Cancer 2024;23(1):101 doi 10.1186/s12943-024-02021-y.

