## Supplementary Figures for "Genomic alterations enable *BRCA1* methylation loss and promoter bypass to drive resistance in high-grade serous ovarian cancer"

#### Supplementary Figure legends

**Supplementary Figure 1. HRD prediction of models using HRDetect.** **(A)** HRDetect scores for each model. The horizontal dashed line indicates the HRDetect threshold (≥0.7, HRD; <0.7, HRP). Bars are colored by *BRCA1* status: heterozygous promoter methylation (het me*BRCA1*, orange), homozygous promoter methylation (hom me*BRCA1*, yellow), loss-of-function *BRCA1* mutation (mut *BRCA1*, dark green), and wild-type *BRCA1* (wt *BRCA1*, green). **(B)** Contributions of individual HRD score components, including loss of heterozygosity (LOH, green), large-scale transitions (LST, yellow), and telomeric allelic imbalance (TAI, black). **(C)** Contributions of COSMIC single-base substitution signatures (SBS v2). The y-axis indicates the proportion of signatures with contributions >0.1, including SBS3 (HRD-associated, burgundy), SBS4 (smoking-associated, light green), SBS5 (ubiquitous, green), SBS1 (age-associated, turquoise), and SBS8, SBS12, and SBS16 (unknown etiology, grey). **(D)** Proportion of indels exhibiting microhomology across models. **(E)** Contributions of HRDetect rearrangement signatures. The y-axis indicates the proportion of rearrangement signatures with contributions >0.1, including HRD-associated RS3 (lilac) and RS5 (purple), and RS2 and RS4 (grey). Models are grouped into homologous recombination–proficient negative controls (HRP Control), homologous recombination–deficient positive controls (HR Scar Control), and patient-derived xenograft (PDX) models.

**Supplementary Figure 2. Promoter methylation profiles of *RAD51C*, *BRCA1*, *BRIP1*, and *PALB2* across cell line and PDX models.** Percentage of 5-methylcytosine (5mC) at CpG sites across promoter regions of homologous recombination repair (HRR)–related genes, previously reported to exhibit promoter methylation (Xu et al., 2024), is shown for patient-derived xenograft (PDX) models and ovarian cancer cell line models. For each model, CpG methylation levels across promoter regions previously identified as differentially methylated are plotted against chromosomal coordinates (GRCh38). Each set of connected points represents an individual model. Samples are color-coded by model identity.

**Supplementary Figure 3. Cisplatin responses in *meBRCA1* PDX models.** **(A)** Mean PDX tumor volume (mm3) ± 95% CI (hashed lines are individual mice) and corresponding Kaplan–Meier survival analysis. Censored events are represented by crosses on Kaplan–Meier plot. **(B)** Mean cisplatin response measured by CTG assay across the included cell lines (n=3 independent experiments). IC50 values are 0.144, 0.264, 0.376, 0.637 and 3.34 for OVCAR8 A6-meth, WEHI CS62, UWB1, UWB1+BRCA1 and OVCAR8, respectively.

**Supplementary Figure 4. Loss of heterozygosity across chromosome 17 in all models.** Each model is displayed as two panels. The upper panel shows the allelic frequency (AF) at common single nucleotide polymorphism (SNP) positions derived from the 1000 Genomes Project. The lower panel shows the corresponding $log_2$-transformed read counts (L2R) at each genomic position. Across all models included in this study, a consistent pattern of loss of heterozygosity (LOH) is observed along chromosome 17.

**Supplementary Figure 5. Allele-resolved CpG methylation profiling of the *BRCA1* promoter by Nanopore direct DNA sequencing.** Allele-resolved CpG methylation profiles across the *BRCA1* promoter were determined by Nanopore direct DNA sequencing in cell lines and PDX models. Each line represents an individual DNA molecule, and each dot corresponds to a CpG site, colored by methylation status (methylated, red; unmethylated, blue). Samples are color-coded according to *BRCA1* methylation status: homozygous me*BRCA1* (green), heterozygous me*BRCA1* (brown), and unmethylated *BRCA1* (pink).

**Supplementary Figure 6. Base coverage and upstream structural variants at the *BRCA1* locus in PDX #585 from Nanopore direct RNA sequencing.** Per-base read coverage across the *BRCA1* genomic region on chromosome 17 is shown for PDX #585 using Nanopore direct long-read sequencing. Two upstream structural variants (SVs) with strong read support are indicated by grey dashed lines, while one candidate SV events supported by two reads from whole genome sequencing and ten reads from adaptive sequencing is indicated by red dashed lines. Reference transcript annotations include the MANE-selected *BRCA1* transcript (NM_007294.4; matched annotation from NCBI and EMBL-EBI), the non-coding transcript *NBR2* (ENST00000657841.1), and *NBR1* (NM_005899.5).

**Supplementary Figure 7. Global methylation patterns across PDX and OVCAR8 cell line models. (A)** Distribution of median DNA methylation levels across PDX models. **(B)** Density plots showing genome-wide DNA methylation. Genomic regions were stratified into three categories: CpG-dense regions that are typically highly methylated (CpG islands), large megabase-scale regions lacking CpG islands and exhibiting high methylation variability (partially methylated domains, PMDs), and the remaining genomic regions ("Other"). PDX models are stratified by *BRCA1* promoter methylation zygosity: homozygous methylation (me*BRCA1*; green), heterozygous methylation with silencing (brown), and heterozygous methylation without silencing (grey).

**Supplementary Figure 8. Genes exhibiting reduced promoter methylation loss in heterozygous me*BRCA1* PDX models.** CpG methylation profiles are shown for six genes (*BRCA1*, *BNC1*, *HS3ST2*, *KATNAL2*, *MROH6*, and *SYCP2*) displaying promoter-associated methylation differences between homozygous (hom; N = 2) and heterozygous (het; N = 5 with available RNAseq data) me*BRCA1* PDX models. For each gene, the genomic locus and chromosomal coordinates (GRCh38) are indicated below the plot. The x-axis represents chromosomal position, and the y-axis indicates the percentage of 5-methylcytosine (5mC) at each CpG site. Each set of points connected by lines represents an individual PDX model. The genes exhibited sufficient mRNA expression across the PDX cohort (⩾10 reads mapped to the gene in >70% of samples), with expression levels shown as log-transformed transcripts per million (logTPM). Samples are color-coded by me*BRCA1* zygosity: homozygous me*BRCA1* (green) and heterozygous me*BRCA1* (brown).

**Supplementary Figure 9. Validation of promoter methylation changes in paired primary and relapsed ICGC HGSOC samples. (A)** Two of three paired primary and relapsed ICGC high-grade serous ovarian cancer (HGSOC) samples show loss of *BRCA1* promoter methylation in relapsed tumors. **(B)** Restoration of *BRCA1* mRNA expression in relapsed samples from the same donors. Primary tumor samples are shown as solid lines and relapsed samples as dashed lines. Samples are color-coded by donor identity: AOCS-091 (blue), AOCS-093 (orange), and AOCS-094 (pink). **(C)** Promoter methylation and gene expression changes in paired primary and relapsed ICGC HGSOC samples for genes consistent with me*BRCA1*-linked promoter methylation loss. CpG methylation profiles are shown for five genes (*BNC1, HS3ST2, KATNAL2, MROH6*, and *SYCP2*) selected based on promoter-associated methylation differences between heterozygous and homozygous me*BRCA1* status in PDX models. For each gene, the genomic locus and chromosomal coordinates (GRCh38) are indicated below the plot. The x-axis represents chromosomal position, and the y-axis indicates the percentage of 5-methylcytosine (5mC) at each CpG site. Each connected set of points represents an individual patient sample. For genes with sufficient mRNA expression across samples (⩾10 reads mapped in >70% of samples), corresponding expression levels are shown as log-transformed TPM. *MROH6* expression could not be evaluated due to insufficient detected expression level in the ICGC cohort. **(D)** Correlation between *BRCA1* and *BNC1* promoter methylation in primary tumors and normal tissues from the ICGC HGSOC cohort. Each point represents an individual sample, including primary, relapsed tumors and solid tissue normal samples. The x- and y-axes indicate average promoter methylation levels of *BRCA1* and *BNC1*, respectively. Samples are color-coded by predicted *BRCA1* promoter methylation status: me*BRCA1* (red) and *BRCA1*-unmethylated (blue).

**Supplementary Figure 10. Base-modification profiles across an extended region around *BRCA1* promoter (±10 kb). (A)** Percentage of 5-methylcytosine (5mC) at CpG sites across the *BRCA1* promoter and flanking regions are shown from nine xenografts (PDXs: #11, #48, #62, #69, #428, #495, #585, #334, #865 and #951) and two ovarian cancer cell line models (OVCAR8 and UWB1). Each line represents an individual sample and is colored by me*BRCA1* zygosity: homozygous (blue) and heterozygous (brown). **(B)** Base-modification profiles from the WEHI-CS62 cell line, derived from the same patient material as PDX #62 (purple), and the engineered homozygous me*BRCA1* OVCAR8 A6-meth cell line (green), shown separately for clarity. The engineered homozygous me*BRCA1* OVCAR8 A6-meth cell line has introduced deletion downstream from *BRCA1* gene, aligning with the drop in methylation levels. Modification frequencies were smoothed using a 13-bp Hanning window. The *BRCA1* promoter region and the nearest upstream CpG island are highlighted in each panel. The bottom track indicates the MANE-selected *BRCA1* transcript within the same genomic interval.

**Supplementary Figure 11. Single-nucleotide variants, small insertions and deletion within the *BRCA1* promoter region. (A)** Single-nucleotide variants (SNVs) identified in PDX models and cell line models, grouped and ordered by me*BRCA1* status. Only variants present in ⩾10% of sequencing reads and passing Clair3 quality filters are shown. **(B)** Small insertions (⩽50 bp) detected within the *BRCA1* promoter region across tested models. Insertion size and inferred zygosity are indicated as shown in the legend. **(C)** Small deletions (⩽50 bp) detected within the *BRCA1* promoter region across tested models. Only deletions present in ⩾10% of sequencing reads and passing Clair3 quality filters are shown. Deletion size and inferred zygosity is encoded by symbol size, as indicated in the legend.

**Supplementary Figure 12. Short-read validation of a *BRCA1* structural rearrangement identified by Nanopore sequencing in PDX #48.** Illumina short-read whole-genome sequencing of the patient tumor sample used to generate PDX #48 identified four split reads supporting the translocation detected by Nanopore long-read sequencing in the main figure, while no supporting reads were observed in the matched germline sample. Read-depth coverage (green) and the reciprocal breakpoint locus (blue) are shown across a 5 kb window. Dashed vertical lines indicate breakpoint positions, and bracket annotations denote the number of supporting split reads.

**Supplementary Figure 13. CUT&RUN profiling of repressive histone modifications across the extended *BRCA1* locus in cell line models with distinct *BRCA1* methylation states.** CUT&RUN was performed to profile activating histone marks (H3K4me3 and H3K4me1 – shown in separate supplementary figure), repressive marks (H3K27me3 and H3K9me3), and IgG control across four cell lines: homozygous me*BRCA1* (WEHI-CS62 and OVCAR8 A6-meth), heterozygous me*BRCA1* (OVCAR8), and unmethylated *BRCA1* (UWB1). Reads per genomic content (RPGC)-normalized read coverage is shown across an extended region of the *BRCA1* locus (chr17:42,825,532–43,145,192), including representative regions containing positive enrichment peaks. The lower panel displays reference transcript annotations, including the MANE-selected *BRCA1* transcript (NM_007294.4).

**Supplementary Figure 14. CUT&RUN profiling of active histone modifications across the extended *BRCA1* locus in cell line models with distinct *BRCA1* methylation states.** CUT&RUN was performed to profile activating histone marks (H3K4me3 and H3K4me1), repressive marks (H3K27me3 and H3K9me3 - shown in separate supplementary figure), and IgG control (shown in separate supplementary figure) across four cell lines: homozygous me*BRCA1* (WEHI-CS62 and OVCAR8 A6-meth), heterozygous me*BRCA1* (OVCAR8), and unmethylated *BRCA1* (UWB1). Reads per genomic content (RPGC)-normalized read coverage is shown across an extended region of the *BRCA1* locus (chr17:42,825,532–43,145,192), including representative regions containing positive enrichment peaks. The lower panel displays reference transcript annotations, including the MANE-selected *BRCA1* transcript (NM_007294.4).

**Supplementary Figure 15. Workflow for identifying sequence alteration–associated methylation changes around structural variant breakpoints.** Somatic clonal sequence alteration events (including indels, large deletions, insertions, duplications, and translocations) were identified as variants present exclusively in engineered cell lines (OVCAR8 H4-KO, OVCAR8 A6-meth, and UWB1+BRCA1) and absent in their respective parental lines (OVCAR8 and UWB1). Methylation changes were considered for heterozygous variants supported by ⩾4 reads for both alternative (ALT) and reference (REF) alleles.

**Supplementary Figure 16. Global relationships between deletion events and *in cis* CpG methylation changes. (A)** Density plot showing CpG methylation changes relative to the nearest deletion breakpoint for small deletions (INDEL, <50 bp). Each point represents an individual CpG site. The x-axis indicates distance from the closest breakpoint (negative values, upstream; positive values, downstream), and the y-axis shows the methylation difference on the altered allele relative to the reference allele. CpGs exhibiting significant methylation loss (Fisher's exact test, adjusted p-value ⩽ 0.05 and methylation difference ⩾80%) are marked with circles; all other CpGs are shown as crosses. Point color reflects CpG density, ranging from high (yellow) to low (navy). **(B)** Proportion of deletion events associated with increased, decreased, or unchanged CpG methylation across deletion classes and models. Events are stratified by deletion type (small deletions – INDEL and large deletions (≥50 bp) classified as structural variants – SV), and by model, including events shared between OVCAR8 A6-meth and OVCAR8 H4-KO, as well as events unique to OVCAR8 A6-meth, OVCAR8 H4-KO, or UWB1. Bar colors indicate the direction of methylation change on the altered allele relative to the reference allele: increased (red), decreased (blue), or no change (grey). Event counts are shown within each bar. **(C)** Relationship between deletion length and the presence of significant CpG methylation changes for small deletions (INDEL). Deletion lengths are compared between events associated with increased, decreased, or no significant methylation change, stratified by reference allele methylation status (methylated versus unmethylated). **(D)** Relationship between deletion length and significant CpG methylation changes for large deletions (SV), shown using the same stratification and color scheme as in panel C. Statistical significance of differences in deletion length distributions was assessed using a two-tailed Wilcoxon signed-rank test.

**Supplementary Figure 17. Global relationships between insertions and *in cis* CpG methylation changes.** Density plots showing CpG methylation changes relative to the nearest breakpoint for **(A)** small insertions (INS-INDEL, <50 bp); and **(B)** large insertions classified as structural variants (INS-SV, ⩾50 bp). In each panel, each point represents an individual CpG site. The x-axis indicates the distance from the closest breakpoint (negative values, upstream; positive values, downstream), and the y-axis shows the methylation difference on the altered allele relative to the reference allele. CpGs exhibiting significant methylation change (Fisher's exact test, adjusted p-value ⩽ 0.05 and absolute methylation difference ⩾ 80%) are marked with circles, while all other CpGs are shown as crosses. Point color reflects CpG density, ranging from high (yellow) to low (navy). **(C)** Number of insertion events across models, stratified by events shared between OVCAR8 A6-meth and OVCAR8 H4-KO, and events unique to OVCAR8 A6-meth, OVCAR8 H4-KO, or UWB1. Bar colors indicate the direction of methylation change on the altered allele relative to the reference allele: increased (red), decreased (blue), or no change (grey). Event counts are shown within each bar. **(D-E)** Relationship between insertion length and the presence of significant CpG methylation changes for small insertions (INDEL INS) **(D)** and large insertions (SV INS) **(E)**, stratified by reference allele methylation status (methylated versus unmethylated).

**Supplementary Figure 18. Global relationships between duplications, translocations and *in cis* CpG methylation changes. (A)** Density plots showing CpG methylation changes relative to the nearest breakpoint for duplication. Each point represents an individual CpG site. The x-axis indicates the distance from the closest breakpoint (negative values, upstream; positive values, downstream), and the y-axis shows the methylation difference on the altered allele relative to the reference allele. CpGs exhibiting significant methylation change (Fisher's exact test, $P \leqslant 0.05$ and absolute methylation difference ⩾80%) are marked with circles, while all other CpGs are shown as crosses. Point color reflects CpG density, ranging from high (yellow) to low (navy). **(B)** Number of duplication events across models, stratified by events shared between OVCAR8 A6-meth and OVCAR8 H4-KO, and events unique to OVCAR8 A6-meth, OVCAR8 H4-KO, or UWB1. Bar colors indicate the direction of methylation change on the altered allele relative to the reference allele: increased (red), decreased (blue), or no change (grey). Event counts are shown within each bar. **(C)** Relationship between length of duplications and the presence of significant CpG methylation changes, stratified by reference allele methylation status (methylated versus unmethylated). **(D)** Number of translocation events across models, shown using the same stratification and color scheme as in panel (**B)**.

**Supplementary Figure 19. Microhomology usage is not associated with *in cis* CpG methylation changes following sequence deletions.** Heatmaps summarizing microhomology patterns surrounding deletion (SV and indel) breakpoints for OVCAR8-derived **(A)** events associated with significant CpG methylation change and **(B)** events without methylation change. Heatmaps represent aggregated base-matching frequencies between the 3′ and 5′ flanking sequences at deletion junctions, normalized to the maximum observed value. Color intensity reflects the relative frequency of base matches, with higher values (red) indicating stronger microhomology. **(C)** Distribution of deletion repair categories stratified by methylation outcome and sample type (OVCAR8 versus UWB1). Deletions are classified as theta-mediated end joining (TMEJ; deletions ⩾4 bp with ⩾2 bp microhomology) or non-microhomology deletions. Bars indicate the proportion of events with or without associated CpG methylation change. No significant enrichment of microhomology-mediated repair is observed among deletions associated with methylation change ($\chi^2$ test shown).

**Supplementary Figure 20. Strand-specific promoter demethylation following CRISPR-induced deletions in OVCAR8 A6-meth cells.** Frequency of CpG methylation alleles across the *BRCA1* promoter region in parental OVCAR8 A6-meth **(A)** and following CRISPR editing of OVCAR8 A6-meth cells using guide RNA #3, performed without any clonal selection **(B)**. **(C)** Allele-resolved CpG methylation and corresponding genomic alterations across the *BRCA1* promoter following CRISPR editing of OVCAR8 A6-meth cells using guide RNA #3 by adaptive Nanopore direct DNA sequencing. Reads are stratified by deletion size (⩽2–3 bp versus >10 bp) and by alignment strand relative to the reference genome (positive or negative strand), as indicated. Each dot represents a CpG site, colored by methylation status (methylated, red; unmethylated, blue). The *BRCA1* promoter region is highlighted in grey. Black blocks indicate the genomic extent of CRISPR-induced deletions.

**Supplementary Figure 21. Olaparib selection enriched for loss of *BRCA1* promoter methylation in OVCAR8 A6-meth cells.** Distribution of *BRCA1* promoter epialleles across OVCAR8 A6-meth parental and CRISPR-edited cell populations following X days of olaparib selection. Each bar represents the proportion of sequencing reads classified as fully methylated (red) or fully unmethylated (blue). Numbers within each bar indicate the absolute read counts for each epiallele class.

**Supplementary Figure 22. Dose–response analysis of parental and CRISPR-edited cells treated with olaparib and cisplatin**. Olaparib selected cells (parental (blue), guide #1 (orange), and guide #2 (green)) were treated with increasing concentrations of olaparib (top panel) or cisplatin (bottom panel), and survival (%) was measured relative to untreated controls using CTG assay. Data represents mean ± error bars (3 independent replicates). For cisplatin, dose–response curves were fitted using a four-parameter log-logistic model, and guide #1 (IC50:1.52μM) and guide #2 (IC50:1.3μM) exhibited significantly increased resistance compared to parental cells (IC50:0.71μM), as indicated by higher IC50 values (guide #1 vs parental, $p < 0.001$; guide #2 vs parental, $p = 0.0046$), while no significant difference was observed between guide #1 and guide #2. For olaparib, no clear dose–response relationship was observed across the full concentration range; therefore, IC50 could not be reliably determined. However, at higher concentrations (⩾2.5 μM), guide #1 and guide #2 exhibited significantly higher survival compared to parental cells ($p < 0.001$; linear model with interaction), consistent with increased resistance. No significant differences were observed between guide #1 and guide #2.

### Supplementary Figure 1

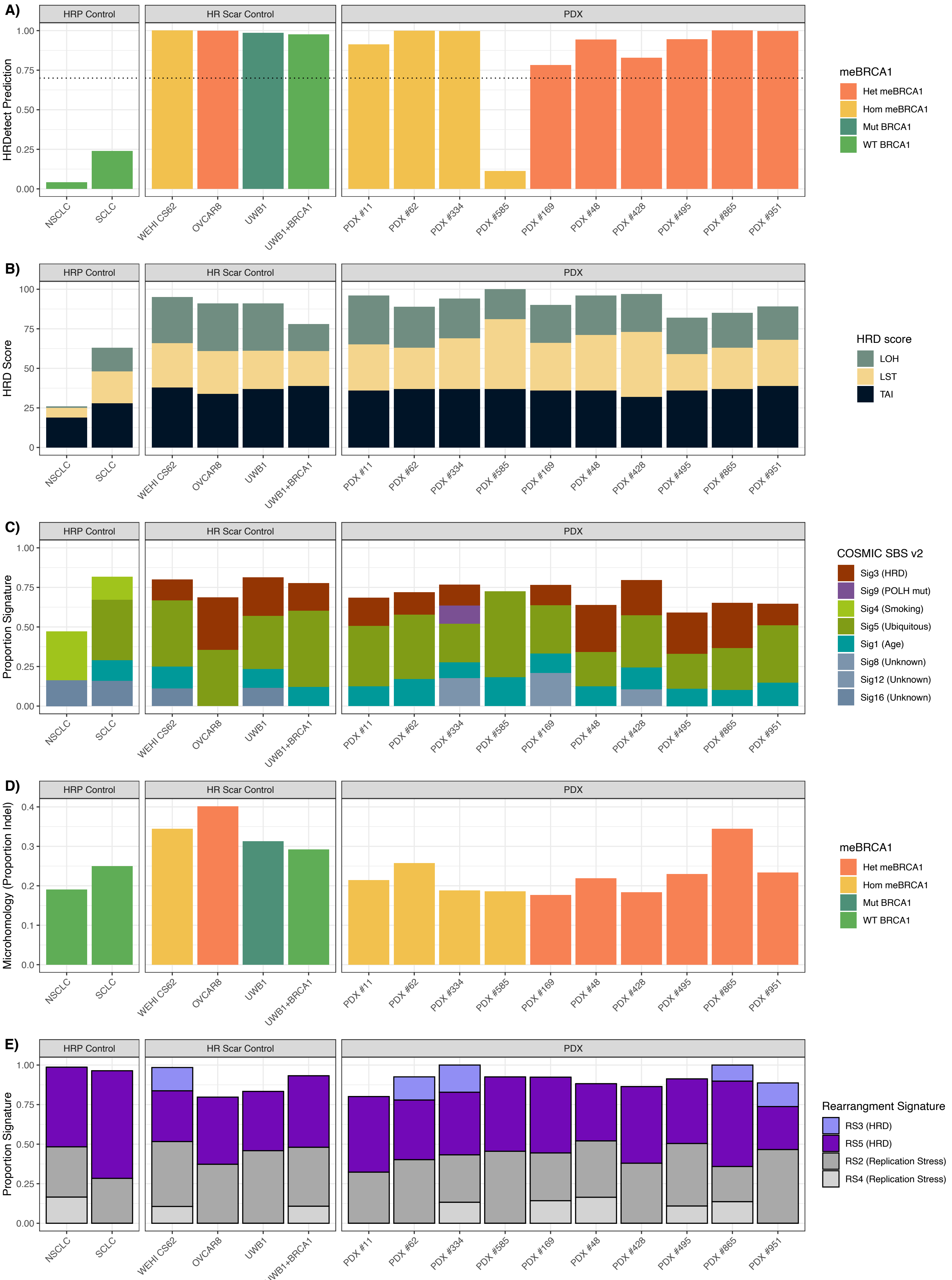

#### Supplementary Figure 2

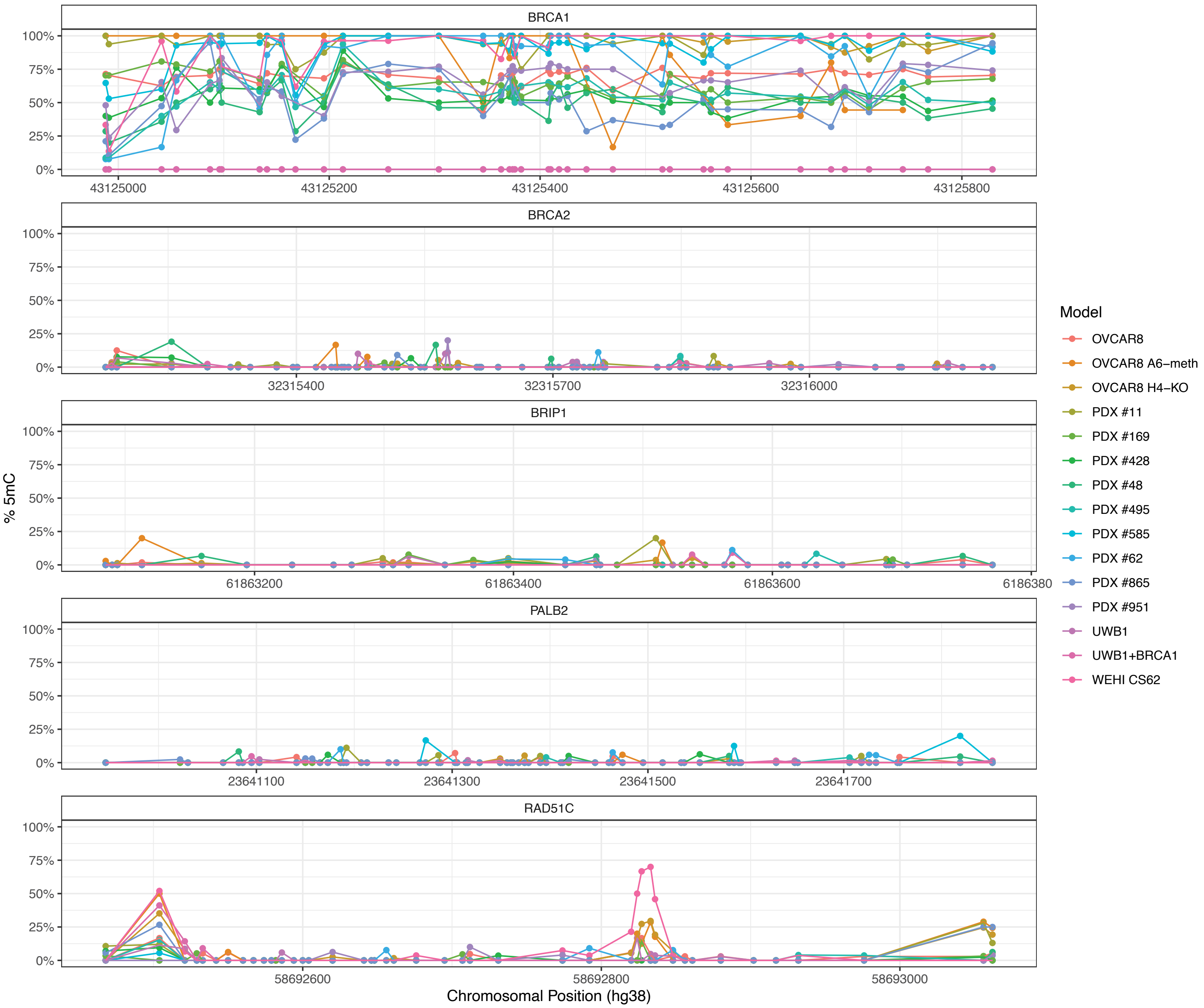

### Supplementary Figure 3

A)

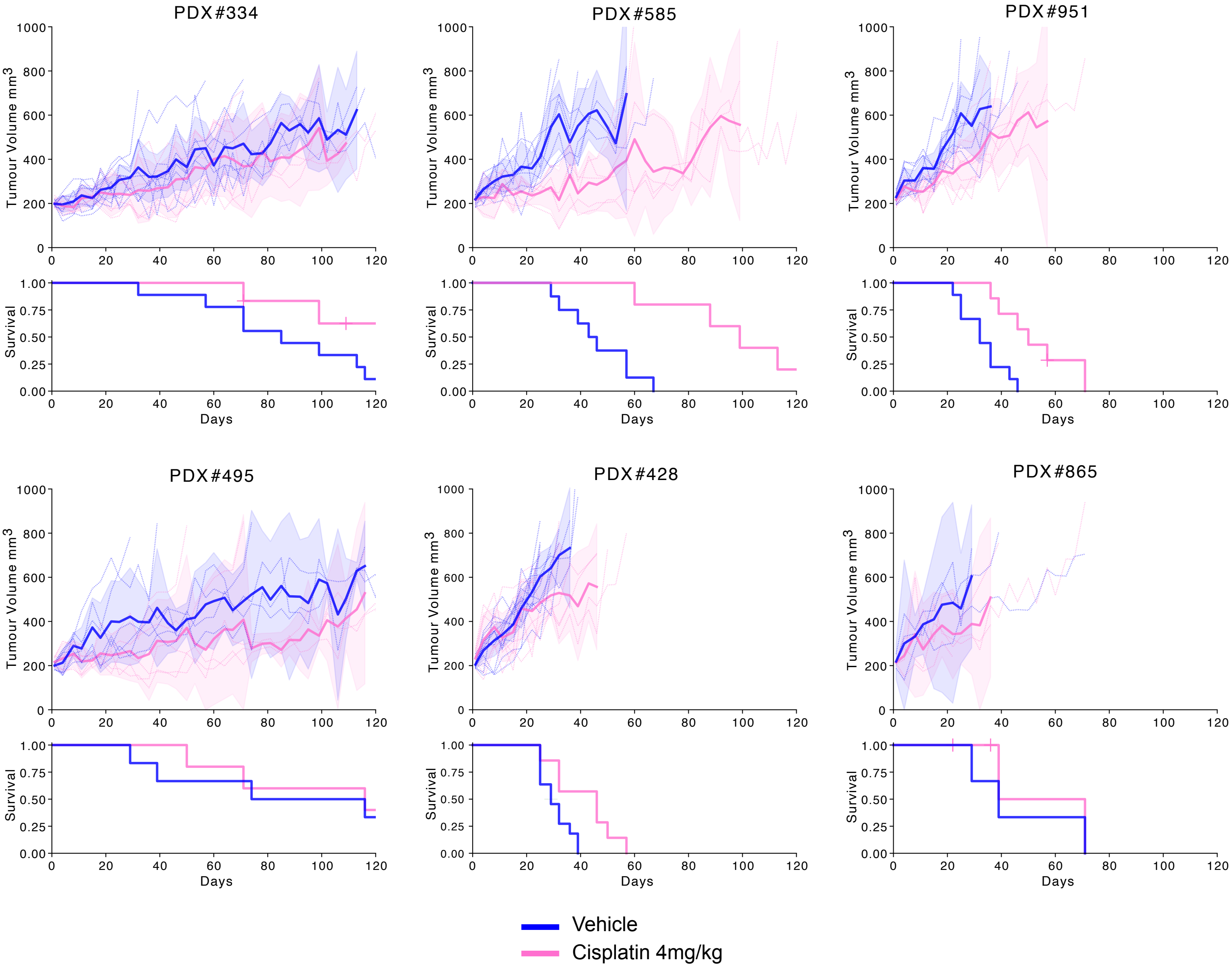


B)

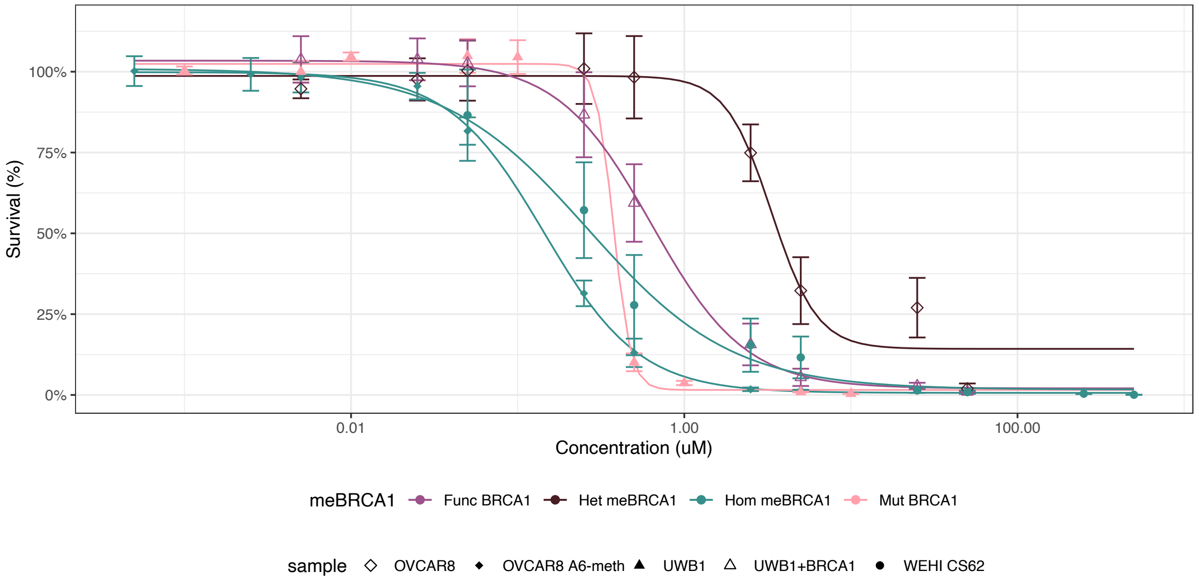

#### Supplementary Figure 4

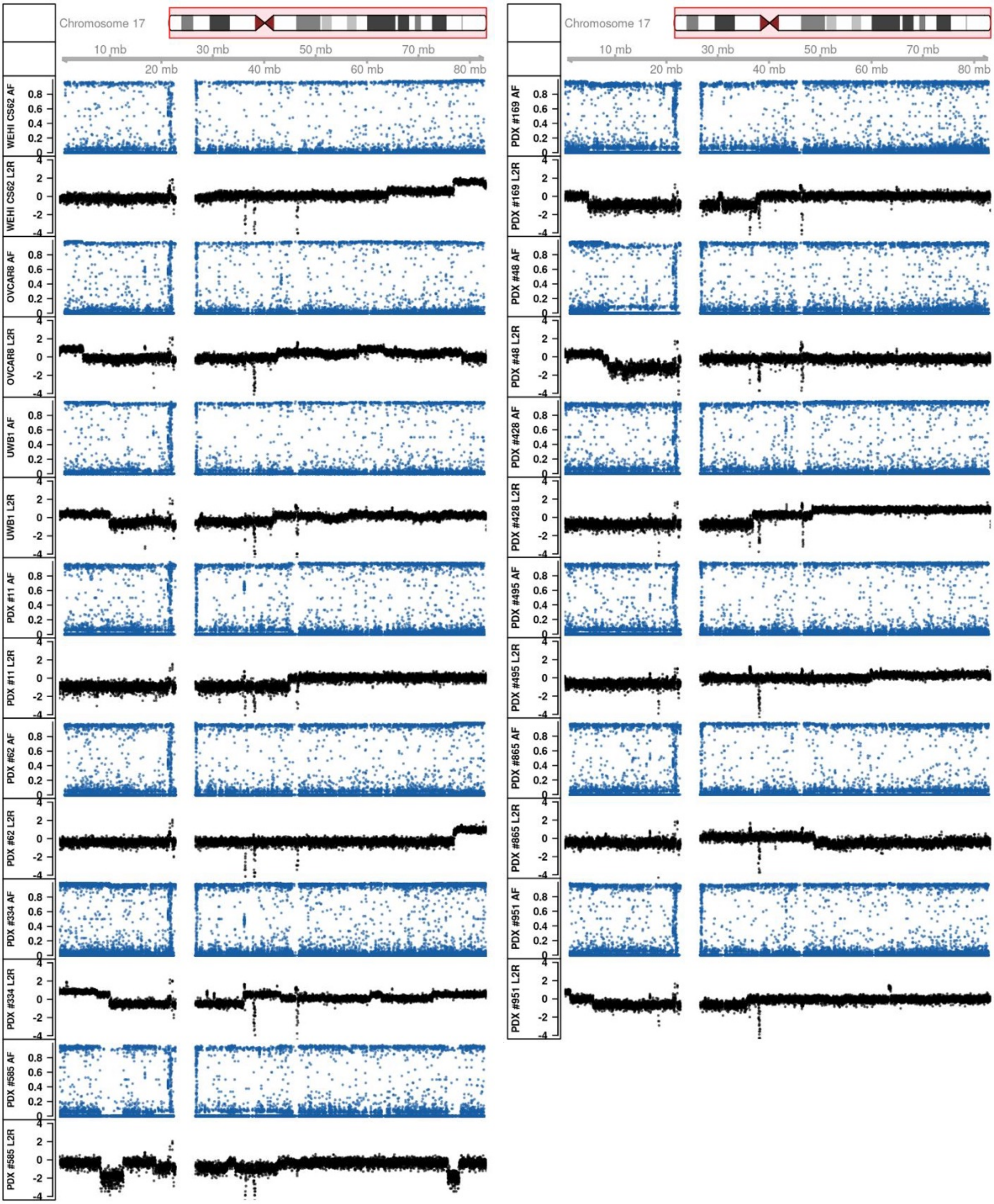

### Supplementary Figure 5

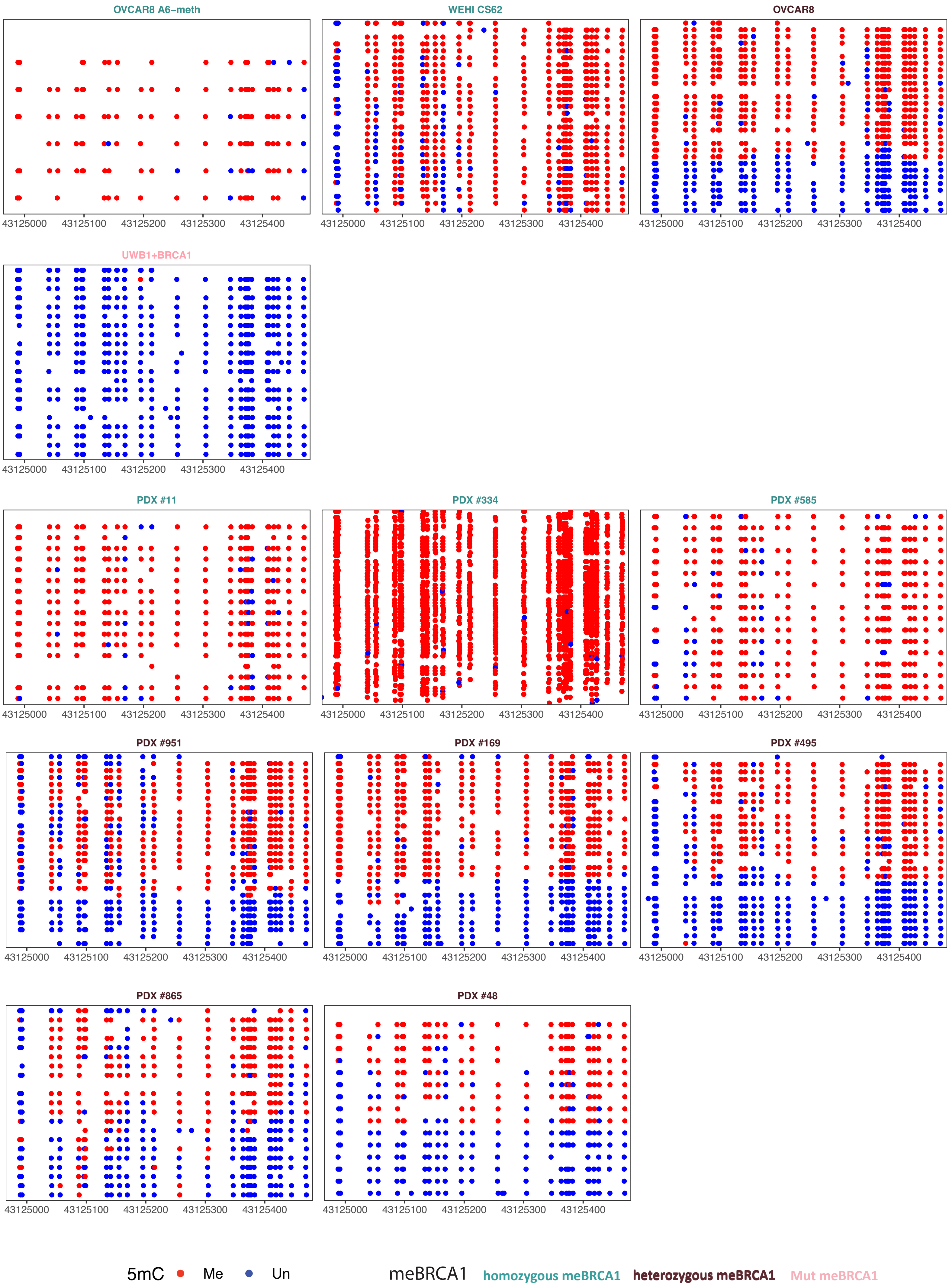

#### Supplementary Figure 6

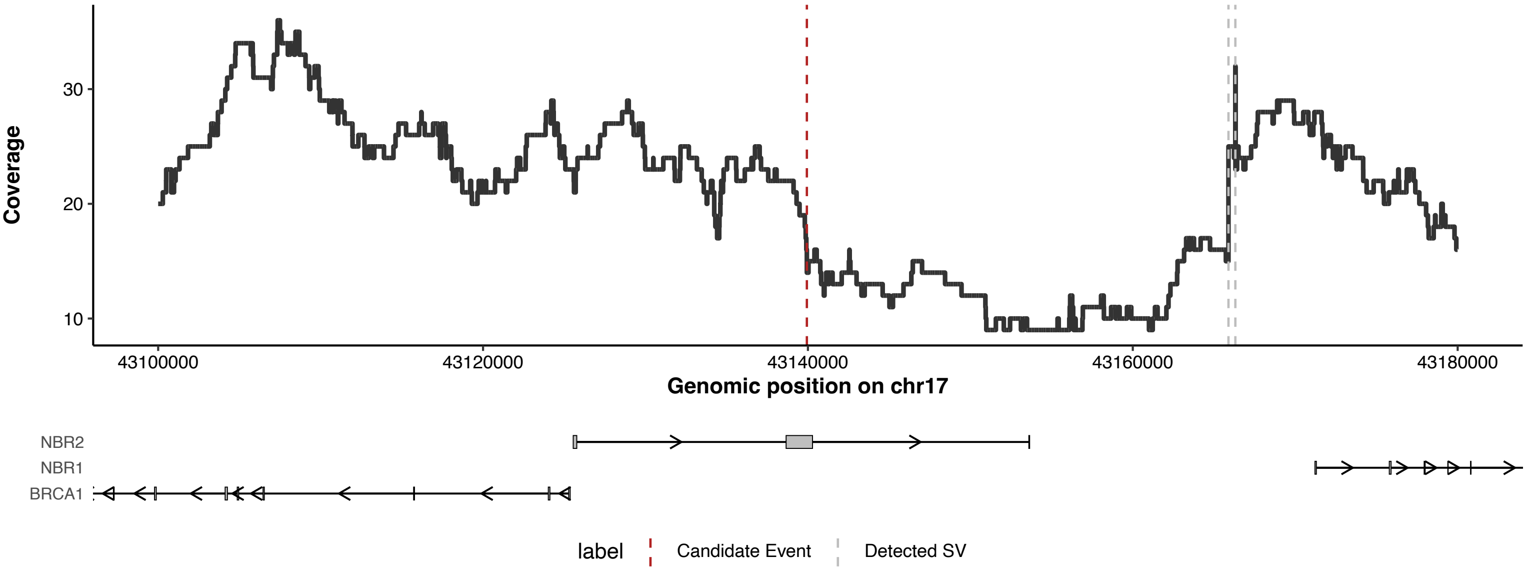

**Supplementary Figure 7**

A)

B)

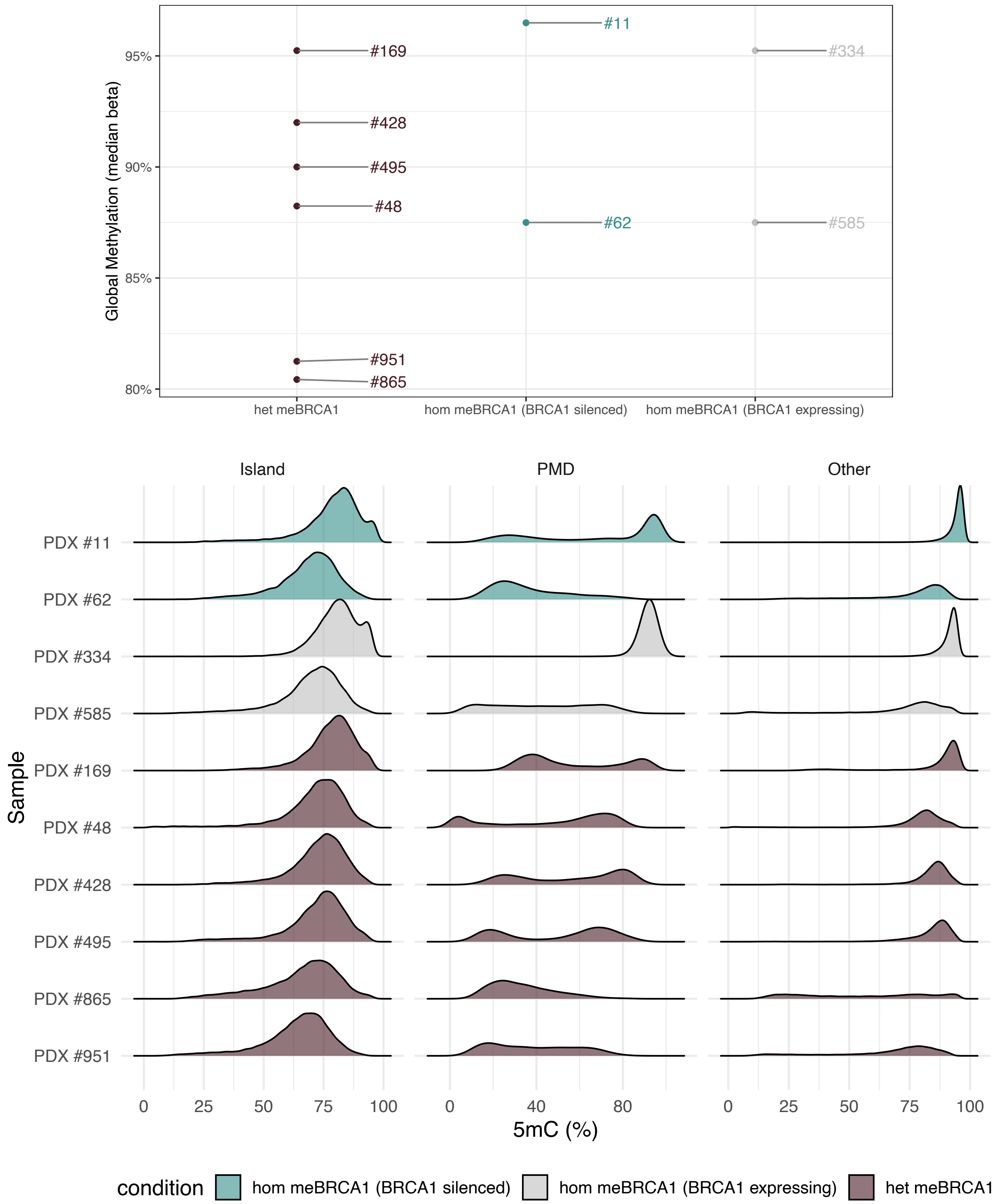

### Supplementary Figure 8

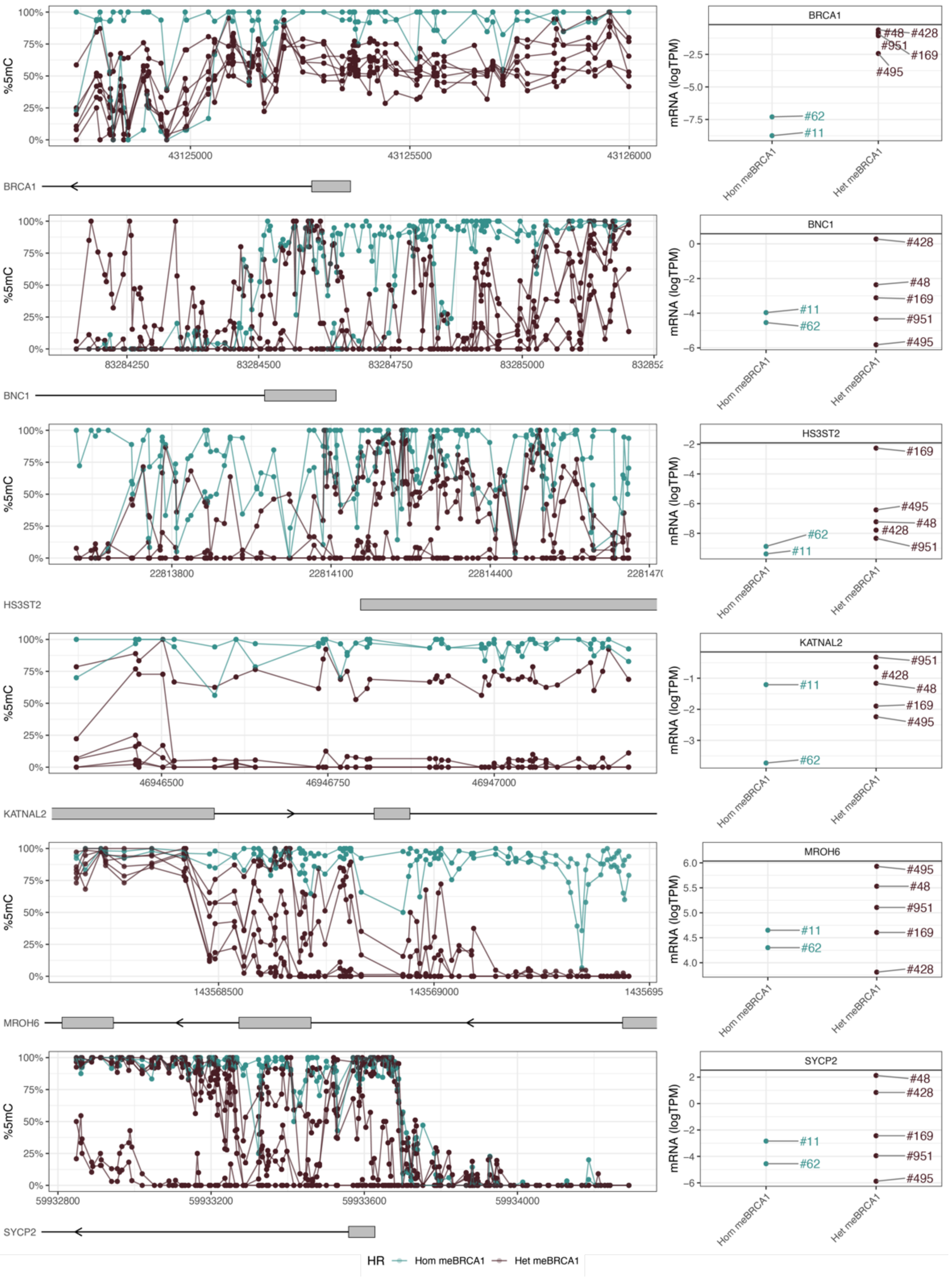

### Supplementary Figure 9

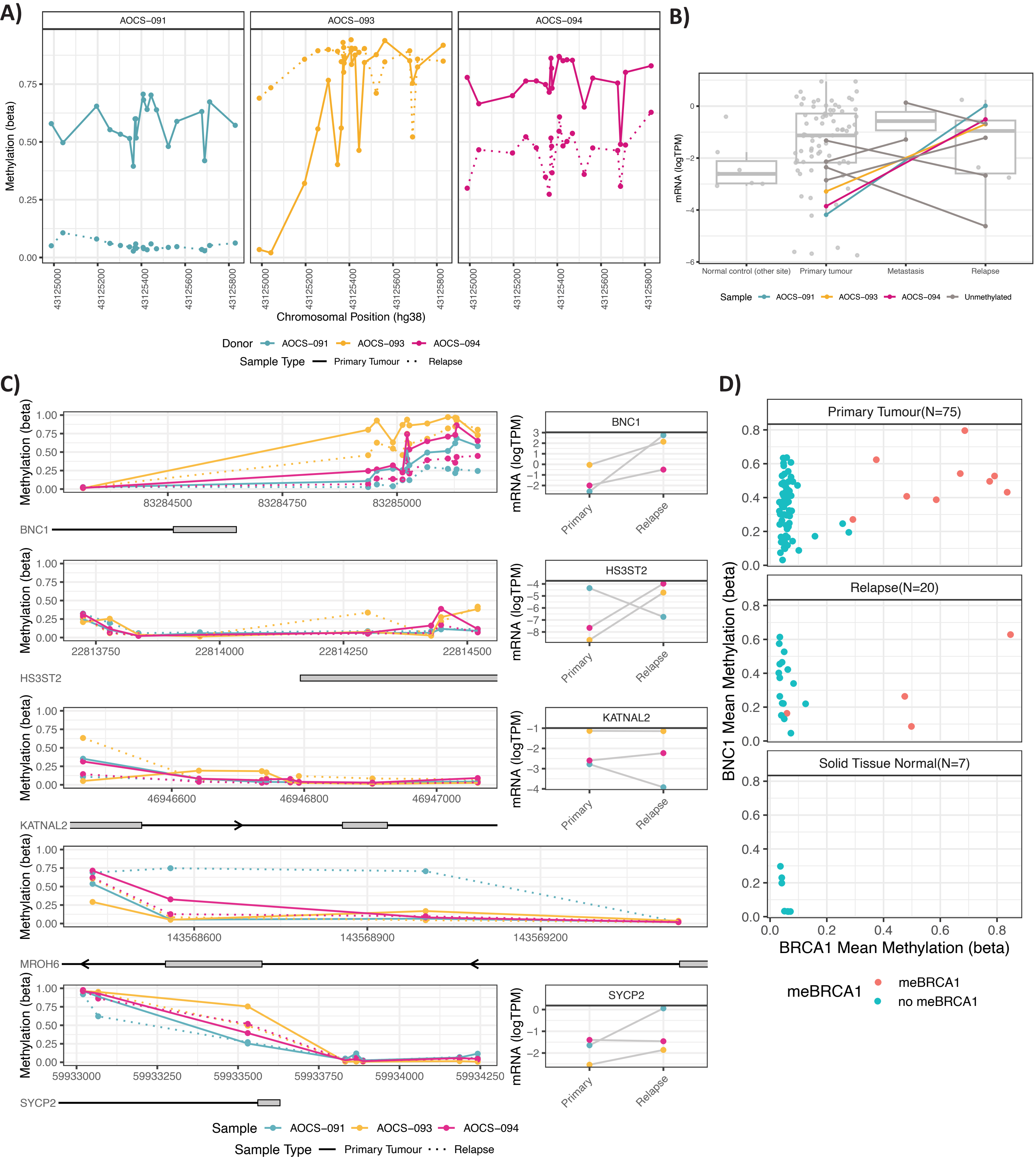

**Supplementary Figure 10**

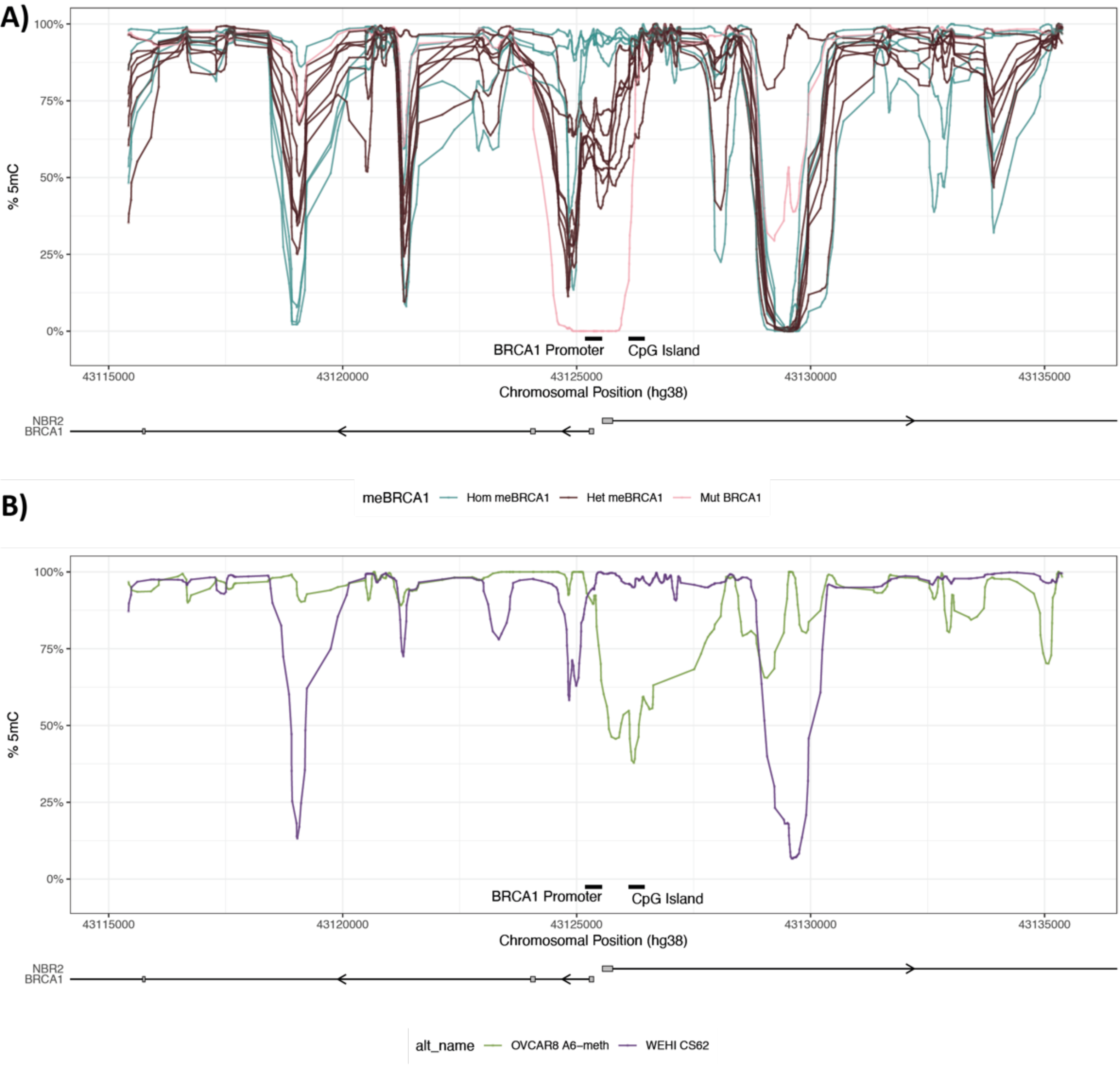

### Supplementary Figure 11

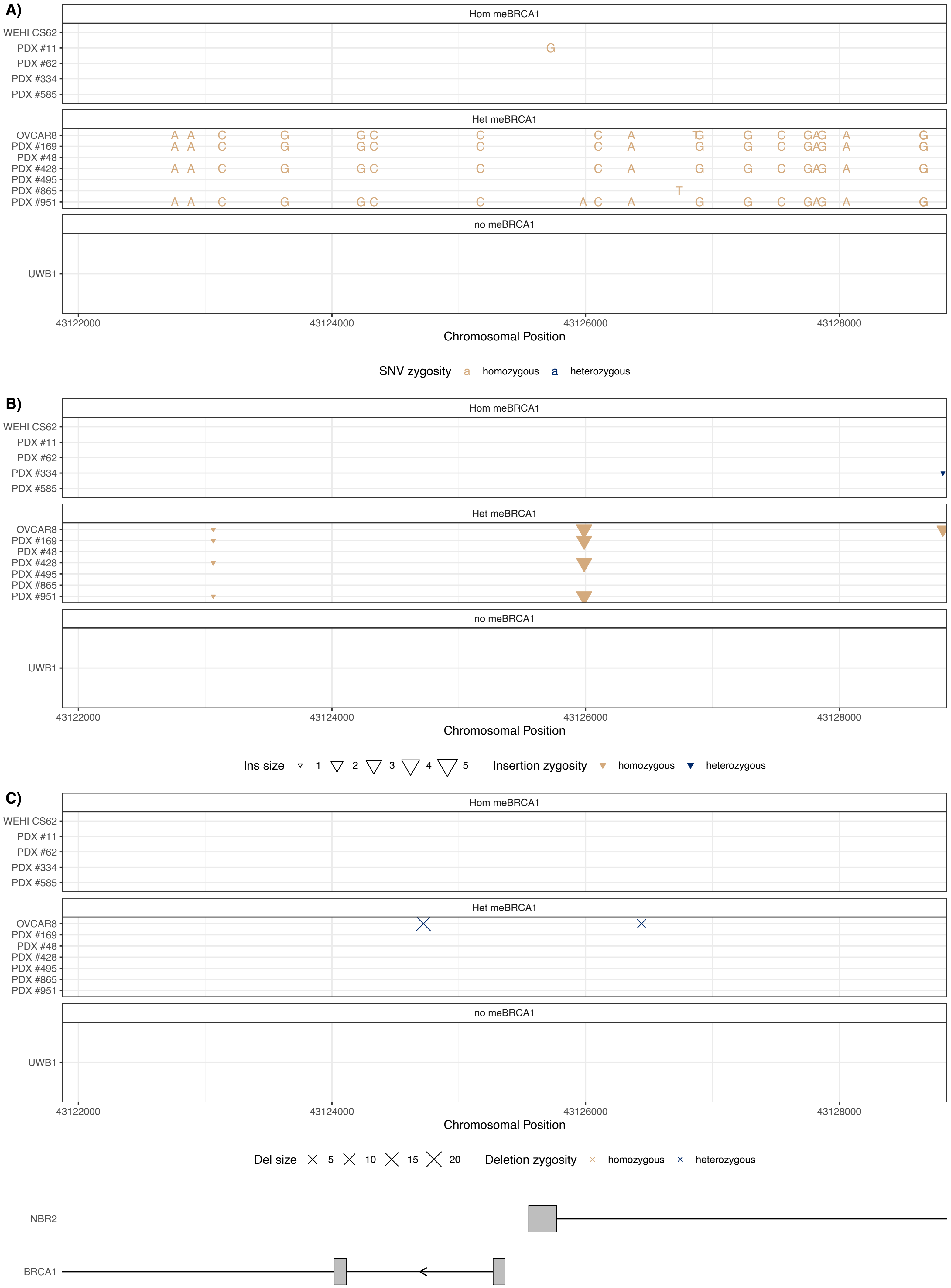

**Supplementary Figure 12**

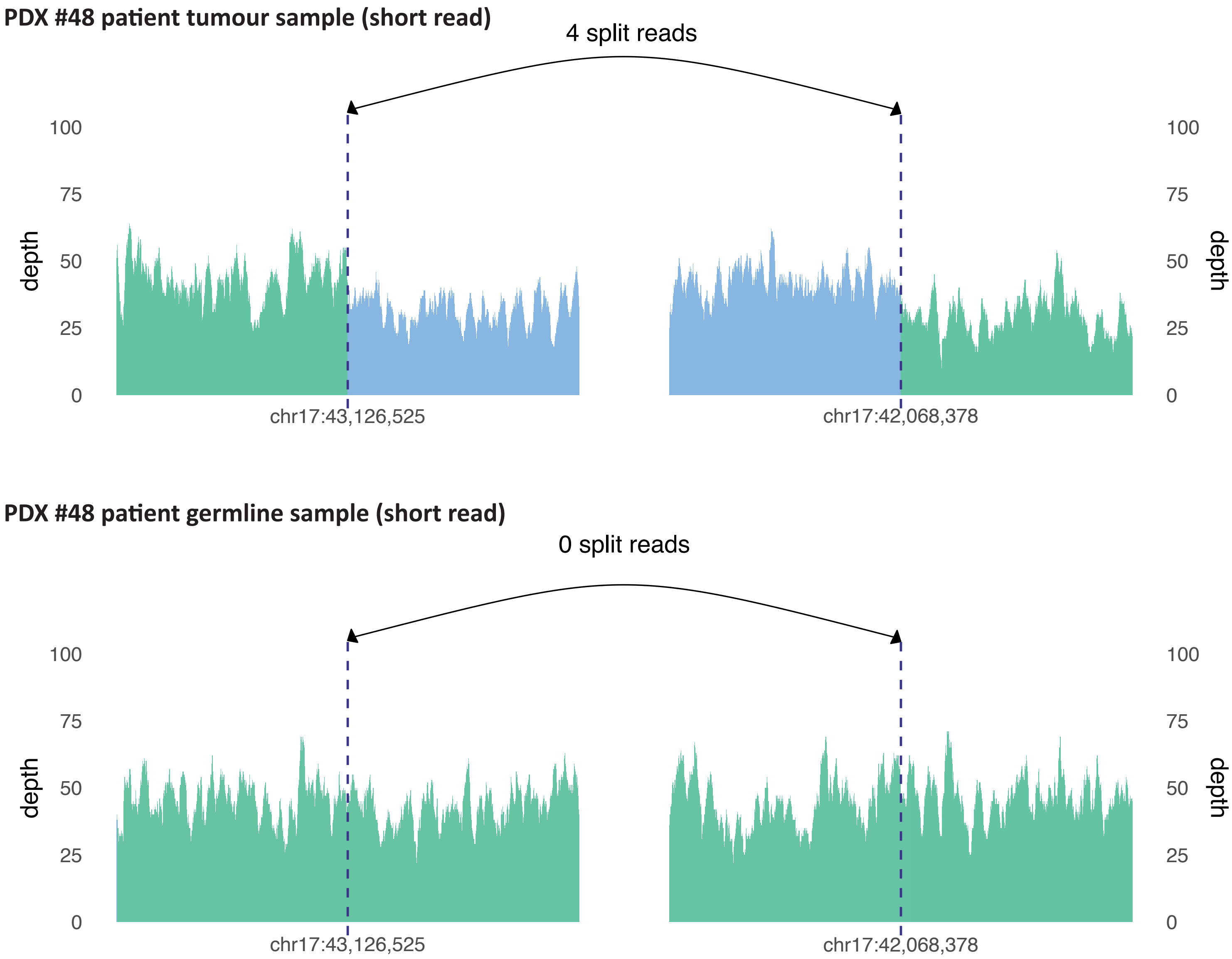

**Supplementary Figure 13**

#### Igg

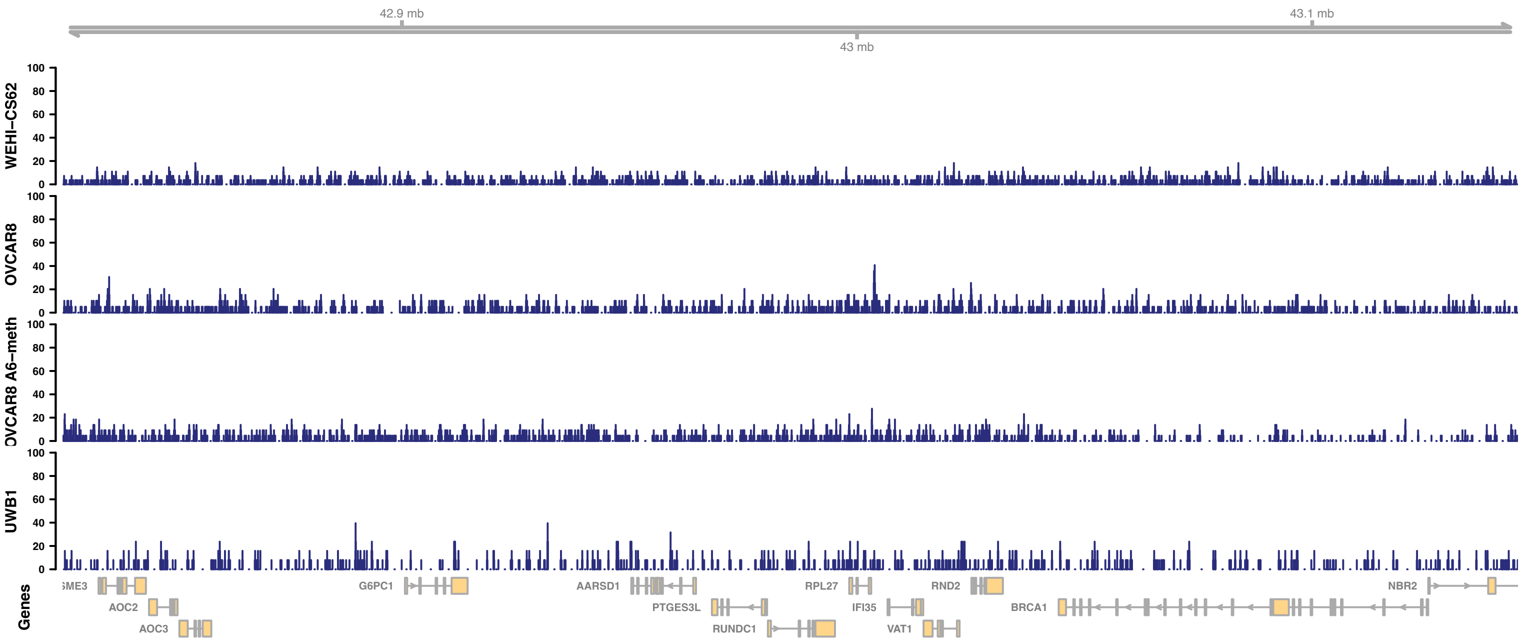


## H3K27me3

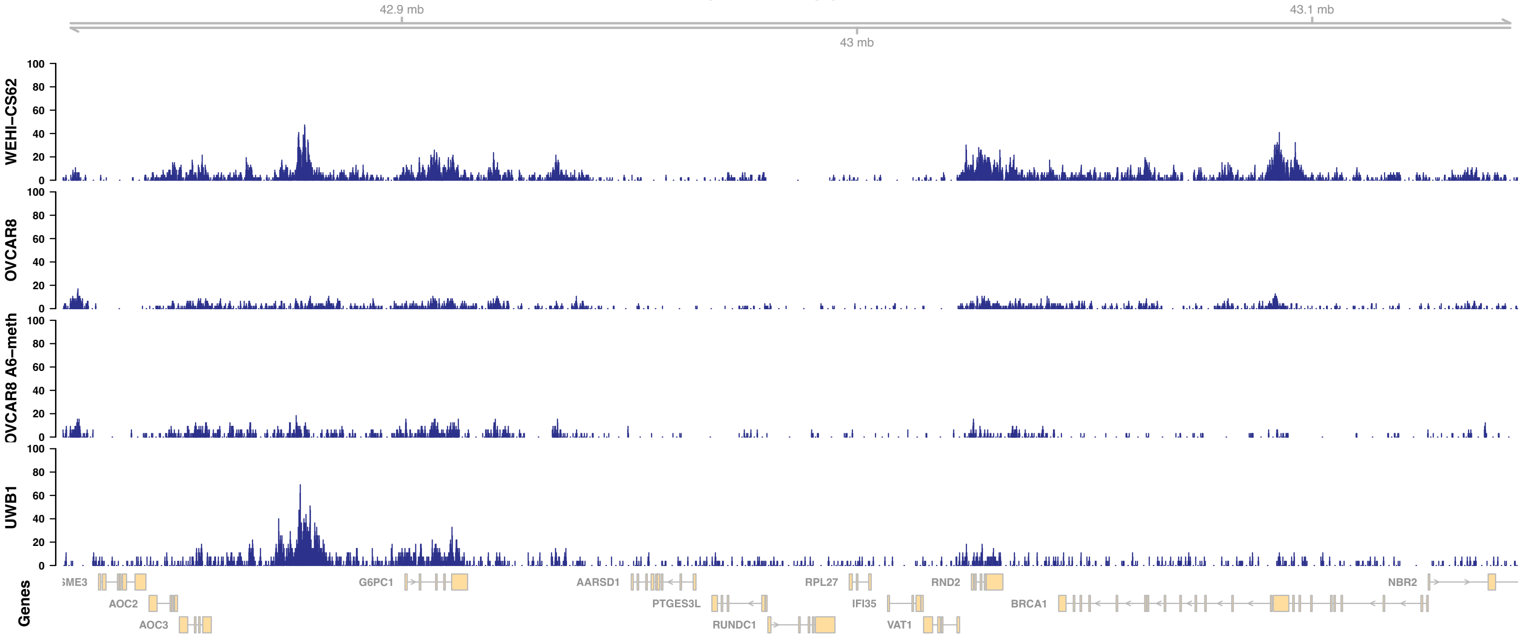


## H3K9me3

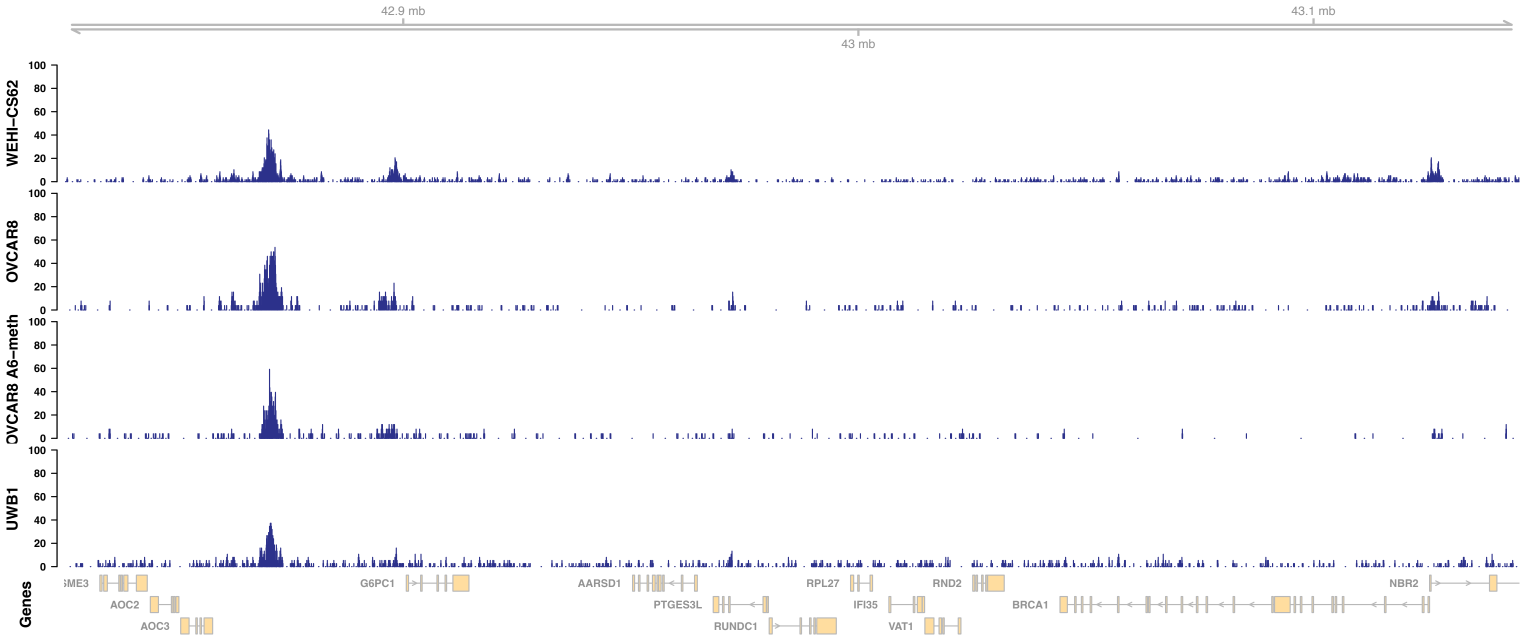

### Supplementary Figure 14

## H3K4me1

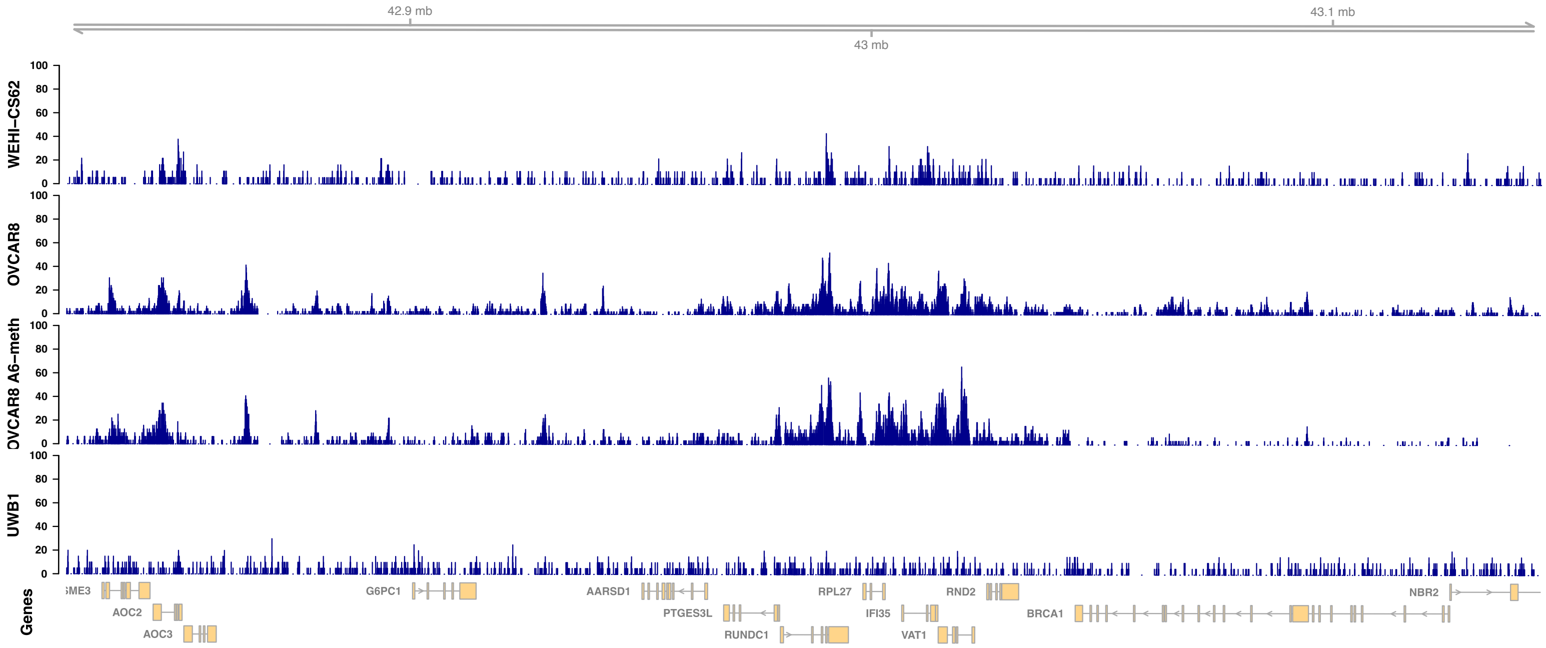


## H3K4me3

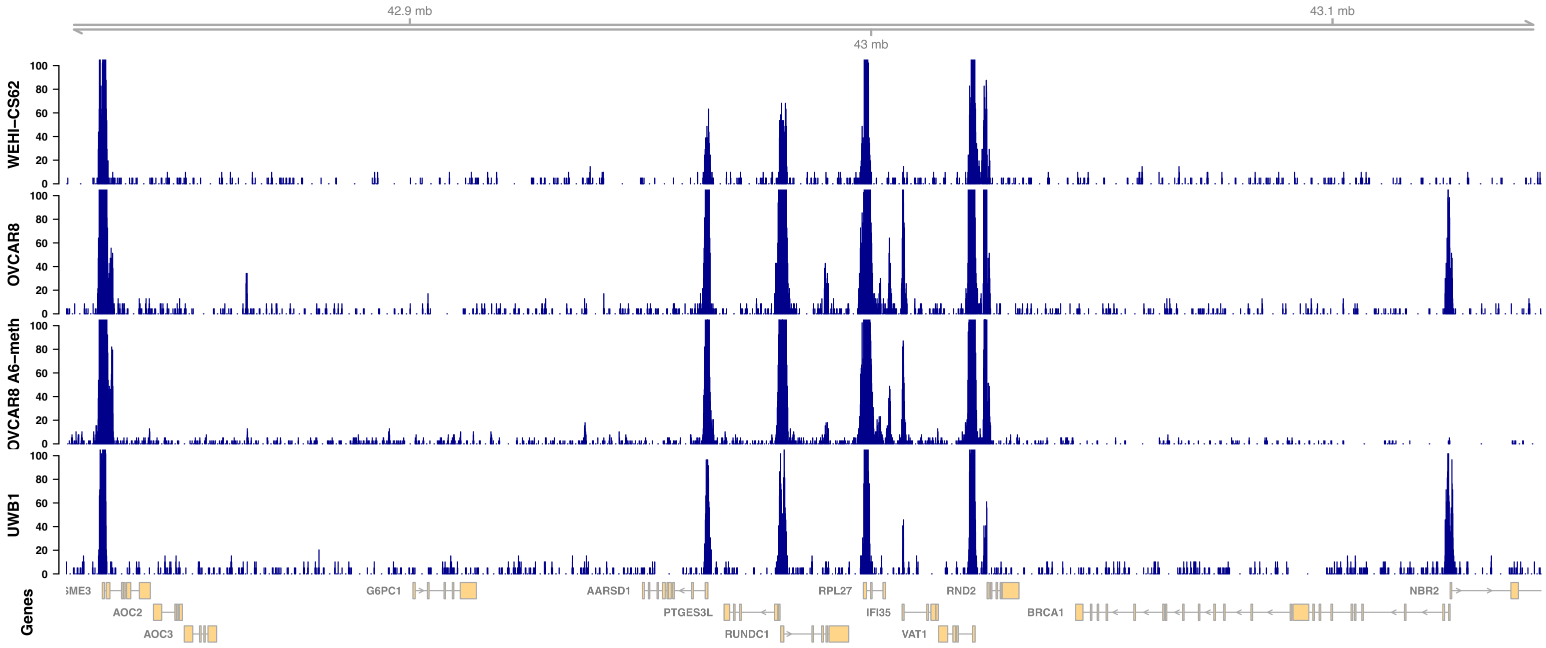

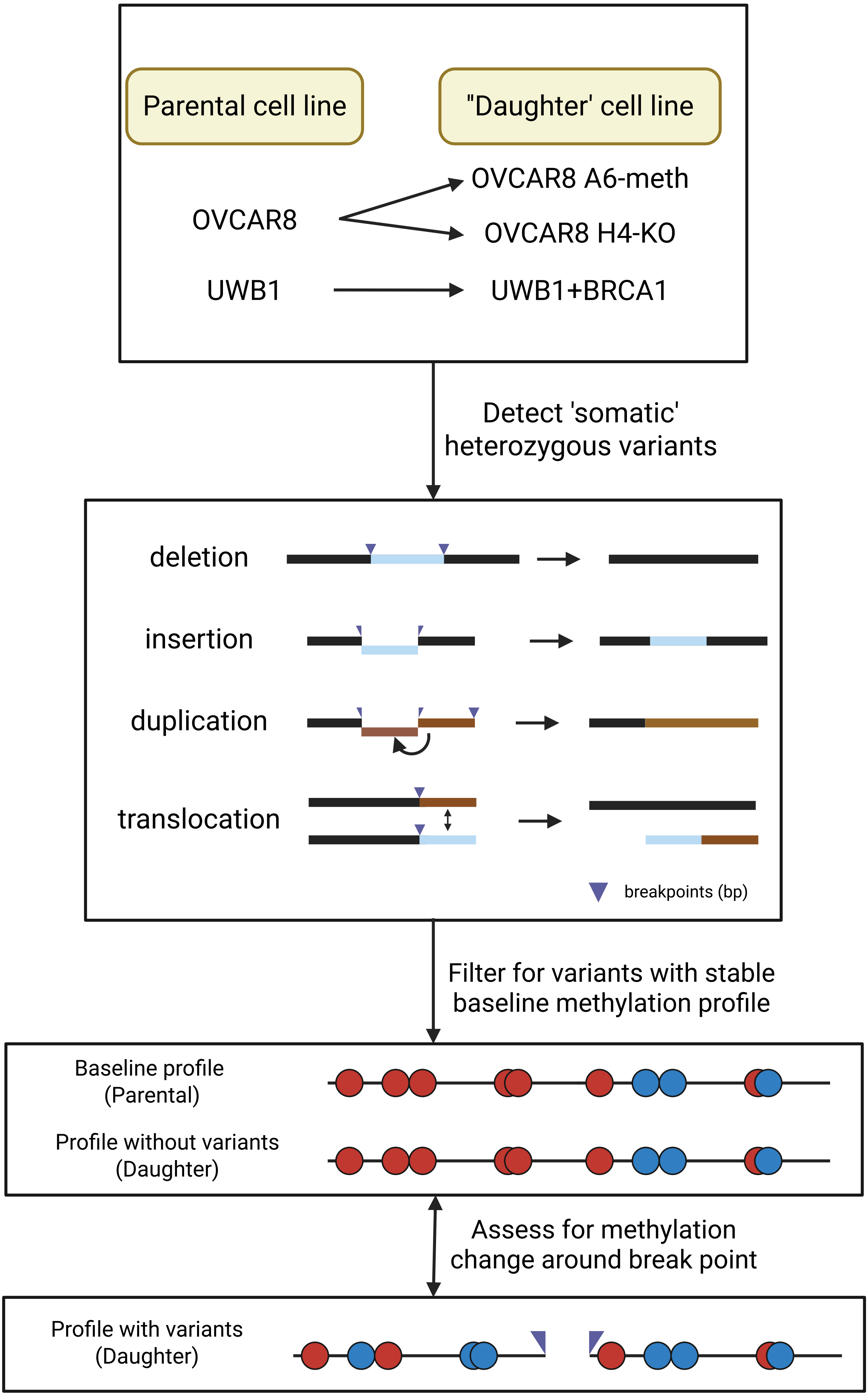
Parental cell line
"Daughter' cell line
OVCAR8
OVCAR8 A6-meth
OVCAR8 H4-KO
UWB1
UWB1+BRCA1
Detect 'somatic' heterozygous variants
deletion
insertion
duplication
translocation
breakpoints (bp)
Filter for variants with stable baseline methylation profile
Baseline profile (Parental)
Profile without variants (Daughter)
Assess for methylation change around break point
Profile with variants (Daughter)

**Supplementary Figure 16**

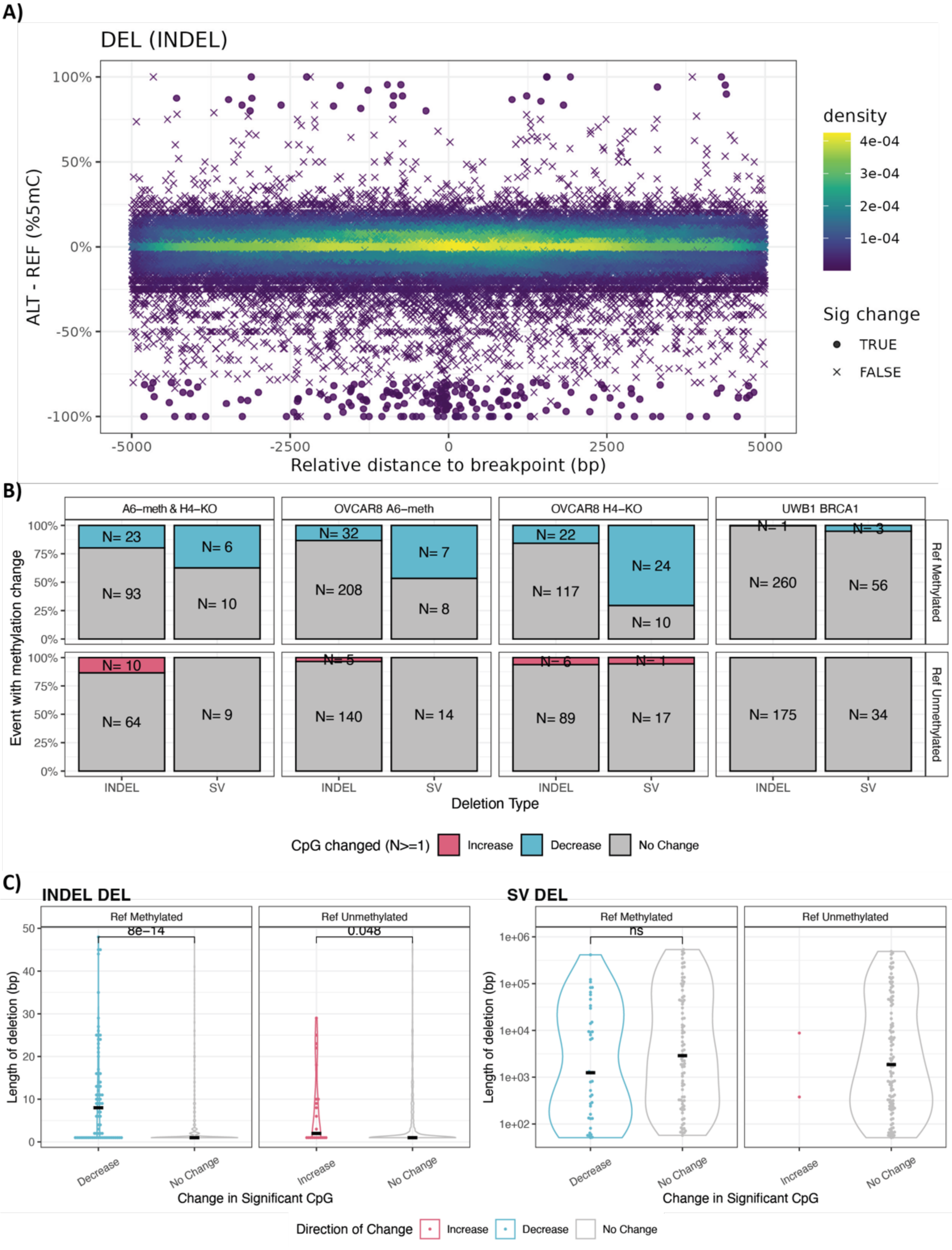

**Supplementary Figure 17**

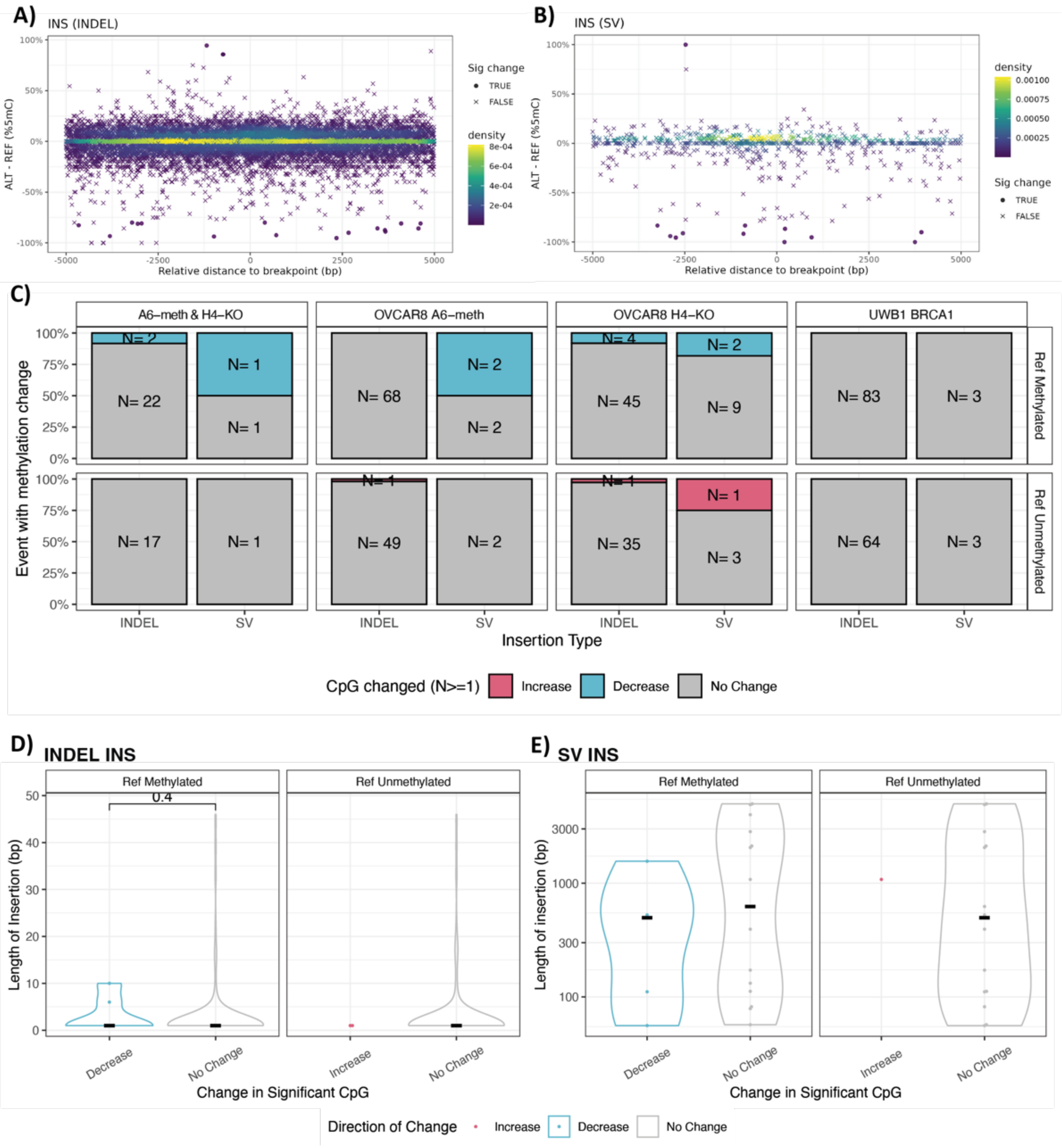

**Supplementary Figure 18**

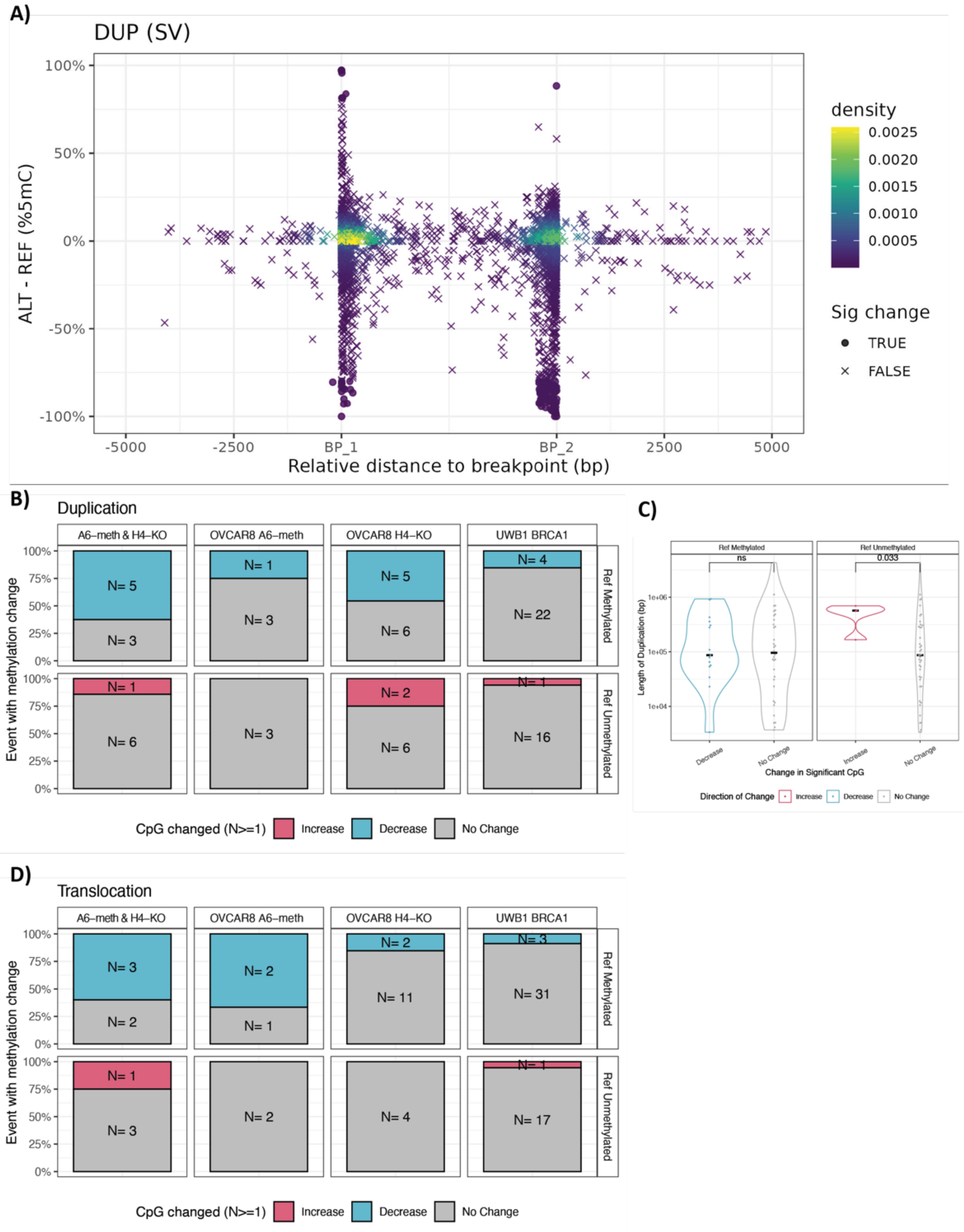

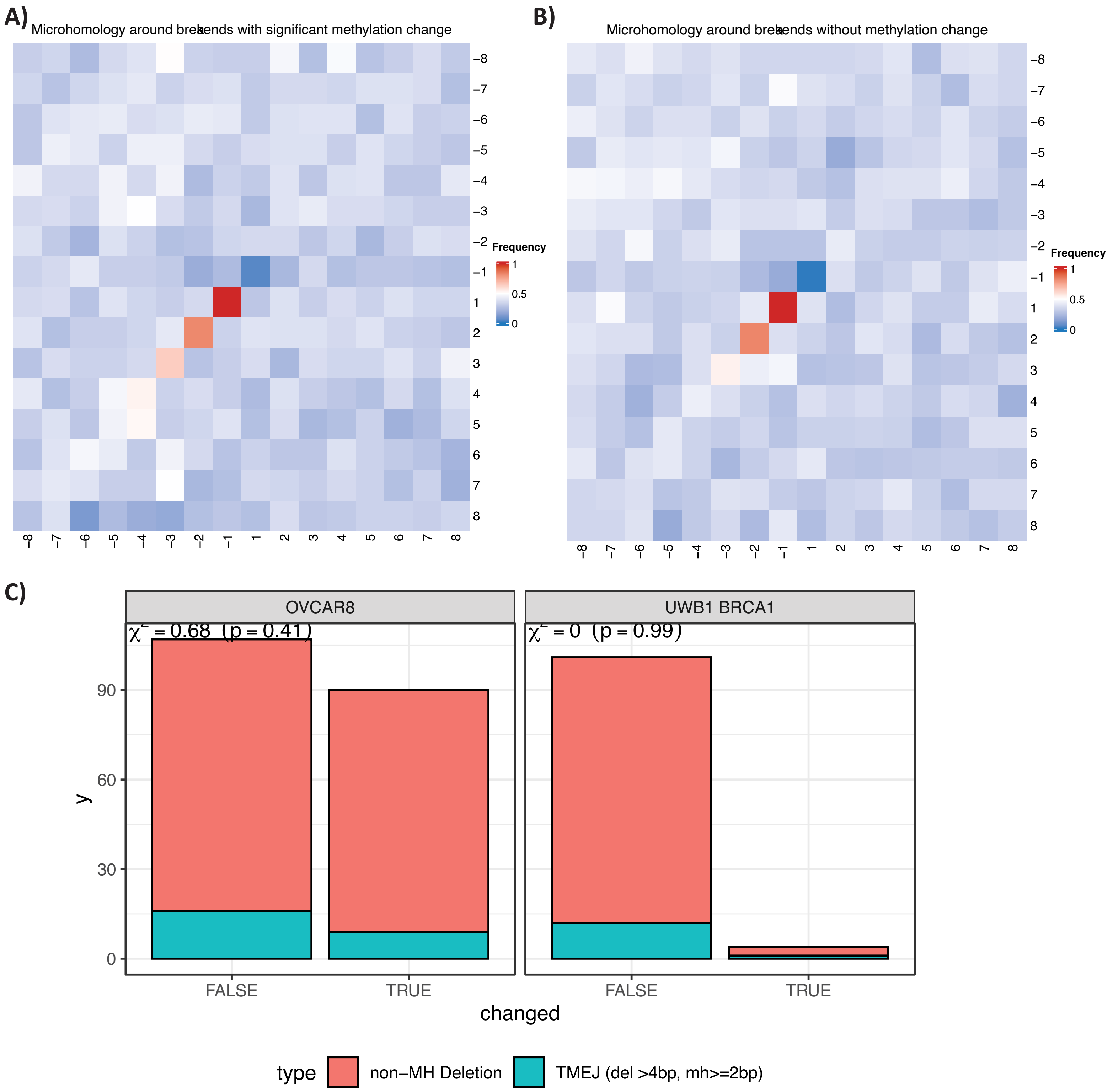
A)
Microhomology around breakends with significant methylation change
Frequency
1
0.5
0
−8
−7
−6
−5
−4
−3
−2
−1
1
2
3
4
5
6
7
8
B)
Microhomology around breakends without methylation change
C)
OVCAR8
UWB1 BRCA1
χ² = 0.68 (p = 0.41)
χ² = 0 (p = 0.99)
y
0
30
60
90
FALSE
TRUE
changed
type
non−MH Deletion
TMEJ (del >4bp, mh>=2bp)

### Supplementary Figure 20

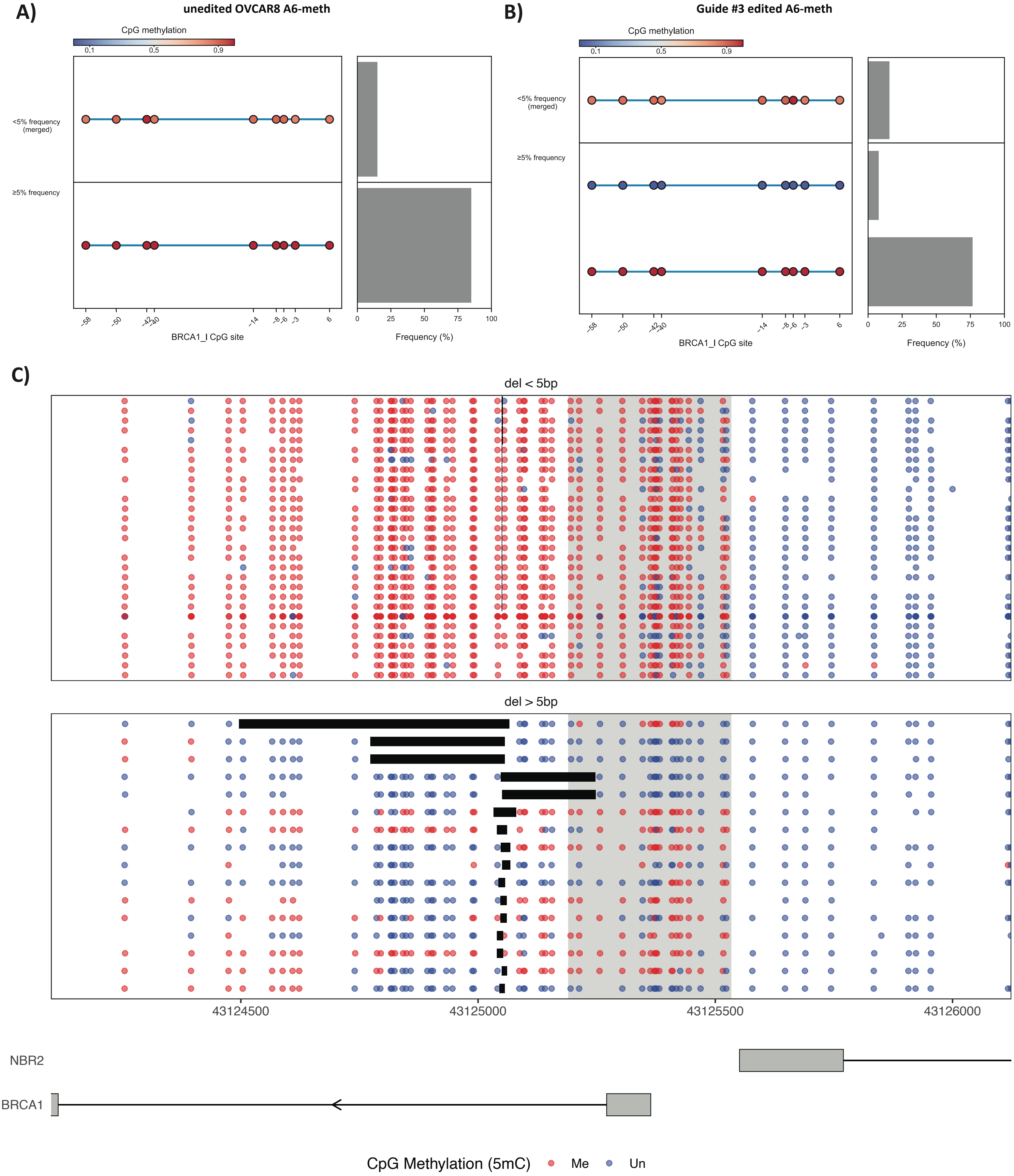

### Supplementary Figure 21

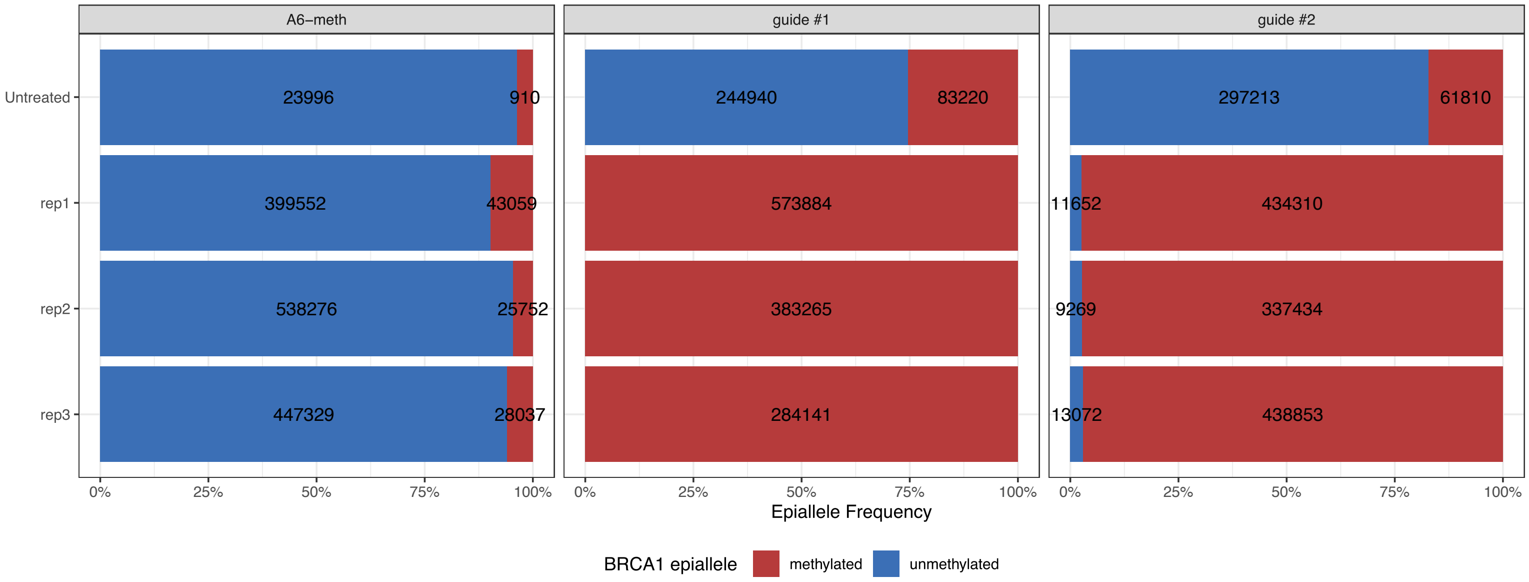

**Supplementary Figure 22**

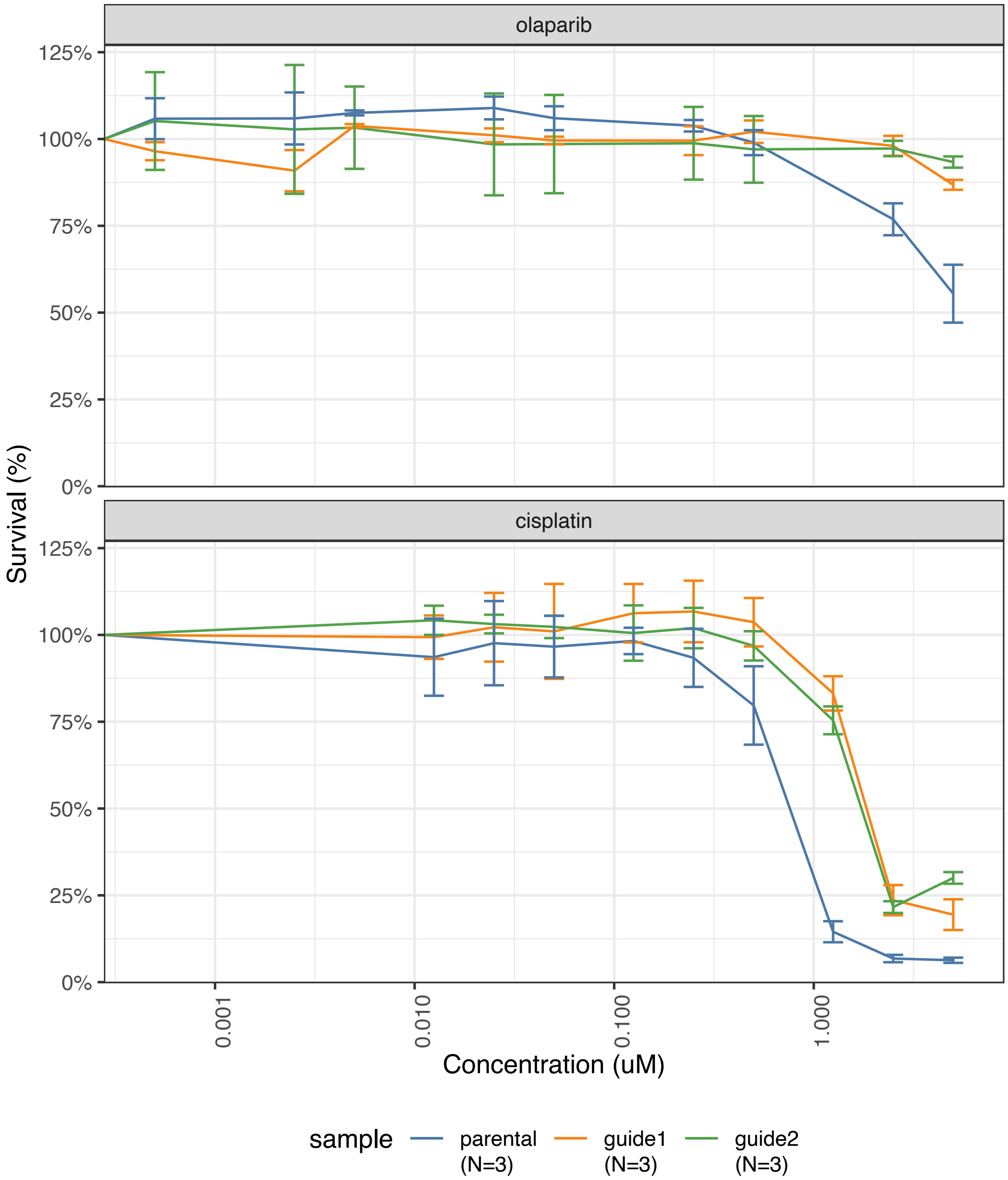
