## Supplementary Material for "Genomic alterations enable *BRCA1* methylation loss and promoter bypass to drive resistance in high-grade serous ovarian cancer"

### 1. Assessment of Batch Effects in RNA Sequencing of PDX Models

Due to the nature of tissue acquisition, RNA sequencing of patient-derived xenograft (PDX) samples was performed in multiple batches. To evaluate potential batch effects arising from library preparation, two sequencing batches were analysed:

- Batch 1: PDX #11, PDX #48, PDX #62, PDX #169, PDX #428, PDX #495, PDX #585 and PDX #951
- Batch 2: PDX #585, PDX #495 and PDX #334

Importantly, PDX #585 and PDX #495 were included in both batches using the same extracted RNA, while library preparation was performed independently for each batch, enabling direct assessment of batch-related technical variation.

Principal component analysis (PCA) across the top four dimensions demonstrated no clear separation of samples by batch, including for the duplicated PDX #585 and PDX #495 samples, indicating minimal global batch effects (Figure 1).

Given that downstream analyses focused specifically on *BRCA1* and *NBR2* expression, we further assessed the impact of batch on transcript-level detection. No significant differences were observed between batches for either gene (Figure 2), supporting the robustness of expression measurements.

For all analyses presented in the main text, expression values were calculated as the average across batches where duplicate samples were available.

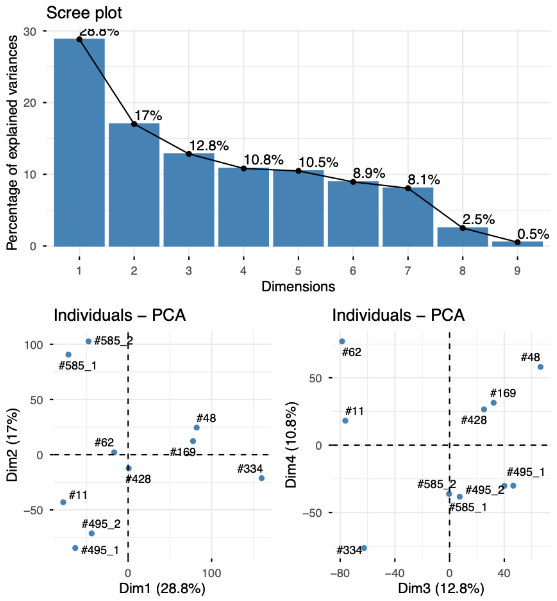


Figure 1. Assessment of batch effects in RNA sequencing of PDX models.

Top panel: Scree plot showing the percentage of variance explained by the first principal components derived from RNA-seq data across PDX samples. The first four principal components capture the majority of variance in the dataset.

Bottom panels: Principal component analysis (PCA) plots of individual samples projected onto PC1 vs PC2 (left) and PC2 vs PC3 (right). Samples are labelled by PDX identifier. Replicate samples processed in different sequencing batches (PDX #585 and PDX #495) cluster closely together without clear separation by batch, indicating minimal batch effect arising from library preparation.

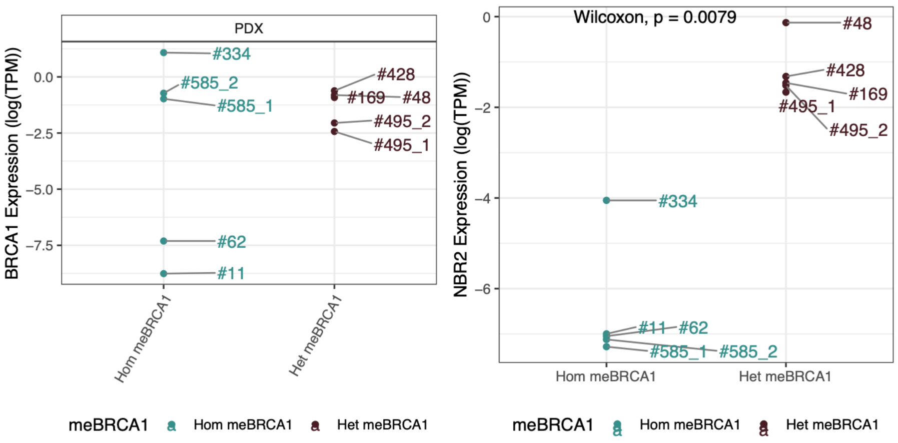


Figure 2. *BRCA1* and *NBR2* expression in relation to *BRCA1* promoter methylation status in PDX models.

*BRCA1* (left) and *NBR2* (right) expression (logTPM) across PDX models stratified by promoter methylation status (homozygous vs heterozygous me*BRCA1*). Higher inter-sample difference found compared to intrasample difference induced between batches for #585(#585_1 and #585_2) or #495 (#495_1 and #495_2).

#### 2. Data quality metrics confirm suitability for downstream analysis

We conducted whole-genome Nanopore sequencing on HGSOC models with varying me*BRCA1* levels. Before proceeding with the analysis, we assessed data quality to ensure that the sequencing reads were sufficient in quantity and length for profiling *BRCA1* at the allelic level, given its approximate size of 10 kb. The median read length ranged from 4 kb to 10kb (Figure 3A). The average sequencing depth across autosomal chromosomes exceeded 25x for all samples (26x – 48x; Figure 3B), ensuring adequate coverage for downstream genomic analysis. Additionally, the median of the mean base quality score per read was above 20 across all samples (median: 22.5 – 25), indicating a probability of an incorrect base call of less than 1 in 100 (Figure 3C).

Nanopore native DNA sequencing allows for the simultaneous detection of both DNA sequence and base-level modifications within a single read. To evaluate its performance in modified base detection, I compared the methylation profiles of three cell-line samples obtained through Nanopore sequencing with those from the widely used methylation array (EPIC V2) (data from [1]). A key limitation of the EPIC array is its inability to distinguish between 5mC and 5hmC modifications, as it captures a combined methylation signal after bisulfite conversion. To account for this limitation, I compared the methylation levels from the EPIC array with the sum of 5mC% and 5hmC% from Nanopore sequencing. This comparison revealed a high level of concordance across all three samples (correlation coefficient 0.96 in all three samples; Figure 4), reinforcing the reliability of Nanopore sequencing for detecting modified bases and highlighting its advantage in distinguishing between different methylation states.

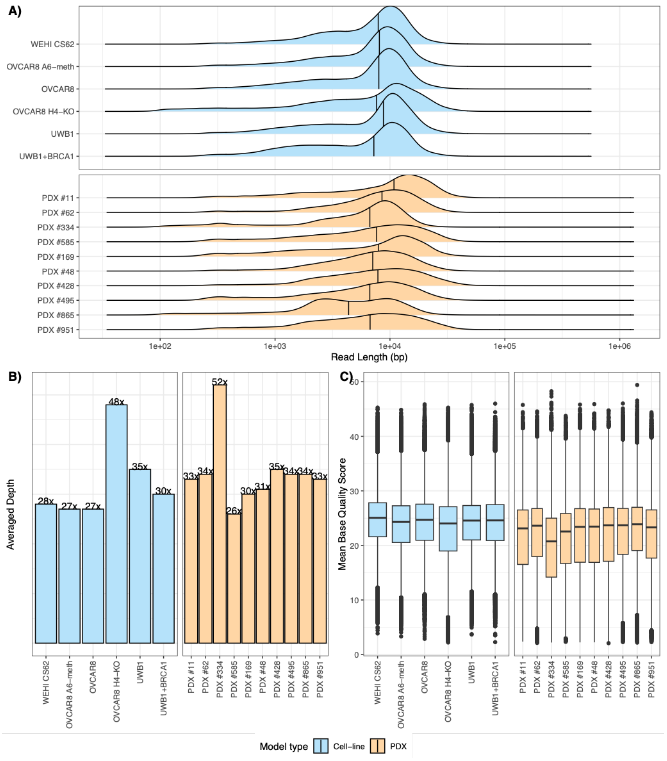


Figure 3: Quality assessment of autosomal reads by Nanopore sequencing for cell-lines and PDX models 1/100 of the reads were sub-sampled to illustrate the (A) read length distribution (vertical line indicates median read length), (B) averaged read depth covering autosomal chromosomes and (C) distribution of mean base quality scores per read.

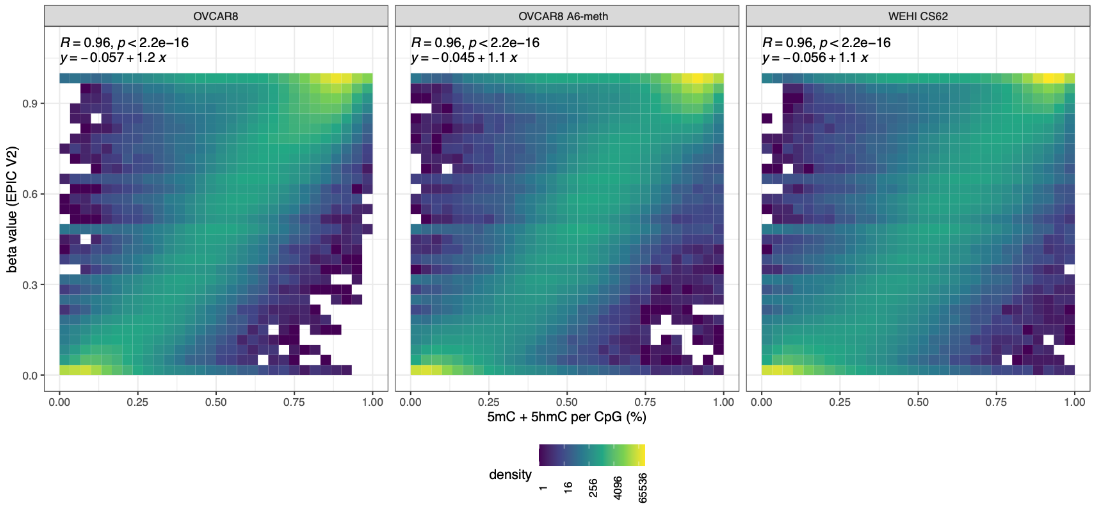


Figure 4: Base paired correlation between EPIC beta value and proportion of modified bases called by 5hmC+5mC basecalling model in OVCAR8, OVCAR8 A6-meth and WEHI-CS62. The x-axis represents the proportion of modified bases (5hmC + 5mC) called by the Nanopore basecalling model at each CpG, calculated as a percentage of total bases at the corresponding position. The y-axis represents the β -values from the Illumina EPIC methylation array, which approximate the proportion of methylated cytosines at the same positions. Colour density indicates the distribution of CpG sites with specific combinations of EPIC and Nanopore methylation levels: low density is shown in navy, while high density is shown in yellow.

#### 3. Consistent methylation status associated with sequence alteration in isogenic pairs

Prior to global methylation analysis, we first assessed whether methylation changes proximal to genomic alterations exhibited consistent directional trends—either increased or decreased methylation, rather than occurring in a random or stochastic manner. We analysed methylation correlation patterns within regions flanking sequence alterations in two engineered cell lines: OVCAR8 H4-KO ($BRCA1^{KO}$) and OVCAR8 A6-meth ($BRCA1^{me/-/-}$). These lines harbour both shared and unique indels and SVs. The shared variants likely arose prior to or during the insertion of the landing pad, while the unique variants most likely emerged following single-cell clone expansion after landing pad integration (Figure 3.2). Given the sequencing read length (median = 7 kb), the analysis was constrained to a 5 kb window upstream and downstream of each breakpoint to ensure adequate coverage and resolution of local methylation dynamics.

As the size of genomic alterations could theoretically have differing impacts on local methylation, we categorised the sequence alterations into large-scale variants (SVs; ≥ 50bp) and small indels (INDELs; <50bp). Focusing only on CpG sites that were either hypermethylated (≥0.8) or hypomethylated (≤0.2) in the parental line and the reference allele (REF) in the derivative line showed similar methylation levels (% methylation difference <0.2), we assessed the concordance of methylation in the altered allele (ALT) shared between H4-KO and A6-meth. Across most SV types, including small indels (INDEL-DEL and INDEL-INS; Figure 5A–B), large deletions (SV-DEL; Figure 5C), insertions (SV-INS; Figure 5D), and duplications (DUP; Figure 5E), methylation levels remained highly correlated between OVCAR8 H4-KO and OVCAR8 A6 -meth (R values ranging from 0.87 to 0.93). In contrast, sequences disrupted by translocations (BND) displayed lower methylation concordance (R value of

0.71; Figure 5F), suggesting that this type of sequence alteration may have more profound disruption on the methylation patterns with high variation in the resulting methylation pattern.

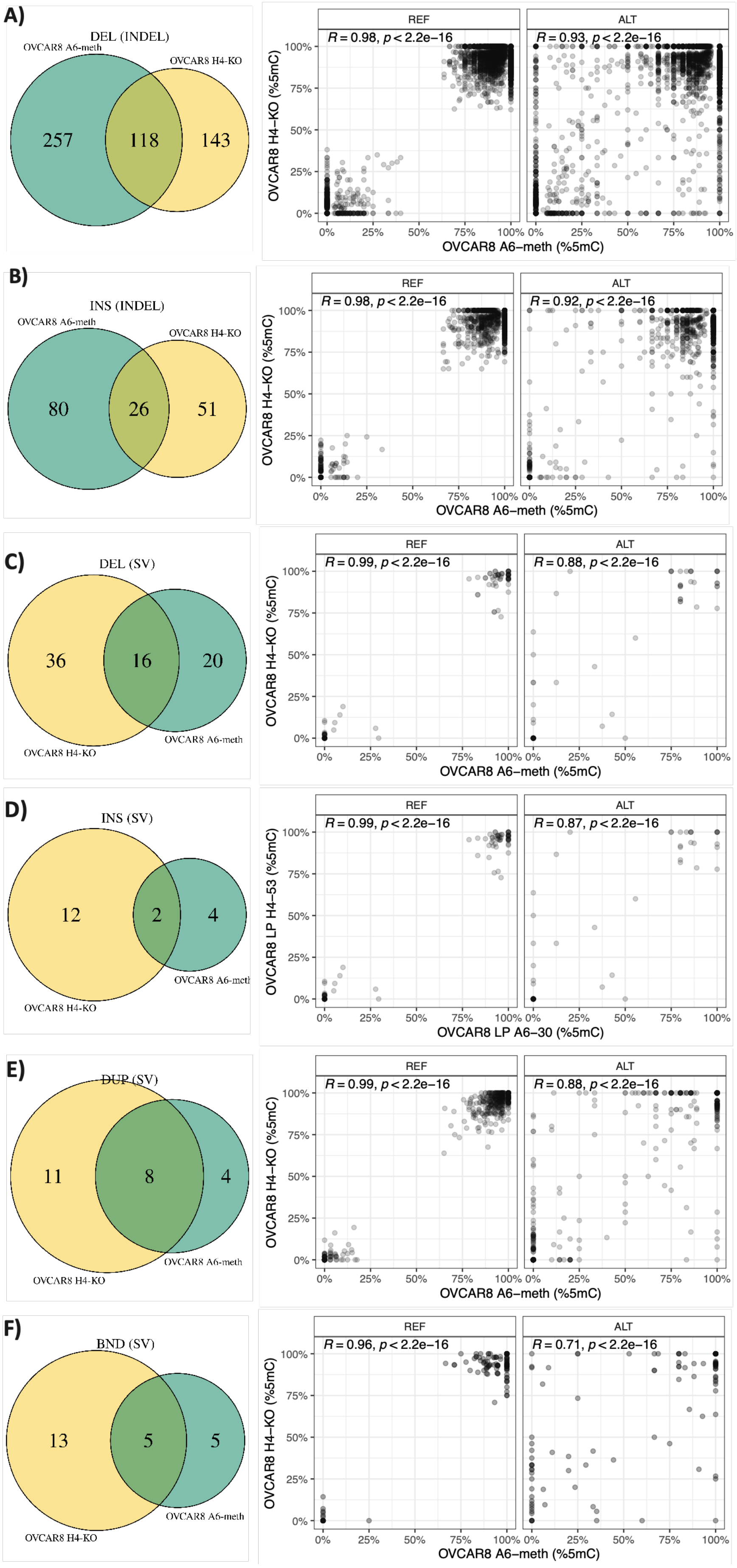
A)
DEL (INDEL)
OVCAR8 A6-meth
OVCAR8 H4-KO
257
118
143
REF
ALT
R = 0.98, p < 2.2e−16
R = 0.93, p < 2.2e−16
OVCAR8 H4−KO (%5mC)
OVCAR8 A6−meth (%5mC)
B)
INS (INDEL)
OVCAR8 A6-meth
OVCAR8 H4-KO
80
26
51
REF
ALT
R = 0.98, p < 2.2e−16
R = 0.92, p < 2.2e−16
OVCAR8 H4−KO (%5mC)
OVCAR8 A6−meth (%5mC)
C)
DEL (SV)
36
16
20
OVCAR8 H4-KO
OVCAR8 A6-meth
REF
ALT
R = 0.99, p < 2.2e−16
R = 0.88, p < 2.2e−16
OVCAR8 H4−KO (%5mC)
OVCAR8 A6−meth (%5mC)
D)
INS (SV)
12
2
4
OVCAR8 H4-KO
OVCAR8 A6-meth
REF
ALT
R = 0.99, p < 2.2e−16
R = 0.87, p < 2.2e−16
OVCAR8 LP H4−53 (%5mC)
OVCAR8 LP A6−30 (%5mC)
E)
DUP (SV)
11
8
4
OVCAR8 H4-KO
OVCAR8 A6-meth
REF
ALT
R = 0.99, p < 2.2e−16
R = 0.88, p < 2.2e−16
OVCAR8 H4−KO (%5mC)
OVCAR8 A6−meth (%5mC)
F)
BND (SV)
13
5
5
OVCAR8 H4-KO
OVCAR8 A6-meth
REF
ALT
R = 0.96, p < 2.2e−16
R = 0.71, p < 2.2e−16
OVCAR8 H4−KO (%5mC)
OVCAR8 A6−meth (%5mC)
0%
25%
50%
75%
100%

Figure 5: Correlation of CpG methylation levels in reads with or without shared sequence alterations between isogenic OVCAR8 derivatives (H4-KO and A6-meth.) CpGs that were either fully methylated (≥80%) or unmethylated (≤ 20%) in the parental OVCAR8 line and remained concordant in reference allele (REF) (methylation difference <20%) in both derivatives, were included. Methylation concordance was assessed across six categories: small (<50bp) (INDEL) deletions (DEL) (A) and (B) insertions (INS), (C) large (SV; ≥ 50bp) DEL and (D) INS (E) duplications (DUP), and (F) translocations (BND). For each variant class, the number of events detected in OVCAR8 H4-KO (yellow) and A6-meth (teal) is shown in the pie charts. The right panel displays methylation correlation at CpG sites located within the altered allele (ALT) shared by both cell lines.

### Reference


1. Nesic, K., et al., *Demethylating agents drive PARP inhibitor resistance in ovarian carcinomas with BRCA1 gene silencing*. bioRxiv, 2025: p. 2025.04.07.647418.
